# Detection of *Ramularia collo-cygni* from barley (*Hordeum vulgare*) in Australia using triplex quantitative and digital PCR

**DOI:** 10.1101/2021.07.04.451076

**Authors:** N. L. Knight, A. Moslemi, F. Begum, K. N. Dodhia, L. Covarelli, A. L. Hills, F. J. Lopez-Ruiz

## Abstract

Ramularia leaf spot (RLS), caused by *Ramularia collo-cygni*, is an emerging threat to barley (*Hordeum vulgare*) production. RLS has been reported in Australia; however only minimal information is available regarding its detection and distribution. Due to initial asymptomatic growth *in planta*, slow growth *in vitro* and symptomatic similarities to net blotch and physiological leaf spots, detection of this pathogen can be challenging. Quantitative PCR-based methods for *R. collo-cygni*-specific identification and detection have been described, however these assays (based upon the internal transcribed spacer [ITS] region) have been demonstrated to lack specificity. False-positive detections may have serious implications, thus we aimed to design a robust *R. collo-cygni*-specific PCR method. Using the phylogenetically informative RNA polymerase II second largest subunit (*rpb2*) and translation elongation factor 1-α (*tef1-α*) genes, along with the *tef1-α* gene of *H. vulgare*, a triplex assay was developed for both quantitative and digital PCR. The triplex assay was used to assess DNA of barley leaves from New South Wales, South Australia, Tasmania, Victoria and Western Australia, along with DNA of seeds from Western Australia. Detection of *R. collo-cygni* DNA was confirmed for leaf samples from New South Wales, South Australia, Tasmania, Victoria and Western Australia, indicating a distribution ranging across the southern barley growing regions of Australia. No *R. collo-cygni* DNA was detected in seed from Western Australia. The *R. collo-cygni*-specific assay will be a valuable tool to assist with monitoring the distribution of *R. collo-cygni* in Australia and other regions.

## Introduction

*Ramularia collo-cygni* infection of barley (*Hordeum vulgare*) has gained increasing attention over the past 30 years (Dussart et al. 2020; Havis et al. 2015b). The resulting disease, Ramularia leaf spot (RLS), has been reported in many barley growing regions of the world including Chile, Colombia, Europe, Mexico, New Zealand, South Africa and the United States (Beukes et al. 2016; Dussart et al. 2020; Havis and Brown 2019; Spencer et al. 2019; Walters et al. 2008). The first detections of *R. collo-cygni* in Australia were reported between 2010 and 2018 in New South Wales, Tasmania and Western Australia (Biosecurity Tasmania 2020; GRDC 2021; Oxley et al. 2010; Spencer et al. 2019); however, limited information regarding these detections is available.

*R. collo-cygni* typically grows asymptomatically during the early growing season, with RLS symptoms appearing late in the season, typically around anthesis and after head emergence (Kaczmarek et al. 2017; Walters et al. 2008). RLS is characterised by small necrotic lesions, usually with a chlorotic halo (Havis et al. 2015a; Sutton and Waller 1988). As the disease progresses these lesions coalesce, leading to large areas of the leaf being affected (Walters et al. 2008). Environmental conditions, including light intensity, leaf surface wetness, temperature and water availability, may affect RLS development (Havis et al. 2015b; Hoheneder et al. 2021b; Makepeace et al. 2008; McGrann and Brown 2018). Yield losses of up to 70% have been reported but are more commonly between 5 to 25%, with reductions in grain size and quality (Harvey 2002; Havis et al. 2015b; Pereyra et al. 2014; Pinnschmidt and Jørgensen 2009; Greif 2002 as cited in Sghyer and Hess 2019).

Seed-borne *R. collo-cygni* has been described as the main source of inoculum and as the primary mechanism for spread of the pathogen (Harvey 2002; Havis et al. 2014; Matusinsky et al. 2011). Alternative hosts, such as wheat and perennial grasses (Kaczmarek et al. 2017), and colonised plant debris may also play a role in harbouring inoculum. Dussart et al. (2020) provide a thorough review of the current understanding of *R. collo-cygni* epidemiology. RLS can be managed using *R. collo-cygni* free seeds, fungicides, resistant varieties, crop rotation and stubble reduction (Dussart et al. 2020; Hoheneder et al. 2021a; Oxley et al. 2010). However, the emergence of resistance to quinone-outside inhibitors (QoI) (Fountaine and Fraaije 2009; Matusinsky et al. 2010) and reduced sensitivity to succinate dehydrogenase inhibitors (SDHI) and demethylation inhibitors (DMI) (FRAC 2015; Rehfus et al. 2019) has intensified the need for varietal resistance and alternative control measures.

Effective management of RLS relies on early and accurate detection of the pathogen. Detection, however, can be challenging due to asymptomatic growth *in planta* (Havis et al. 2014; Kaczmarek et al. 2017), a lack of visible symptoms on seed (Oxley et al. 2010) and slow growth in culture (Walters et al. 2008). Visual identification is also problematic as RLS symptoms may appear similar to symptoms caused by *Pyrenophora teres* f. *maculata* (spot form net blotch), *P. teres* f. *teres* (net form net blotch) (Sachs et al. 1998 as cited in Walters et al. 2008) or physiological leaf spotting (Wu and Tiedemann 2002). DNA detection can be a reliable method for identifying the presence of pathogens, and several *R. collo-cygni*-specific PCR assays, based on sequences of the internal transcribed spacer (ITS) region (Frei et al. 2007; Havis et al. 2006; Matusinsky et al. 2011; Taylor et al. 2010), have been reported. These assays have been used to detect and quantify *R. collo-cygni* DNA in host tissue, including leaves and seeds (Havis et al. 2014).

PCR detection of a specific pathogen is reliant upon unique DNA sequences being associated with the pathogen of interest. However, as sequences of a greater number of fungal species, including *Ramularia* species (Videira et al. 2016), become available, the specificity of PCR assays requires confirmation. This should include critical assessment across different environments with potentially undescribed microflora. The specificity of *R. collo-cygni* PCR assays has been investigated using a range of plant pathogen DNA templates and genetic databases (Frei et al. 2007; Havis et al. 2006; Matusinsky et al. 2011; Taylor et al. 2010), however the utilisation of sensitive techniques such as quantitative PCR (qPCR) may increase the risk of false negative detection in the presence of similar sequences. Indeed, weak amplification of *R. indica* and *R. vallisumbrosae* was reported by Taylor et al (2010) using a qPCR assay. Confidence in detection, especially in new geographic regions, requires reassessment of previously described assays in light of new sequence data, along with development of alternative species-specific assays.

A range of methods for PCR detection of target DNA are available, including conventional PCR, real-time qPCR and digital PCR (dPCR). Quantitative PCR provides sensitive detection of low quantities of target DNA (approximately 1 pg *R. collo-cygni* DNA; Taylor et al. 2010) and is effective for plant pathogen detection (Schaad and Frederick 2002). Digital PCR is an emerging technology which uses assay design similar to qPCR and provides similarly sensitive detection of target DNA (Jones et al. 2016). The key difference is the need for a standard curve to quantify target DNA in qPCR, while dPCR relies on Poisson Distribution Analysis to assess end-point PCR fluorescence across partitioned droplets of reaction mixture (The dMIQE Group and Huggett 2020). Both qPCR and dPCR allow for simultaneous detection of multiple DNA targets (Klein 2002; Zhong et al. 2011). Such multiplexing can allow detection of pathogen and host DNA within the same sample, confirming DNA template quality when pathogen DNA is absent. This can be especially valuable when evaluating seed samples which may contain PCR inhibitors (Knight et al. 2020).

The spread of *R. collo-cygni* into new regions, such as Australia, has important management implications, especially considering the unknown nature of cultivar resistance or fungicide sensitivity. The first aim of this study was to provide improved confidence in *R. collo-cygni* identification and detection by developing alternative species-specific assays which can be adapted for use in both qPCR and dPCR platforms. The second aim was to assess leaf and seed samples from Australia for the presence of *R. collo-cygni* DNA.

## Materials and Methods

### DNA for assessing PCR specificity

DNA of *R. collo-cygni* (isolate Rcc_Pg_1), *R. endophylla* (isolate CBS 113265) and *R. pusilla* (isolate CBS 124973) was provided by the University of Perugia, Italy. DNA was extracted from mycelia harvested from 4-week-old cultures grown on potato dextrose agar (PDA) using the CTAB extraction method described by Covarelli et al. (2015). Rcc_Pg_1 was identified as *R. collo-cygni* based on morphology and partial sequence of the internal transcribed spacer (ITS) region amplified using RC3 and RC5 primers designed by Frei et al. (2007). *R. endophylla* (isolate CBS 113265) and *R. pusilla* (isolate CBS 124973) were obtained from the collection at the Westerdijk Fungal Biodiversity Institute. Dry DNA was sent to the Centre for Crop and Disease Management, Curtin University and re-suspended in 1% TE buffer (15.76 g Tris-Cl, 2.92 g EDTA, 1 L distilled water) and stored at −80 °C.

DNA of plant pathogenic fungi collected from Western Australian cereal growing regions was extracted for species-specificity assessment of the Ram6, *Rcc*_139_*rpb2, Rcc*_88_*tef1-α* and *Hv*_116_*tef1-α* PCR assays. These fungi included one single-spored isolate each of *Alternaria alternata*, *Ascochyta lentis*, *Blumeria graminis* f. sp. *hordei*, *Blumeria graminis* f. sp. *tritici*, *Botrytis cinerea*, *Curvularia trifolii*, *Diaporthe toxica*, *Parastagonospora nodorum*, *Pleiochaeta setosa, P. teres* f. *maculata* (isolate SG1; Mair et al. 2020), *P. teres* f. *teres* (isolate K0103; Mair et al. 2020), *Rhizoctonia* sp., *Septoria tritici* and *Stemphylium solani* provided by the Department of Primary Industries and Regional Development (DPIRD) and Centre for Crop and Disease Management, Curtin University. *B. graminis* f. sp. *hordei* and *B. graminis* f. sp. *tritici* were grown on detached barley and wheat leaves, respectively. Briefly, leaves of seven-day-old barley (cv. Baudin) and wheat (cv. Trojan) seedlings were detached from plants grown at room temperature with a 12-h photoperiod and placed into benzimidazole amended agar (10 g L^-1^ agar + 50 mg L^-1^ benzimidazole). Leaves were inoculated with spores and incubated under a 16-h photoperiod at 21 and 15 °C (light and dark, respectively) for seven days. The remaining isolates were grown on V8-PDA (Mair et al. 2016) and mycelia was harvested from seven-day-old cultures. Leaves or mycelia were frozen in liquid nitrogen and ground with two 9 mm diameter steel balls at 30 frequency for 60 s (MM400, Retsch). DNA was extracted using a BioSprint 15 DNA Plant Kit according to the manufacturer’s instructions (Qiagen, Australia) and stored at −20 °C. Pure barley DNA was extracted from leaves of seven-day-old barley seedlings (cv. Baudin) grown at room temperature with a 12-h photoperiod using the Biosprint method as described above. DNA concentrations were measured with a Qubit Flex Fluorometer (Thermo Fisher Scientific) using a Qubit dsDNA BR assay kit (Thermo Fisher Scientific) and adjusted to 2 ng μL^-1^.

### Specificity of published *R. collo-cygni* quantitative PCR assays

Specificity of the *R. collo-cygni* detection assay Ram6 described by Taylor et al. (2010) was investigated using a BLAST search of the GenBank nucleic acid database, aligning primer, probe and amplicon sequences against sequences of the *Ramularia* genus (taxid:112497) and by qPCR assessment of the fungal isolates described above. A similar alignment assessment was performed for the assay described by Matusinsky et al. (2011), but no PCR was included.

Quantitative PCR using the Ram6 assay was conducted following methods modified from Taylor et al. (2010). Reactions were performed in a CFX96 or CFX384 real-time system (Bio-Rad). Each 20 μL reaction consisted of 10 μL of SensiFAST Probe No-ROX Mix (2×) (Bioline), 0.25 μM each of RamF6 and RamR6 primers (Sigma, Australia), 0.15 μM of molecular beacon Ram6 (Sigma, Australia) and 5 μL of template DNA. Thermal cycling conditions consisted of 5 min at 95 °C, followed by 40 cycles of 95 °C for 10 s and 55 °C for 40 s. Fluorescence emission was recorded at the 55 °C step of each cycle. Each sample template was assessed in duplicate. DNA of *R. collo-cygni* (isolate Rcc_Pg_1) was included as a positive control, DNA of barley (cv. Baudin) was included as a negative control and nuclease free water was included as a no template control (NTC). Mixtures of *R. collo-cygni* and barley DNA were also assessed. Six ten-fold serial dilutions of *R. collo-cygni* DNA (ranging from 10 ng to 0.01 pg) were included for sensitivity analysis. A standard curve was generated using the CFX Maestro Software v. 1.1 (Bio-Rad) by plotting the logarithm of *R. collo-cygni* DNA concentrations against the quantification cycle (Cq). The coefficient of determination (*R^2^*), slope, y-intercept and reaction efficiency (E) were reported by the software for each standard curve.

### Development of species-specific PCR assays

Two *R. collo-cygni* specific assays (Table 1) were developed from sequences of the phylogenetically informative genes RNA polymerase II second largest subunit (*rpb2*) and translation elongation factor 1-alpha (*tef1-α*) (Videira et al. 2016). Sequences of *rpb2* and *tef1-α* from seven *R. collo-cygni* isolates and 181 isolates of other *Ramularia* species and related genera (Supp. Table S1; Videira et al. 2016), were imported from the National Center for Biotechnology Information (NCBI) GenBank database into Geneious v. 6.1.8 (https://www.geneious.com) and aligned using Geneious Alignment with default settings. DNA sequences unique to *R. collo-cygni* were manually identified and primers and probes were designed against these unique sequences and assessed using Primer3 v. 0.4.0 (Koressaar and Remm 2007; Untergasser et al. 2012) and PCR Primer Stats analysis in Sequence Manipulation Suite v. 2 (Stothard 2000). Primers were designed to include discriminatory polymorphic regions at the 3’ terminus (Petruska et al. 1988) and encompass unique sequences to which probes were aligned. Primer and probe sequences (Table 1) were assessed for specificity using a Basic Local Alignment Search Tool (BLAST) search of the Standard databases: Nucleotide Collection (nr/nt) in GenBank. Differences in the expect value (Lobo 2008) between *R. collo-cygni* and the next closest species hit in the BLAST search were used for assessing the likely specificity of the primer and probe sequences.

**Table 1.**
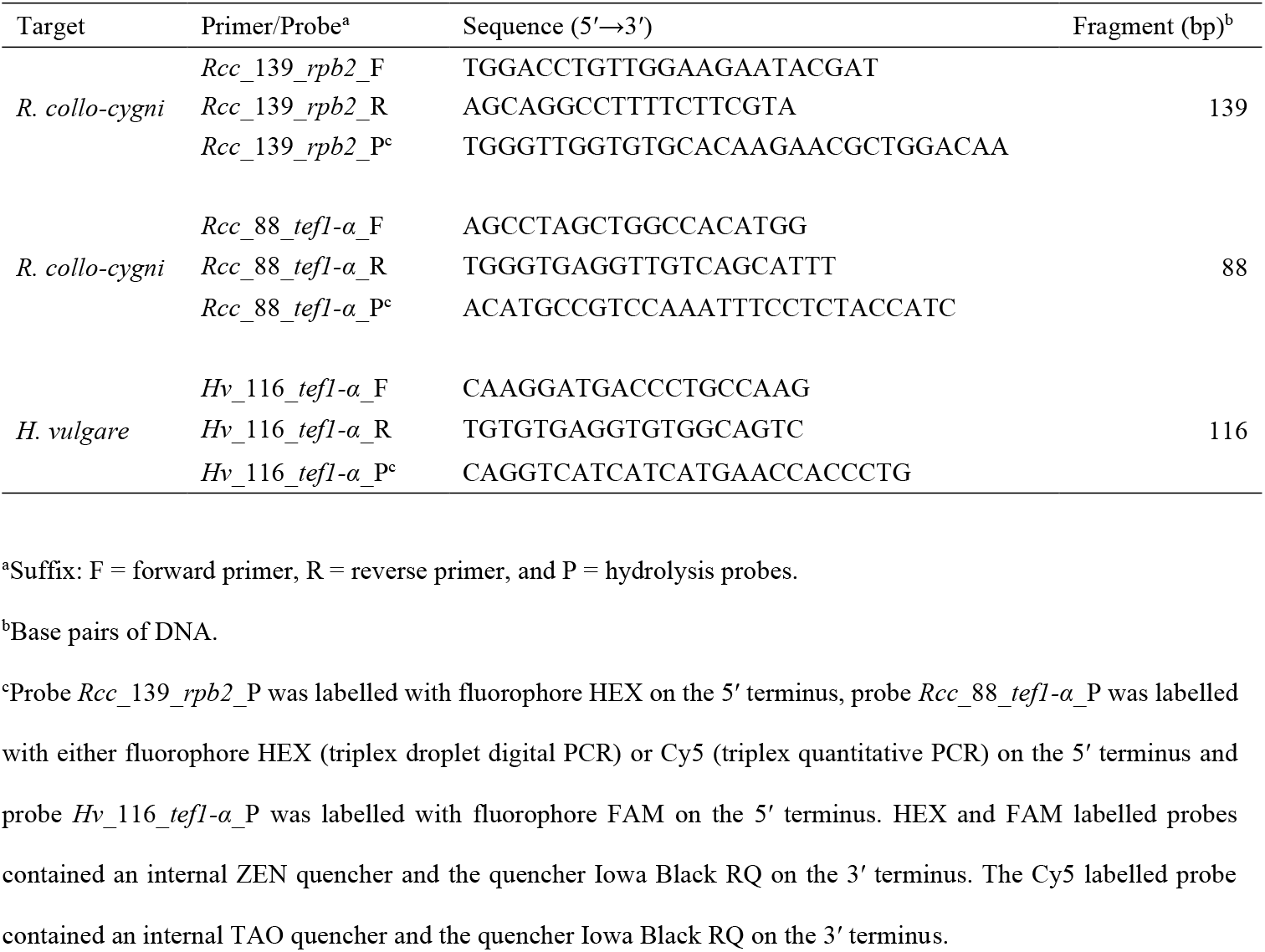
Primer/probe sets for detection and quantification of *Ramularia collo-cygni* and *Hordeum vulgare* DNA.

A *H. vulgare* specific assay (Table 1) was developed from sequences of the *tef1-α* gene. This assay acted as an internal positive control during PCR assessment of plant DNA templates. The design process for *H. vulgare* specific primers and probe was similar to that described above for the *R. collo-cygni* assays. Briefly, four *tef1-α* sequences of *H. vulgare* (Supp. Table S2) were identified in the NCBI sequence database and aligned in Geneious. Primers and probe were designed against the conserved regions and assessed for specificity by a BLAST search of the GenBank nucleic acid database.

### Quantitative PCR assessment of alternative *R. collo-cygni* specific assays

Uniplex qPCR reactions were assessed for assays *Rcc*_139_*rpb2*, *Rcc*_88_*tef1-α* and *Hv*_116 *tef1-α* (Table 1). Each 20 μL reaction consisted of 10 μL of SensiFAST Probe No-ROX Mix (2×) (Bioline, Australia), 0.25 μM of each forward and reverse primer (Integrated DNA Technologies, USA), 0.15 μM of probe and 5 μL of template DNA. Thermal cycling conditions were as described above, except for annealing temperatures of 66 °C for *Rcc_139_rpb2*, 64 °C for *Rcc_88_tef1-α* and 64 or 66 °C for *Hv*_116_*tef1-α*.

Simultaneous detection of multiple DNA targets was also assessed. In duplex qPCR, the *Hv*_116_*tef1-α* assay was combined with either *Rcc_139_rpb2* or *Rcc_88_tef1-α* assays. For triplex qPCR, all three assays were combined. Each 20 μL reaction was as described above, with the same individual primer and probe concentrations for duplex and triplex assays. Cycling conditions were as described above, with the annealing temperature adjusted to 64 °C for duplex and 60 °C for triplex assays. The DNA controls and standards described above were assessed in duplicate with each uniplex, duplex and triplex assay. The serial dilutions of *R. collo-cygni* DNA were also assessed in triplicate for the triplex assay and a trend line was fitted in Microsoft Excel 2016. The collection of DNA of fungal isolates was assessed with the duplex and triplex assays. For the triplex assay, intra- and inter-assay variation were reported as the mean DNA quantity and coefficient of variation of replicate samples of a mixed *R. collo-cygni* and *H. vulgare* DNA template.

### Droplet digital PCR assessment of alternative *R. collo-cygni* specific assays

The *Rcc*_88_*tef1-α, Rcc*_139_*rpb2* and *Hv*_116_*tef1-α* assays were further assessed in droplet digital PCR (ddPCR) in uniplex, duplex and triplex as described above. Droplet digital PCR was performed on a QX200 system (Bio-Rad), following the manufacturer’s instructions. Briefly, each 22 μL reaction consisted of 11 μL of 2× Bio-Rad ddPCR Supermix for Probes (no dUTP), 0.25 μM of each forward and reverse primer, 0.15 μM of probe and 5 μL of template DNA. Primer and probe concentrations were the same for uniplex and duplex assessment. Triplex assessment utilised amplitude-based multiplexing, with the *Hv*_116_*tef1-α* assay (FAM fluorophore) and *Rcc*_139_*rpb2* assay (HEX fluorophore) at the primer and probe concentrations described above, and the *Rcc*_88_*tef1-α* assay (HEX fluorophore) consisting of 0.9 μM of each forward and reverse primer and 0.25 μM of probe.

Thermal cycling conditions consisted of 95 °C for 10 min followed by 50 cycles of 94 °C for 30 s and 60 °C for 60 s, with a final denaturation at 98 °C for 10 min, followed by a hold step at 4 °C. The ramp rate between temperatures was 2 °C s^-1^. Results were retrieved from the QX Manager Standard Edition software v. 1.2 (Bio-Rad). The DNA controls and standards described above were assessed in duplicate with each uniplex, duplex and triplex assay. The serial dilutions of *R. collo-cygni* DNA were also assessed in triplicate for the triplex assay and a trend line was fitted in Microsoft Excel 2016. For the triplex assay, intra- and inter-assay variation were reported as the mean copy number μL^-1^ and coefficient of variation of replicate samples of a mixed *R. collo-cygni* and *H. vulgare* DNA template.

To be considered positive, positions of droplets had to align with droplets of standard DNA samples. For low copy numbers, a minimum requirement for positive detection was for droplets with each single DNA target to be present and to be above the limit of detection. Samples not meeting these requirements were re-tested. When the *Hv*_116_*tef1-α* assay or the *Rcc*_139_*rpb2* and *Rcc*_88_*tef1-α* assays reported more than 10 000 or 5000 copies μL^-1^, respectively, a ten-fold dilution of the DNA template was assessed.

### Suspected Ramularia leaf spot affected leaves

Barley leaves exhibiting symptoms resembling Ramularia leaf spot were collected from New South Wales, South Australia, Tasmania, Victoria, and Western Australia (Table 2). Cultivars sampled from two locations in Tasmania in 2016 included GrangeR, RGT Planet and Westminster. Samples of cultivar Baudin were collected from a single field in NSW in 2016. Samples of cultivar RGT Planet were collected from single fields in South Australia, Tasmania and Victoria in 2020. Samples of cultivars Oxford and Rosalind were collected from fields in Western Australia in 2018 and 2020, respectively.

**Table 2.** Detection and quantification of *Ramularia collo-cygni* DNA in barley leaf samples from New South Wales (NSW), South Australia (SA), Tasmania (Tas), Victoria (Vic) and Western Australia (WA) using the *R. collo-cygni*-specific triplex and Ram6 quantitative PCR assays.

| Region | State | Year | n<sup>a</sup> | *Rcc_139_rpb2*<sup>c</sup> |  |  | *Rcc_88_tefI-α*<sup>d</sup> |  | Ram6<sup>f</sup> |  |  |
| --- | --- | --- | --- | --- | --- | --- | --- | --- | --- | --- | --- |
|  |  |  |  | Incidence (%)<sup>b</sup> | Cq | DNA (pg) | Cq | DNA (pg) | Incidence (%)<sup>e</sup> | Cq | DNA (pg) |
| Mulwala | NSW | 2016 | 6 | 17 | 30.8<sup>g</sup> | 8 | 31.6 | 5 | 50 | 31.7 ± 1.92 | 1 ± 0.95 |
| Dairy Plains | TAS | 2016 | 9 | 100 | 27.1 ± 1.01 | 244 ± 112.55 | 27.4 ± 1.05 | 271 ± 119.87 | 100 | 23.3 ± 1.06 | 159 ± 68.7 |
| Hagley | TAS | 2016 | 10 | 70 | 28.7 ± 1.4 | 133 ± 64.44 | 29.2 ± 1.32 | 121 ± 58.79 | 90 | 25.7 ± 1.36 | 96 ± 65.03 |
| South Stirling | WA | 2018 | 60 | 0 | —<sup>h</sup> | — | — | — | 3 | 31.8 ± 2.47 | 3 ± 2.64 |
| Conmurra | SA | 2020 | 10 | 100 | 26.6 ± 0.22 | 185 ± 17.21 | 28 ± 0.62 | 101 ± 11.11 | 100 | 23.7 ± 0.64 | 75 ± 30.46 |
| Hagley | TAS | 2020 | 10 | 100 | 21.1 ± 0.77 | 1385 ± 206.6 | 24.5 ± 0.69 | 1459 ± 249.88 | 100 | 19.7 ± 0.75 | 1267 ± 619.29 |
| Gnarwarre<sup>i</sup> | VIC | 2020 | 10 | 100 | 22.9 ± 0.06 | 1266 ± 36.92 | 23.5 ± 0.18 | 1337 ± 47.52 | 100 | 18.6 ± 0.18 | 1185 ± 111.54 |
| South Stirling<sup>i</sup> | WA | 2020 | 80 | 1 | 33.6 | 5 | 34.4 | 2 | 14 | 33.8 | 1 |

<sup>a</sup> Number of leaf samples.
<sup>b</sup> Percentage of leaf samples for which both *Rcc\_139\_rpb2* and *Rcc\_88\_tef1-α* were detected prior to a quantification cycle (Cq) of 35.
<sup>c</sup> Mean ± standard error of the Cq and the calculated DNA quantity for *Rcc\_139\_rpb2*. Samples with no detection for either *Rcc\_139\_rpb2* or *Rcc\_88\_tef1-α* were not included in the calculation.
<sup>d</sup> Mean ± standard error of the Cq and the calculated DNA quantity for *Rcc\_88\_tef1-α*. Samples with no detection for either *Rcc\_139\_rpb2* or *Rcc\_88\_tef1-α* were not included in the calculation.
<sup>e</sup> Percentage of leaf samples for which Ram6 (Taylor et al. 2010) was detected prior to a Cq of 35.
<sup>f</sup> Mean ± standard error of the Cq and the calculated DNA quantity for Ram6. Samples with no detection were not included in the calculation.
<sup>g</sup> For values with no standard error value, *Rcc\_139\_rpb2* and *Rcc\_88\_tef1-α* or Ram6 were detected in one leaf sample.
<sup>h</sup> A dash (—) indicates the absence of a Cq.
<sup>i</sup> Samples were diluted ten-fold prior to assessment in triplex quantitative PCR based on initial failure to amplify target DNA of the *Hv\_116\_tef1-α* assay. Values represent the ten-fold dilutions.

For each leaf, 5 cm of tissue was removed, cut into pieces and ground with two steel balls at 30 frequency for 2× 60 s (MM400, Retsch). DNA was extracted using the BioSprint 15 protocol described above. Leaf DNA samples were assessed in qPCR and ddPCR using the triplex assay designed in this study and in qPCR using the Ram6 assay described by Taylor et al. (2010). Reaction conditions and standard templates were as described above. When amplification of the *Hv*_116_*tef1-α* assay failed, a ten-fold dilution of the DNA template was assessed.

### PCR assessment of barley grain samples

Ninety-five barley grain samples (200 g), originating from the southern Western Australian grain growing region, were provided by DPIRD from seed lots produced in 2019 and 2020. Samples were supplied based on a suspected association with Ramularia leaf spot symptoms in the field. Four replicate 15 g sub-samples of each grain sample were processed separately in a blender (Breville, Australia) to produce a fine powder. One gram of ground material from each replicate was removed and mixed together. DNA was extracted from each 4 g sample using a modified CTAB DNA extraction method described by Beccari et al. (2019). DNA concentrations were determined as described above and samples were assessed in PCR at the extracted concentration or as a ten-fold dilution if detection the *Hv*_116_*tef1-α* assay failed. Grain DNA samples were assessed in qPCR and ddPCR using the triplex assay designed in this study and in qPCR using the Ram6 assay described by Taylor et al. (2010). Reaction conditions and standard templates were as described above.

## Results

### Characteristics of published *R. collo-cygni* quantitative PCR assays

A BLAST search of the RamF6 and RamR6 primers and Ram6 molecular beacon (Taylor et al. 2010) based on ITS region sequences indicated 100% sequence identity with *R. collo-cygni*. Individual primer and probe sequences also shared 100% sequence identity with a range of other *Ramularia* and fungal species. In particular, each primer and probe sequence was 100% identical to *R. grevilleana* and *R. pusilla*. Nine other *Ramularia* species were identified with highly similar sequence identities (Supp. Fig. S1). Similar assessment of the RCCjlF, RCCj3R and RCCSON primers and probe (Matusinsky et al. 2011) indicated 100% shared sequence identity with 16 *Ramularia* species (Supp. Fig. S2). No further assessment of this assay was performed.

Using the Ram6 assay *R. collo-cygni* DNA was quantifiable from 0.001 to 10 ng. The standard curve of the quantification cycle and log of standard DNA quantities was calculated for the Ram6 assay (*R^2^* > 0.99, y = − 3.9x + 20.3, E = 80.6). PCR assessment of the 2 ng μL^-1^ DNA of the fungal collection indicated positive detection of *R. collo-cygni* (Cq = 15.6), *R. endophylla* (Cq = 15.8) and *R. pusilla* (Cq = 22.6). Fluorescence was detected for DNA of the remaining fungal isolates between 34 and 37 cycles. A cut-off point was set at 35 cycles.

### Characteristics of alternative *R. collo-cygni* and *H. vulgare* quantitative PCR assays

Sequences unique to *R. collo-cygni* were identified from multiple species sequence alignments of the *rpb2* and *tef1-α* genes, and informed the primer and probe positions for assays *Rcc*_139_*rpb2* and *Rcc*_88_*tef1-α* (Fig. 1). A BLAST search for each primer and probe indicated 100% sequence identity with *R. collo-cygni*. No primers or probes were 100% similar to any other fungal or plant species in the Standard databases: Nucleotide Collection (nr/nt) in GenBank (Supp. Figs. S3 and S4).

**Fig. 1.**
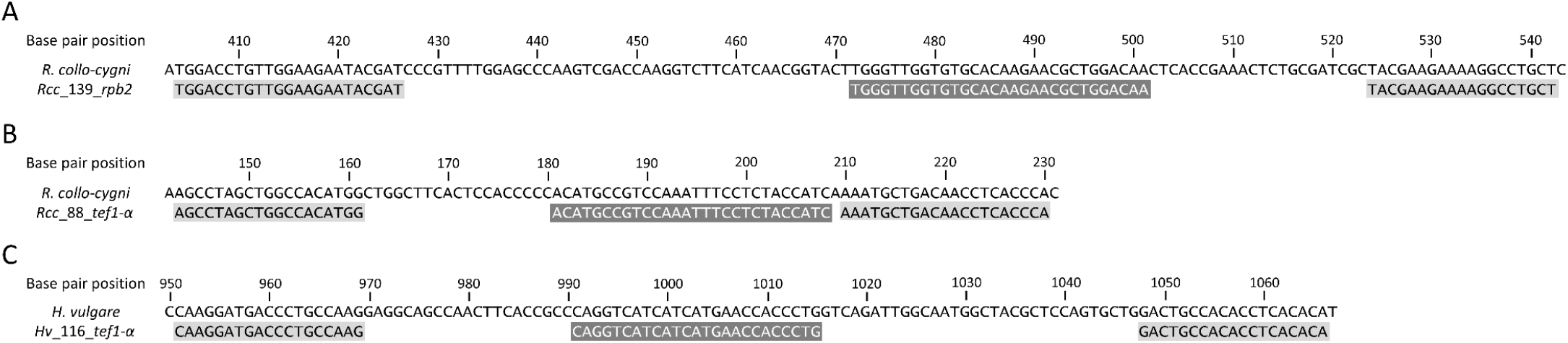
Alignment of primers (black text on grey) and probes (white text on grey) against the second largest subunit of RNA polymerase II (A) and translation elongation factor 1-α (B) gene sequences of *Ramularia collo-cygni*, and the translation elongation factor 1-α gene sequence of *Hordeum vulgare*. GenBank accession numbers KX288543 (A), KX287944 (B) and KP293845 (C) were used for base pair reference positions.

A BLAST search for the primers and probe designed for the *Hv*_116_*tef1-α* assay (Fig. 1) indicated 100% similarity with *H. vulgare* and a range of other plant species. Neither primer was 100% similar to any fungal species, while the probe was 100% similar to species of five fungal genera.

Quantification and standard curve characteristics were similar for duplex (data not shown) and triplex reactions. In triplex, *R. collo-cygni* DNA was quantifiable from 0.001 to 10 ng. The standard curve of the quantification cycle and log of standard DNA quantities was calculated for *Rcc*_139_*rpb2* (*R^2^* > 0.99, y = − 3.5x + 24.2, E = 94) and *Rcc*_88_*tef1-α* (*R^2^* = 0.99, y = − 3.4x + 24.4, E = 97) (Fig. 2 and Supp. Table S3).

**Fig. 2.**
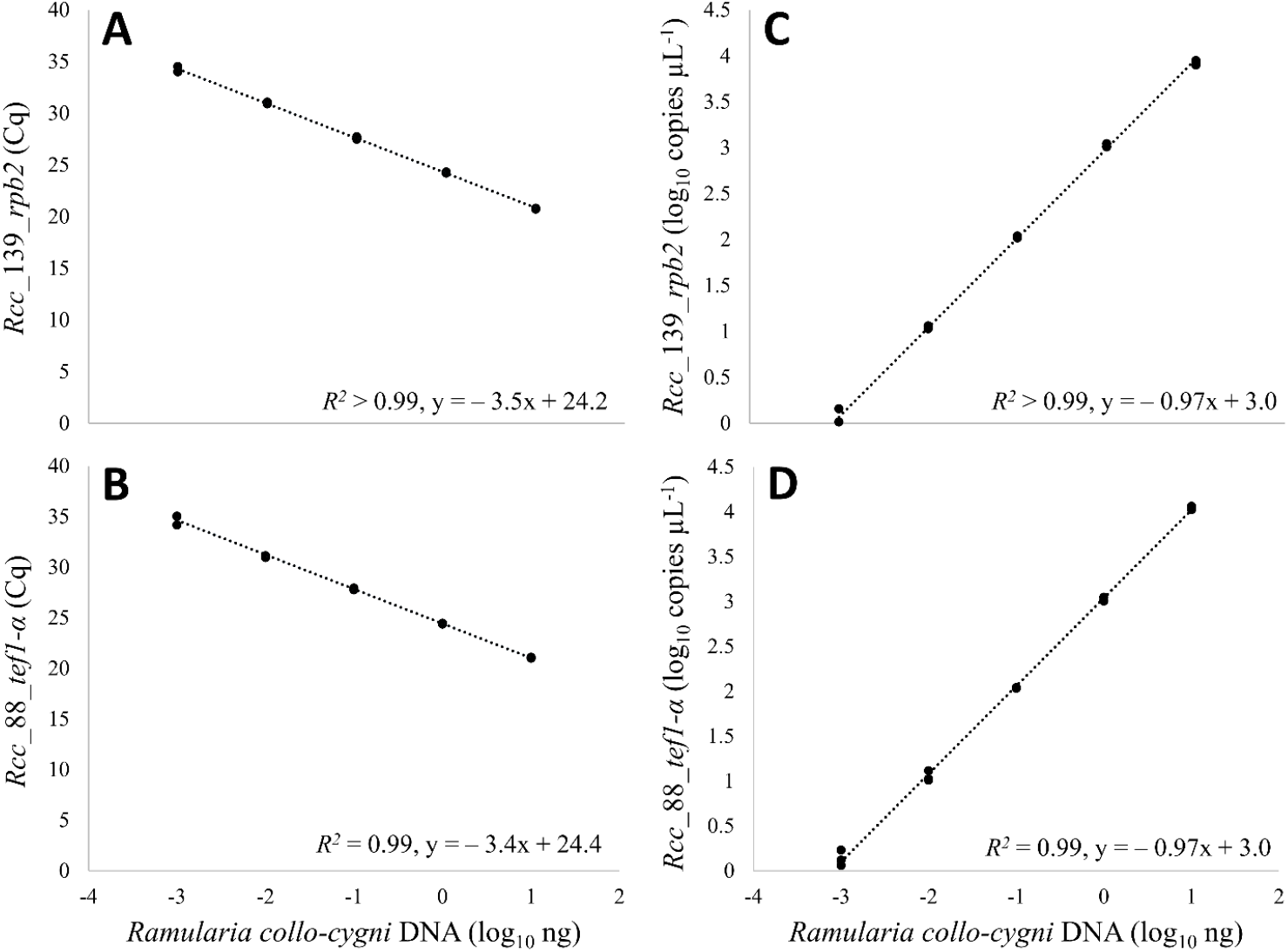
Linear relationship between the quantification cycle (Cq) and the logarithm of the *R. collo-cygni* DNA quantity for the *Rcc*_139_*rpb2* assay (A) and *Rcc*_88_*tef1-α* assay (B) in quantitative PCR, and the logarithm of the copies per microliter and the logarithm of the *R. collo-cygni* DNA quantity for the *Rcc*_139_*rpb2* assay (C) and *Rcc*_88_*tef1-α* assay (D) in droplet digital PCR. *R. collo-cygni* DNA quantities were a ten-fold serial dilution from 10 to 0.001 ng (*n* = 3). The correlation coefficient (*R^2^*) and linear equation (y= mx+c) are provided.

The respective mean values (*n* = 20) and coefficient of variation (CV) of a mixed DNA sample assessed for intra-assay variability were 50 pg *R. collo-cygni* DNA (CV = 8.1%) for *Rcc*_139_*rpb2*, 40 pg *R. collo-cygni* DNA (CV = 7.9%) for *Rcc*_88_*tef1-α* and 1320 pg *H. vulgare* DNA (CV = 8.5%) for *Hv*_116_*tef1-α*. The respective mean values (*n* = 9) and CV of the mixed DNA sample assessed for inter-assay variability were 70 pg *R. collo-cygni* DNA (CV = 59.0%) for *Rcc*_139_*rpb2*, 63 pg *R. collo-cygni* DNA (CV = 61.0%) for *Rcc*_88_*tef1-α* and 1080 pg *H. vulgare* DNA (CV = 43.0%) for *Hv*_116_*tef1-α*.

PCR assessment of the DNA of the fungal collection indicated positive detection of only *R. collo-cygni* DNA after 40 cycles in the *Rcc*_139_*rpb2* and *Hv*_116_*tef1-α* duplex. In the *Rcc*_88_*tef1-α* and *Hv*_116_*tef1-α* duplex only *R. collo-cygni* DNA was detected prior to cycle 38. Low levels of fluorescence were observed after cycle 38 for some samples, but were not consistent between replicates. In the triplex assay only *R. collo-cygni* DNA was detected prior to cycle 38, with late fluorescence inconsistently detected among some samples for both *R. collo-cygni* assays. The *Hv*_116_*tef1-α* assay in each duplex or triplex only detected *H. vulgare* DNA (cv. Baudin).

Based on these results a cut-off point, defined as the cycle number above which any sample response value (quantification cycle) was considered a false positive due to non-specific fluorescence, was set at 35 cycles. Any fluorescence detection after cycle 35 (outside of the detected standard DNA sample concentrations) was not considered a positive detection of *R. collo-cygni* DNA. Positive detection also required fluorescence to be reported for both the *Rcc*_139_*rpb2* and *Rcc*_88_*tef1-α* assays.

### Droplet digital PCR of alternative *R. collo-cygni* and *H. vulgare* assays

Amplitude-based multiplexing allowed clear separation of the individual and combined products of the *Rcc*_139_*rpb2* and *Rcc*_88_*tef1-α* assays, and the *Hv*_116_*tef1-α* assay (Fig. 3). In triplex ddPCR, *R. collo-cygni* DNA was detectable from 0.001 to 10 ng (1 to 10 000 copies μL^-1^, respectively). Uniplex and duplex assays performed similarly (data not shown). The respective mean values (*n* = 20) and coefficient of variation (CV) of a single mixed DNA sample assessed for intra-assay variability were 134 copies μL^-1^ (CV = 3.5%) for *Rcc*_139_*rpb2*, 135 copies μL^-1^ (CV = 2.9%) for *Rcc*_88_*tef1-α* and 85 copies μL^-1^ (CV = 4.1%) for *Hv*_116_*tef1-α*. The respective mean values (*n* = 10) and CV of a single mixed DNA sample assessed for inter-assay variability were 105 copies μL^-1^ (CV = 12.1%) for *Rcc*_139_*rpb2*, 112 copies μL^-1^ (CV = 11.0%) for *Rcc*_88_*tef1-α* and 69 copies μL^-1^ (CV = 12.5%) for *Hv*_116_*tef1-α*. Copy numbers for the *Rcc*_139_*rpb2* and *Rcc*_88_*tef1-α* assays were consistently close to a 1:1 ratio in reactions with less than 10 000 or 5000 copies μL^-1^ reported by the *Hv*_116_*tef1-α* assay or the *Rcc*_139_*rpb2* and *Rcc*_88_*tef1-α* assays, respectively.

**Fig. 3.**
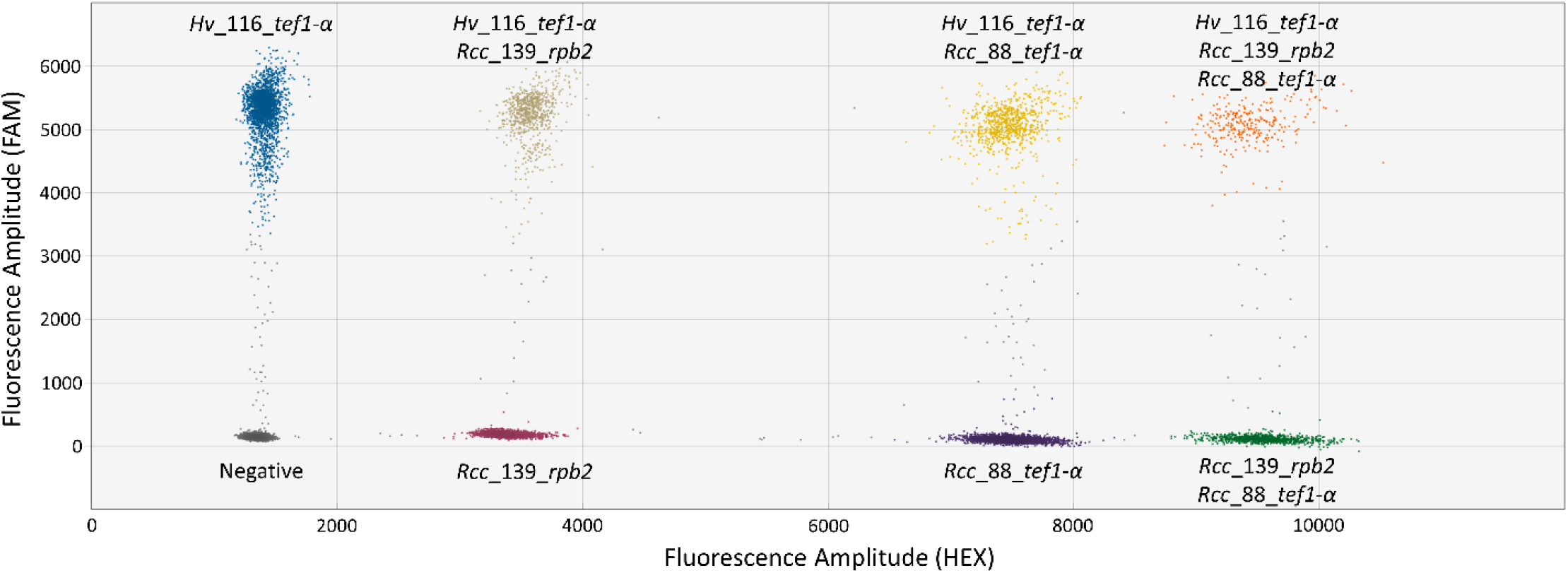
Representative 2D plot of droplet fluorescence during amplitude-based triplex droplet digital PCR combining the *Hv*_116_*tefl-α* (*Hordeum vulgare*; FAM fluorophore), *Rcc*_139_*rpb2* (*Ramularia collo-cygni*; HEX fluorophore) and *Rcc*_88_*tef1-α* (*R. collo-cygni*; HEX fluorophore) assays. The sample template was a ten-fold dilution of DNA extracted from a diseased leaf. Eight droplet clusters are present, indicating the amplification of each DNA target as either single or combined detections within each droplet. The negative cluster indicates droplets which did not contain DNA targets. Copy numbers per microliter for the *Hv*_116_*tef1-α*, *Rcc*_139_*rpb2* and *Rcc*_88_*tefl-α* assays were 346, 362 and 375, respectively.

Rain (droplets emitting fluorescence between the negative and positive clusters) was observed more frequently at higher copy numbers, predominantly for the *Hv*_116_*tef1-α* assay. PCR of the 2 ng μL^-1^ DNA of the fungal collection reported variable values up to 0.6 copies μL^-1^. Detection was inconsistent across replicates for both *R. collo-cygni* assays.

Based on this information a cut-off point, defined as the copy number below which any sample value was considered a false positive, was set at 1 copy μL^-1^. Positive detection also required fluorescence across both the *Rcc*_139_*rpb2* and *Rcc*_88_*tef1-α* assays.

### Detection of *R. collo-cygni* DNA in leaf samples

*R. collo-cygni* DNA was detected in leaves from New South Wales, South Australia, Tasmania, Victoria and Western Australia (Tables 2 and 3). The incidence of detection was 100% in South Australia, Tasmania, and Victoria from samples collected in 2020. In Western Australia one leaf sample from 2020 was positive for *R. collo-cygni* DNA and no leaves from 2018 had detectable *R. collo-cygni* DNA. Samples collected in 2016 from two locations in Tasmania had incidences of detection of 60 to and 100%, while one leaf sample from New South Wales was positive for *R. collo-cygni* DNA. Triplex qPCR and ddPCR assays reported similar positive detections. In comparison, when the incidence of detection was less than 100%, the Ram6 assay consistently had greater incidence of detection values than the triplex assay (Table 2).

**Table 3.** Detection and quantification of *Ramularia collo-cygni* DNA in leaf samples from New South Wales (NSW), South Australia (SA), Tasmania (Tas), Victoria (Vic) and Western Australia (WA) using *R. collo-cygni*-specific triplex droplet digital PCR.

| Region | State | Year | *n*<sup>a</sup> | Incidence (%)<sup>b</sup> | *Rcc_139_rpb2* (copies $\mu\text{L}^{-1}$)<sup>c</sup> | *Rcc_88_tef1-a* (copies $\mu\text{L}^{-1}$)<sup>d</sup> |
| --- | --- | --- | --- | --- | --- | --- |
| Mulwala | NSW | 2016 | 6 | 17 | 1<sup>e</sup> | 1 |
| Dairy Plains<sup>f</sup> | TAS | 2016 | 9 | 89 | 131 ± 54.9 | 162 ± 59.8 |
| Hagley<sup>f</sup> | TAS | 2016 | 10 | 60 | 59 ± 26.2 | 74 ± 32.4 |
| South Stirling | WA | 2018 | 60 | 0 | —<sup>g</sup> | — |
| Conmurra<sup>f</sup> | SA | 2020 | 10 | 100 | 29 ± 12.1 | 32 ± 14.5 |
| Hagley<sup>f</sup> | TAS | 2020 | 10 | 100 | 648 ± 354.9 | 674 ± 371.5 |
| Gnarwarre<sup>f</sup> | VIC | 2020 | 10 | 100 | 4642 ± 588.3 | 4764 ± 612.3 |
| South Stirling | WA | 2020 | 80 | 1 | 2.3 | 4.1 |

<sup>a</sup> Number of leaf samples.
<sup>b</sup> Percentage of leaf samples for which both *Rcc\_139\_rpb2* and *Rcc\_88\_tef1-a* were detected at greater than 1 copy $\mu\text{L}^{-1}$ .
<sup>c</sup> Mean ± standard error of the number of copies per microliter for *Rcc\_139\_rpb2*. Samples with detection of less than 1 copy $\mu\text{L}^{-1}$ for either *Rcc\_139\_rpb2* or *Rcc\_88\_tef1-a* were not included in the calculation.
<sup>d</sup> Mean ± standard error of the number of copies per microliter for *Rcc\_88\_tef1-a*. Samples with detection of less than 1 copy $\mu\text{L}^{-1}$ for either *Rcc\_139\_rpb2* or *Rcc\_88\_tef1-a* were not included in the calculation.
<sup>e</sup> For values with no standard error value, *Rcc\_139\_rpb2* and *Rcc\_88\_tef1-a* were detected in one leaf sample.
<sup>f</sup> Samples were diluted ten-fold prior to assessment in droplet digital PCR. Sample copies per microliter values were multiplied by 10 before calculating the mean and standard error.
<sup>g</sup> A dash (—) indicates the detection of less than 1 copy $\mu\text{L}^{-1}$ for *Rcc\_139\_rpb2* and *Rcc\_88\_tef1-a*.

Leaf samples varied in age and condition, with completely senesced samples appearing to inhibit PCR reactions. For samples where the *Hv*_116_*tef1-α* assay failed to amplify barley DNA, a ten-fold dilution of the template enabled the detection of barley DNA, and the potential detection of *R. collo-cygni* DNA (Supp. Tables S4 and S5).

### Detection of *R. collo-cygni* DNA in seed samples

*R. collo-cygni* DNA was not detected in the 95 seed DNA samples using the triplex assay in qPCR and ddPCR (Table 4 and Supp. Tables S6 and S7). Barley DNA was detected for each sample at greater than 1 copy μL^-1^.

**Table 4.**
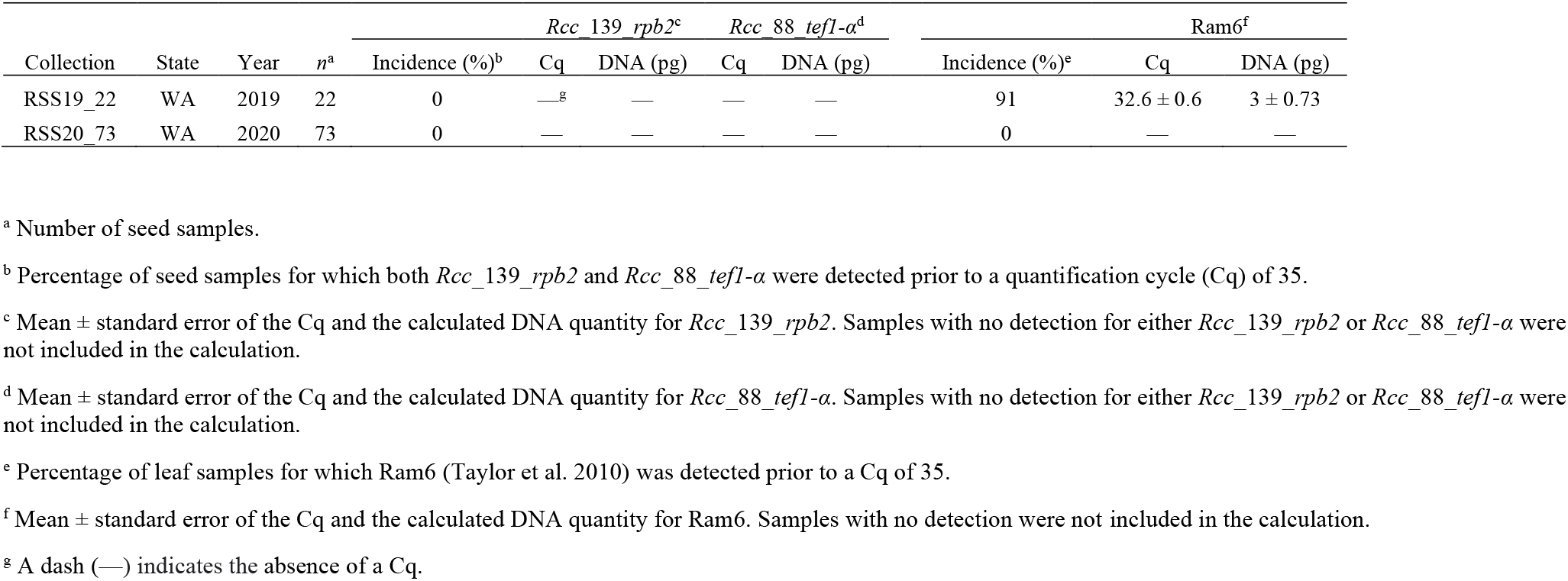
Detection and quantification of *Ramularia collo-cygni* DNA in barley seed sample collections from Western Australia (WA) using the *R. collo-cygni*-specific triplex and Ram6 quantitative PCR assays.

Quantitative PCR with the Ram6 assay reported an incidence of detection of 91% for the 22 DNA samples extracted from seed collected in 2019 (Table 4). No detection was reported for the 73 seed samples from 2020.

## Discussion

Specific detection of pathogens is critical for reporting new incursions and pathogen distribution, and for informing appropriate disease control measures. Methods for detection should be supported by the latest taxonomic information and critically assessed across different regions and environments. Previously reported *R. collo-cygni*-specific PCR assays (Matusinsky et al. 2011; Taylor et al. 2010) have been demonstrated to detect other DNA targets, including other *Ramularia* species, and may report false positive detections. To improve confidence in *R. collo-cygni* detection, a triplex assay was designed to simultaneously detect two *R. collo-cygni*-specific DNA sequences, along with barley DNA as a positive control in plant samples. This assay has been used to confirm the presence of *R. collo-cygni* DNA in barley tissues using both quantitative and droplet digital PCR, suggesting widespread distribution of *R. collo-cygni* across the southern barley growing regions of Australia.

Interest in designing alternative *R. collo-cygni*-specific PCR assays was initiated by reports of *R. collo-cygni* in Australia (Biosecurity Tasmania 2020; GRDC 2021; Oxley et al. 2010; Spencer et al. 2019). Primary investigations of barley tissues were performed using the Ram6 assay (Taylor et al. 2010), with positive detection of *R. collo-cygni* across regions with contrasting environments bringing the assay specificity under scrutiny. The presence of various *Ramularia* species in Australia (Australia’s Virtual Herbarium 2021; Braun et al. 2005; Plant Health Australia 2001), including the likelihood of undescribed species being present in the environment, suggested a risk for false positive detection. In the current study, DNA of *R. endophylla* and *R. pusilla* reported similar Cq values to *R. collo-cygni* DNA, with ITS region sequence alignments suggesting theoretical detection of several other *Ramularia* species. Potential false positive results due to *R. pusilla* detection are a risk, as this fungus has been described on several grass species in Australia, including *Lolium rigidum* (ryegrass) (Braun et al. 2005) which is a widespread weed in cereal fields (Lazarides et al. 1997). The greater frequency of detection for the Ram6 assay in leaf DNA and particularly in DNA from seed collected in 2019 indicates that false positive detection was occurring for field samples. More comprehensive sequencing and taxonomic assessment of the Ramularia genus has been performed since the first *R. collo-cygni*-specific qPCR assays were described (Videira et al. 2016), providing a resource to enable improved *R. collo-cygni*-specific PCR assays to be designed. Videira et al. (2016) reported that the ITS region sequence provided poor resolution within the Ramularia genus, while the actin (*actA*), glyceraldehyde 3-phosphate dehydrogenase (*gapdh*), *rpb2* and *tef1-α* genes provided a larger barcode gap and less overlap between intra- and inter-specific distances. Thus, the *rpb2* and *tef1-α* genes were selected for the design of *R. collo-cygni*-specific PCR assays.

The manual alignment of primers and probes to species-specific regions in the *rpb2* and *tef1-α* gene sequences and inclusion of a host DNA target followed previously reported methods for designing species-specific PCR assays (Knight et al. 2012; Knight and Pethybridge 2020; Knight et al. 2020; Leisova et al. 2006; Winton et al. 2002). Triplex PCR was attempted in the current study to enable confirmation of the presence of *R. collo-cygni* using two DNA regions, reducing the chance of a false positive detection. This is particularly relevant when reporting on a potentially new pathogen incursion. The inclusion of a barley DNA target as an internal positive control added further confidence to the detection system, reducing the possibility of false negatives. This benefit was demonstrated for a selection of *R. collo-cygni* positive leaf DNA samples, where no qPCR detection was initially reported for either the Ram6 or triplex assay. The lack of barley DNA amplification in the triplex assay suggested an inhibitor may have affected the PCR, potentially originating from senesced leaf tissue. While the triplex assay provides robust detection, the use of each assay in uniplex or duplex may also be appropriate for different research objectives.

Development of detection methods across qPCR and ddPCR platforms demonstrated the utility of each technology for sensitive DNA target detection. While each method relies on similar reaction chemistries (The dMIQE Group and Huggett 2020), each has benefits and disadvantages. Quantitative PCR platforms are accessible to a greater number of laboratories compared to ddPCR, require less expensive consumables and can be completed in less time. In comparison, ddPCR does not require a standard curve (although interpretation of results may be informed by inclusion of standards) and reports end-point fluorescence, which may be less ambiguous than late Cq values reported in qPCR (The dMIQE Group and Huggett 2020). Greater sensitivity in ddPCR for low copy number targets has been reported for grape and citrus pathogens (Martinez-Diz et al. 2020; Zhao et al. 2016). In the current study, detection of *R. collo-cygni* in field samples using the triplex assay was similarly reported using both qPCR and ddPCR platforms. The sensitivity of detection across both platforms was 1 pg of *R. collo-cygni* DNA, which was based on detection of *R. collo-cygni* DNA dilutions and non-target fungal DNA templates. While ddPCR offers an alternative platform for detecting PCR products, the similarity in detection and reduced cost and time supports the continued use of qPCR for *R. collo-cygni* detection.

The presence of *R. collo-cygni* DNA in leaves of barley plants grown in New South Wales, South Australia, Tasmania, Victoria and Western Australia indicates a distribution of *R. collo-cygni* encompassing the southern barley growing regions of Australia. Initial reports of *R. collo-cygni* detection in Australia lack detailed information regarding species identification or sampling methods (Biosecurity Tasmania 2020; GRDC 2021; Oxley et al. 2010; Spencer et al. 2019), however they are generally supported by the results of the current study. Infrequent detection of *R. collo-cygni* in leaves from New South Wales and Western Australia and no detection from seed originating from Western Australia suggest *R. collo-cygni* may have a low population density in the areas sampled. In contrast, the levels of *R. collo-cygni* DNA detected in leaves from South Australia, Tasmania and Victoria suggest a greater severity of infection. While this study confirms the presence of *R. collo-cygni* in Australia, assessment of samples from a small number of fields limited the ability to describe the incidence and distribution across barley growing regions. The distribution of *R. collo-cygni* in Australia may be affected by environmental conditions, as RLS is reported to be more severe in cool, wet environments (Dussart et al. 2020; Hoheneder et al. 2021b; Mařík et al. 2011; McGrann and Brown 2018) compared to drier conditions, however distinct disease responses to environmental factors require further investigation.

A likely pathway for introduction of *R. collo-cygni* into Australia is infested barley seed (Harvey 2002; Havis et al. 2014; Matusinsky et al. 2011), with this means of dispersal potentially playing a role in further distribution across the country. Assessment of global and regional *R. collo-cygni* population structure is required to gain an understanding of the pathways for pathogen dispersal (Dussart et al. 2020). The emergence of *R. collo-cygni* in Australia has revealed a range of research questions which must be addressed. The impact of RLS on barley crops in the Australian environment and the potential consequences of climate change at the forefront. These investigations will rely on accurate diagnosis of *R. collo-cygni* in the environment, with the assays described here potentially allowing management options to be implemented, such as those in the UK, where seeds that contain less the 1 pg μL^-1^ of *R. collo-cygni* DNA are recommended for sowing (Oxley et al. 2010). A continued focus on this pathogen and disease will be required in both a regional and global context to understand the implications of its recent recognition as a threat to barley production (Dussart et al. 2020).

## Acknowledgements

This research was supported by Curtin University. We thank E. Ferrari (Curtin University) and F. Tini (University of Perugia) for excellent technical support; and K. Fuhrman (Field Applied Research Australia), N. Poole (Field Applied Research Australia) and K. Jayasena (Department of Primary Industries and Regional Development) for supplying leaf and seed samples.

## Supplementary Figures

**Supplementary Figure S1.**
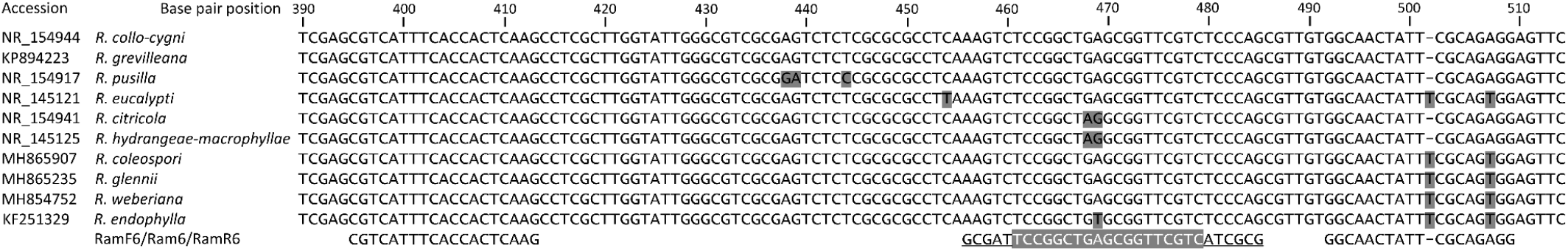
Alignment of primers RamF6 and RamR6 and molecular beacon Ram6 (Taylor et al. 2010) against internal transcribed spacer region sequences of the target fungus *Ramularia collo-cygni* and nine other *Ramularia* species. The nine *Ramularia* species were selected based on similar sequences in the primer and molecular beacon binding regions. *R. collo-cygni* (NR_154944) was used for base pair positions. Bases with grey shading and black text indicate polymorphic sequences. Primers are designated by plain text and the molecular beacon is designated by a grey background with white text (*R. collo-cygni* specific loop sequence) and underlined bases (complementary stem sequences). A (-) indicates the position of an inserted or deleted base in the aligned sequences.

**Supplementary Figure S2.**
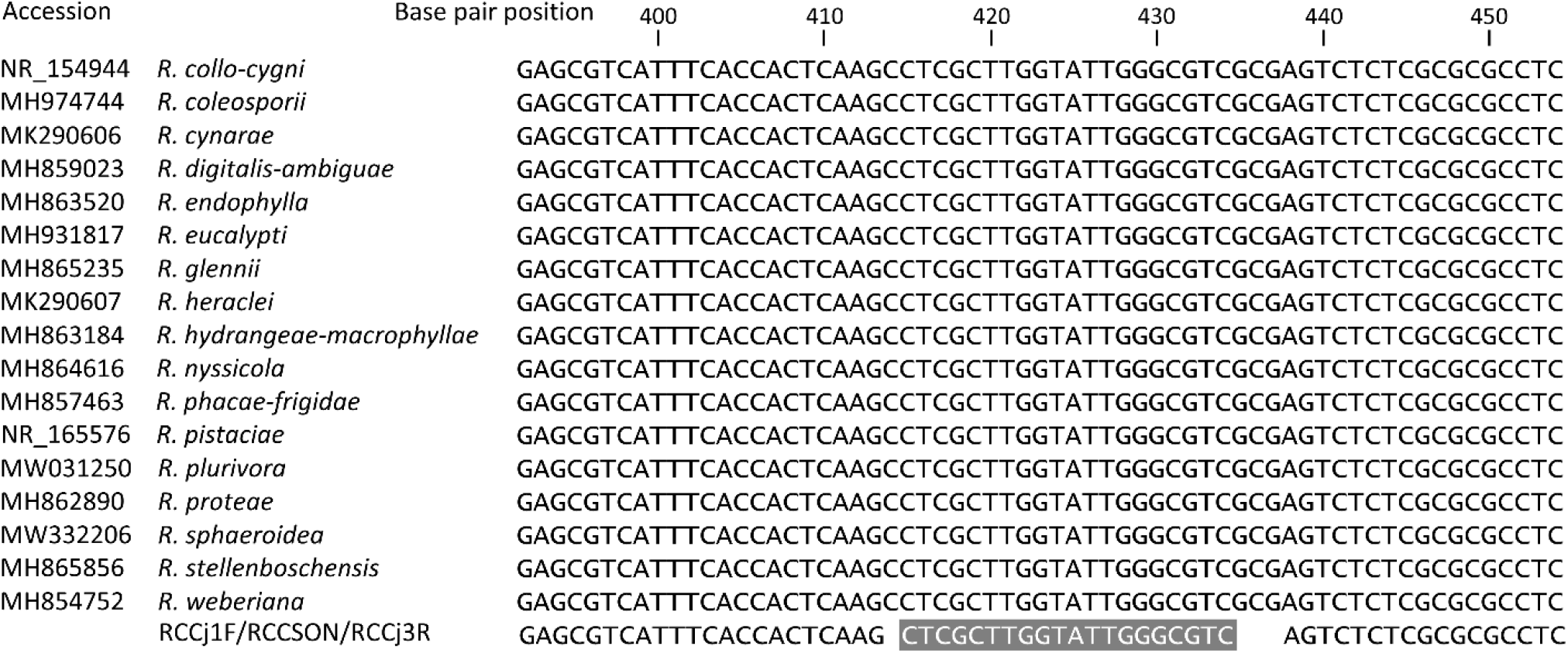
Alignment of primers RCCj1F and RCCj3R and probe RCCSON (Matusinsky et al. 2011) against internal transcribed spacer region sequences of the target fungus *Ramularia collo-cygni* and 16 other *Ramularia* species. The 16 *Ramularia* species were selected based on identical sequences in the primer and probe binding regions. *R. collo-cygni* (NR_154944) was used for base pair positions. Primers are designated by plain text and the probe is designated by a grey background with white text.

**Supplementary Figure S3.**
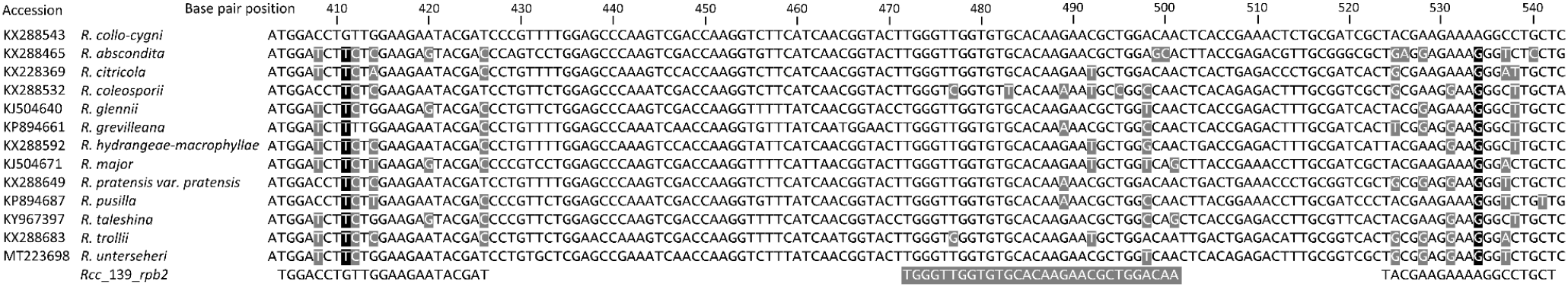
Alignment of the *Rcc*_139_*rpb2* assay primers and probe against RNA polymerase II second largest subunit (*rpb2*) sequences of the target fungus *Ramularia collo-cygni* and 12 other *Ramularia* species. The 12 *Ramularia* species were selected based on similar sequences in the primer and probe binding regions. *R. collo-cygni* (KX288543) was used for base pair positions. Polymorphisms between *R. collo-cygni* and the remaining species sequences are indicated by shading (white text on black indicates a unique *R. collo-cygni* polymorphism and white text on grey indicates a partially discriminating polymorphism). Primers are designated by plain text and the probe is designated by a grey background with white text.

**Supplementary Figure S4.**
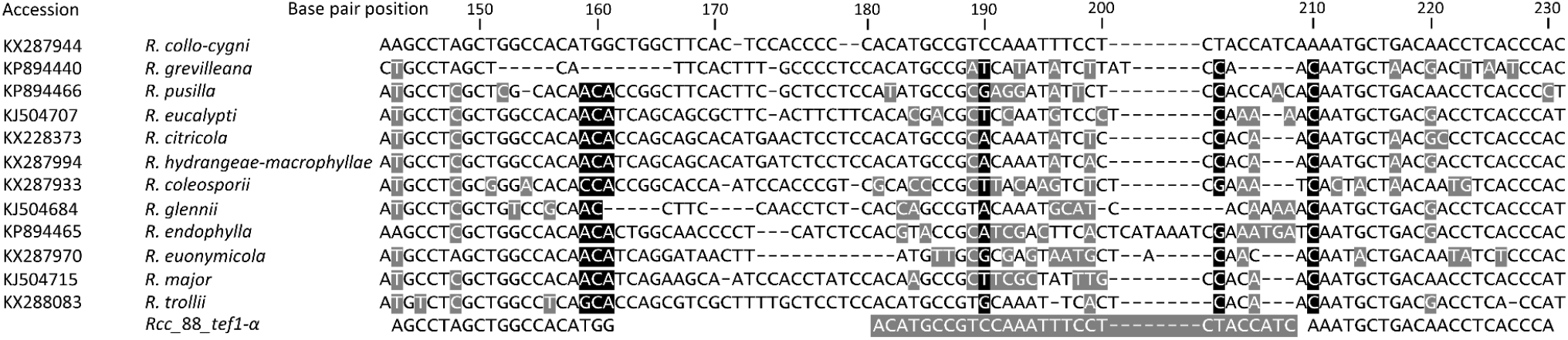
Alignment of the *Rcc*_88_*tef1-α* assay primers and probe against translation elongation factor 1-α (*tef1-α*) sequences of the target fungus *Ramularia collo-cygni* and 11 other *Ramularia* species. The 11 *Ramularia* species were selected based on similar sequences in the primer and probe binding regions. *R. collo-cygni* (KX287944) was used for base pair positions. Polymorphisms between *R. collo-cygni* and the remaining species sequences are indicated by shading (white text on black indicates a unique *R. collo-cygni* polymorphism and white text on grey indicates a partially discriminating polymorphism). Primers are designated by plain text and the probe is designated by a grey background with white text. A (-) indicates the position of an inserted or deleted base in the aligned sequences.

