## Supplementary material for "Detection of *Ramularia collo-cygni* from barley (*Hordeum vulgare*) in Australia using triplex quantitative and digital PCR": Supp Table S1

**Supplementary Table S1.** GenBank accession numbers for DNA sequences of RNA polymerase II second largest subunit (*rpb2*) and translation elongation factor 1-alpha (*tef1-α*) genes of *Ramularia* and related species used for alignments and *Ramularia collo-cygni* specific primer and probe design.

| Species | Culture Number | Host | Country of Origin | GenBank Accession | |
| --- | --- | --- | --- | --- | --- |
|  |  |  |  | *rpb2* | *tef1-α* |
| *Pallidocarcospora heimii* | CPC 11716 | *-* | Brazil | KX348064 | KF903134 |
| *Pa. konae* | CBS 111028 | *Leucadendron* cv. *Safari Sunset* | USA | KX348066 | KF903140 |
| *Pa. crystallina* | CBS 111045 | *Eucalyptus grandis* | South Africa | KX348063 | KF903129 |
| *Pa. acaciigena* | CBS 112515 | *Acacia mangium* | Venezuela | KX348062 | KF903125 |
| *Pa. irregulariramosa* | CBS 111211 | *Eucalyptus saligna* | South Africa | KX348065 | KF903139 |
| *Pseudocercosporella bakari* | CBS 119448 | *Ipomoea indica* | New Zealand | KX288462 | KX287862 |
| *Ramularia abcondita* | CBS 114727 | *Arctium tomentosum* | Sweden | KX288465 | KX287864 |
| *R. acris* | CPC 25898 | *Ranunculus acris* | Netherlands | KX288466 | KX287865 |
| *R. acris* | CPC 25900 | *Ranunculus* sp. | Netherlands | KX288468 | KX287867 |
| *R. acroptili* | CPC 18723 | *Cynara cardunculus* | USA | KX288470 | KX287869 |
| *R. acroptili* | CBS 120252 | *Acroptilon repens* | Turkey | KX288472 | KX287871 |
| *R. acroptili* | CBS 120253 | *Centaurea solstitialis* | Greece | KX288473 | KX287872 |
| *R. actinidia* | CPC 11675 | *Actinidia polygama* | South Korea | KX288474 | KX287873 |
| *R. agostaches* | CPC 10819 | *Agastache rugosa* | South Korea | KX288475 | KX287874 |
| *R. agrimoniae* | CPC 11450 | *Agrimonia pilosa* | South Korea | KX288478 | KX287877 |
| *R. alangiicola* | CPC 10299 | *Alangium platanifolium* | South Korea | KX288483 | KX287882 |
| *R. aplospora* | CBS 545.82 | *Mildew on Alchemilla* | Germany | KP894656 | KP894435 |
| *R. aplospora* | CBS 114118 | *Alchemilla vulgaris* | Sweden | KP894655 | KP894434 |
| *R. archangelicae* | CBS 108991 | *Angelica sylvestris* | Austria | KX288487 | KX287886 |
| *R. archangelicae* | CBS 288.49 | *Angelica sylvestris* | Austria | KX288491 | KX287890 |
| *R. armoraciae* | CBS 241.90 | *Armoracia rusticana* | Germany | KX288492 | KX287891 |
| *R. asteris* | CBS 131.21 | *Aster tripolium* | Netherlands | KX288494 | KX287893 |
| *R. bellunensis* | CBS 118417 | *Argyranthemum frutescens* | New Zealand | KX348078 | KX287894 |
| *R. bellunensis* | CBS 116.43 | *Agrimonia pilosa* | South Korea | KX288495 | KX287895 |
| *R. beticola* | CPC 30065 | *Beta vulgaris* | Denmark | KX288496 | KX287896 |
| *R. bosniaca* | CBS 123880 | *Scabiosa ochroleuca* | Czech Republic | KX288504 | KX287904 |
| *R. buniadis* | CBS 114301 | *Bunias orientalis* | Sweden | KX288506 | KX287906 |
| *R. calsea* | CBS 101613 | *Symphytum* sp. | Germany | KP894658 | KP894437 |
| *R. carneola* | CBS 108975 | *Scrophularia nodosa* | Netherlands | KX288507 | KX287907 |
| *R. cerastiicola* | CBS 115913 | *Cerastium semidecandrum* | Netherlands | KX348079 | KF253180 |
| *R. chamaedryos* | CBS 116577 | *Veronica chamaedrys* | Sweden | KX288512 | KX287912 |
| *R. chamaedryos* | CBS 114731 | *Veronica anagallis-aquatica* | Sweden | KX288516 | KX287916 |
| *R. chelidonii* | CPC 12208 | *Hylomecon vernalis* | South Korea | KX288517 | KX287917 |
| *R. coleosporii* | CPC 10653 | *Coleosporium eupatorii* on *Eupatorium japonicum* | South Korea | KX288520 | KX287920 |
| *R. coleosporii* | CPC 10731 | *Clematidisapiifoliae* on *Clematis apiifolia* | South Korea | KX288522 | KX287922 |
| *R. coleosporii* | CPC 10746 | *Coleosporium eupatorii* on *Eupatorium lindleyanum* | South Korea | KX288525 | KX287925 |
| *R. coleosporii* | CPC 11516 | *Coleosporium plectranthi* on *Plectranthus japonicus* | South Korea | KX288528 | KX287928 |
| *R. coleosporii* | CBS 131753 | *Coleosporium perillae* on *Perilla frutescens var. japonica* | South Korea | KX288529 | KX287929 |
| *R. coleosporii* | CBS 131754 | *Coleosporium asterum* on *Aster pilosus* | South Korea | KX288530 | KX287930 |
| *R. coleosporii* | CBS 131757 | *Coleosporium horianum* on *Codonopsis lanceolata* | South Korea | KX288532 | KX287933 |
| *R. coleosporii* | CBS 131758 | *Coleosporium cacaliae* on *Syneilesis palmata* | South Korea | KX288533 | KX287934 |
| *R. coleosporii* | CBS 131759 | *Coleosporium horianum* on *Codonopsis lanceolata* | South Korea | KX288534 | KX287935 |
| *R. coleosporii* | CBS 131761 | *Coleosporium saussureae* on *Saussurea pulchella* | South Korea | KX288536 | KX287937 |
| *R. coleosporii* | CBS 131762 | *Coleosporium* sp. on *Solidago serotina* | South Korea | KX288537 | KX287938 |
| *R. coleosporii* | CBS 131764 | *Coleosporium eupatorii* on *Eupatorium lindleyanum* | South Korea | KX288539 | KX287940 |
| *R. coleosporii* | CBS 131765 | *Coleosporium asterum* on *Aster pilosus* | South Korea | KX288540 | KX287941 |
| *R. coleosporii* | CBS 131766 | *Coleosporium clerodendri* on *Clerodendron trichotomum* | South Korea | KX288541 | KX287942 |
| *R. coleosporii* | CBS 131767 | *Pileolaria shiraiana* on *Rhus trichocarpa* | South Korea | KX288542 | KX287943 |
| *R. collo-cygni* | CBS 101180 | *Hordeum vulgare* | Austria | KX288543 | KX287944 |
| *R. collo-cygni* | CBS 119439 | *Hordeum vulgare* | Norway | KX288548 | KX287949 |
| *R. collo-cygni* | CBS 119440 | *Hordeum vulgare* | Norway | KX288547 | KX287948 |
| *R. collo-cygni* | CBS 101182 | *Hordeum vulgare* | Germany | KX288544 | KX287945 |
| *R. collo-cygni* | CBS 119441 | *Hordeum vulgare* | Norway | KX288546 | KX287947 |
| *R. collo-cygni* | CBS 119442 | *Hordeum vulgare* | Norway | KX288545 | KX287946 |
| *R. collo-cygni* | CBS 101181 | *Hordeum vulgare* | Germany | KJ504657 | KJ504701 |
| *R. coryli* | CBS 117800 | *Corylus avellana* | Netherlands | KX288549 | KX287950 |
| *R. cupulariae* | CBS 235.73 | *Inula* sp. | Former Czechoslovakia | KX288550 | KX287951 |
| *R. cyclaminicola* | CBS 399.51 | *Cyclamen persicum* | USA | KX288551 | KX287952 |
| *R. cynarae* | CPC 18427 | *Cynara cardunculus* | USA | KX288552 | KX287953 |
| *R. cynarae* | CBS 128779 | *Carthamus tinctorius* | USA | KX288553 | KX287954 |
| *R. cynarae* | CBS 128912 | *Cynara cardunculus* | USA | KX288554 | KX287955 |
| *R. cynarae* | CBS 114728 | *Cirsium arvense* | Sweden | KX288555 | KX287956 |
| *R. cynarae* | CBS 114729 | *Carduus crispus* | Sweden | KX288558 | KX287959 |
| *R. deusta* | CBS 473.50 | *Lathyrus latifolius* | Guadeloupe | KX288559 | KX287960 |
| *R. didyma var. didyma* | CBS 114299 | *Ranunculus repens* | Sweden | KX288560 | KX287961 |
| *R. diervillae* | CPC 16860 | *Diervilla lonicera* | Canada | KX288562 | KX287964 |
| *R. digitalis-ambiguae* | CBS 434.67 | *Digitalis purpurea* | Luxemburg | KX288566 | KX287968 |
| *R. digitalis-ambiguae* | CBS 297.37 | *Digitalis* sp. | Netherlands | KX288567 | KX287969 |
| *R. endophylla* | CBS 113265 | *Quercus robur* | Netherlands | KP894672 | KP894451 |
| *R. endophylla* | CBS 101680 | *Castanea sativa* | Netherlands | KP894680 | KP894458 |
| *R. eucalypti* | CBS 120728 | *Eucalyptus* sp. | Australia | KJ504664 | KJ504708 |
| *R. eucalypti* | CPC 19188 | *Phragmites* sp. | Netherlands | KJ504669 | KJ504713 |
| *R. eucalypti* | CBS 120726 | *Corymbia grandifolia* | Italy | KJ504663 | KJ504707 |
| *R. euonymicola* | CBS 113308 | *Euonymus alatus* | South Korea | KX288568 | KX287970 |
| *R. gaultheriae* | CBS 299.80 | *Gaultheria shallon* | Italy | KX288569 | KX287971 |
| *R. gei* | CBS 344.49 | *Geum urbanum* | Netherlands | KX288570 | KX287972 |
| *R. gei* | CBS 113977 | *Geum* sp. | Sweden | KX288571 | KX287973 |
| *R. geranii* | CBS 159.24 | *Geranium pyrenaicum* | France | KX288572 | KX287974 |
| *R. geranii* | CBS 160.24 | *Geranium sylvaticum* | France | KX288573 | KX287975 |
| *R. geraniicola* | CPC 25912 | *Geranium* sp. | Netherlands | KX288574 | KX287976 |
| *R. glechomatis* | CBS 343.49 | *Glechoma hederacea* | Netherlands | KX288575 | KX287977 |
| *R. glennii* | CBS 129441 | Human bronchial alveolar lavage | Netherlands | KJ504640 | KJ504684 |
| *R. glennii* | CBS 122989 | Human skin | Netherlands | KJ504639 | KJ504683 |
| *R. glennii* | CPC 18468 | Rubber of refrigerator | USA | KJ504646 | KJ504690 |
| *R. glennii* | CBS 120727 | *Corymbia grandifolia* | Italy | KJ504638 | KJ504682 |
| *R. glennii* | CPC 16560 | *Eucalyptus camaldulensis* | Iraq | KJ504643 | KJ504687 |
| *R. grevilleana* | CBS 298.34 | *-* | Netherlands | KP894661 | KP894440 |
| *R. grevilleana* | CBS 719.84 | *Fragaria × ananassa Tioga* | New Zealand | KP894662 | KP894441 |
| *R. grevilleana* | CBS 114732 | *Fragaria ananassa* | Sweden | KP894659 | KP894438 |
| *R. haroldporteri* | CPC 16297 | Unidentified bulb plant | South Africa | KX288577 | KX287979 |
| *R. helminthiae* | CPC 11502 | *Picris hieracioides var.glabrensis* | South Korea | KX288639 | KX288040 |
| *R. heraclei* | CBS 108969 | *Heracleum sphondylium* | Netherlands | KX288578 | KX287980 |
| *R. heraclei* | CBS 108987 | *Heracleum* sp. | Netherlands | KX288580 | KX287982 |
| *R. heraclei* | CPC 11505 | *Heracleum moellendorffii* | South Korea | KX288582 | KX287984 |
| *R. heraclei* | CBS 194.25 | *Pastinaca sativa* | - | KX288586 | KX287988 |
| *R. hieracii-umbellati* | CPC 10690 | *Hieracium umbellatum* | South Korea | KX288587 | KX287989 |
| *R. hydrangea-macrophyllae* | CBS 122273 | *Hydrangea macrophylla* | New Zealand | KX288592 | KX287994 |
| *R. hydrangea-macrophyllae* | CPC 25908 | *Laurus* sp. | Netherlands | KX288593 | KX287995 |
| *R. hydrangea-macrophyllae* | CBS 118410 | *Ligularia clivorum* | New Zealand | KX288594 | KX287996 |
| *R. hydrangea-macrophyllae* | CPC 25905 | *Carex* sp. | Netherlands | KX288595 | KX287997 |
| *R. hydrangea-macrophyllae* | CBS 122625 | *Iris × hollandica hybrid* | New Zealand | KX288596 | KX287998 |
| *R. hydrangea-macrophyllae* | CBS 122272 | *Iris* sp. | New Zealand | KX288597 | KX287999 |
| *R. hydrangea-macrophyllae* | CPC 25902 | *Aesculus hippocastanum* | Netherlands | KX288598 | KX288000 |
| *R. hydrangea-macrophyllae* | CPC 25906 | *Carex* sp. | Netherlands | KX288599 | KX288001 |
| *R. hydrangea-macrophyllae* | CPC 19854 | *Feijoa sellowiana* | Italy | KX288600 | KX288002 |
| *R. hydrangea-macrophyllae* | CPC 19026 | *Phragmites* sp. | Netherlands | KX288601 | KX288003 |
| *R. hydrangea-macrophyllae* | CBS 341.49 | *Angelica sylvestris* | Netherlands | KX288603 | KX288005 |
| *R. hydrangea-macrophyllae* | CPC 25907 | *Juncus* sp. | Netherlands | KX288604 | KX288006 |
| *R. hydrangea-macrophyllae* | CPC 20406 | *Eucalyptus caesia* | USA | KX288605 | KX288007 |
| *R. hydrangea-macrophyllae* | CPC 20484 | *Iris foetidissima* | Netherlands | KX288606 | KX288008 |
| *R. hydrangea-macrophyllae* | CPC 25903 | *Typha* sp. | Netherlands | KX288614 | KX288016 |
| *R. hydrangea-macrophyllae* | CBS 766.84 | *Ulex europaeus* | UK | KX288608 | KX288010 |
| *R. hydrangea-macrophyllae* | CBS 159.82 | *Sparganium ramosum* | Netherlands | KX288609 | KX288011 |
| *R. hydrangea-macrophyllae* | CBS 114117 | *Filipendula vulgaris* | Sweden | KX288611 | KX288013 |
| *R. hydrangea-macrophyllae* | CBS 113614 | *Sparganium ramosum* | Netherlands | KX288613 | KX288015 |
| *R. hydrangea-macrophyllae* | CBS 118408 | *Helleborus niger* | New Zealand | KX288615 | KX288017 |
| *R. inaequalis* | CPC 15815 | *Taraxacum* sp. | Mexico | KX288616 | KX288018 |
| *R. inaequalis* | CPC 25741 | *Taraxacum officinale* | Netherlands | KP894666 | KP894445 |
| *R. inaequalis* | CPC 25742 | *Corylus avellana* | Netherlands | KP894667 | KP894446 |
| *R. kriegeriana* | CPC 10825 | *Plantago asiatica* | South Korea | KX288617 | KX288019 |
| *R. lamii var. lamii* | CBS 108970 | *Lamium album* | Netherlands | KX288620 | KX288022 |
| *R. leonuri* | CPC 11312 | *Leonurus sibiricus* | South Korea | KX348080 | KF253178 |
| *R. lethalis* | CPC 25910 | *Acer pseudoplatanus* | Netherlands | KX288630 | KX288032 |
| *R. ligustrina* | CBS 379.52 | *Ligustrum vulgare* | Italy | KX288631 | KX288033 |
| *R. major* | CPC 12542 | *Petasites japonicus* | South Korea | KX288633 | KX288034 |
| *R. mali* | CBS 129581 | Apple in cold storage | Italy | KJ504649 | KJ504693 |
| *R. malicola* | CBS 119227 | *Malus sp.* | USA | KX288635 | KX288036 |
| *R. miae* | CBS 120121 | *Wachendorﬁa thyrsifolia* | South Africa | KJ504672 | KJ504716 |
| *R. miae* | CPC 19770 | *Leonotis leonurus* | South Africa | KJ504676 | KJ504720 |
| *R. miae* | CPC 19835 | *Gazania rigens* var*. uniﬂora* | South Africa | KJ504675 | KJ504719 |
| *R. neodeusta* | CPC 13568 | *Lathyrus odoratus* | New Zealand | KX288637 | KX288038 |
| *R. neodeusta* | CPC 13567 | *Vicia faba* | New Zealand | KX288638 | KX288039 |
| *R. nyssicola* | CBS 127665 | *Nyssa ogeche × sylvatica hybrid* | USA | KJ504636 | KJ504680 |
| *R. nyssicola* | CBS 127664 | *Nyssa ogeche × sylvatica hybrid* | USA | KP894670 | KP894449 |
| *R. osterici* | CPC 10750 | *Ostericum koreanum* | South Korea | KX288642 | KX288043 |
| *R. parietariae* | CBS 123730 | *Parietaria ofﬁcinalis* | Czech Republic | KX288645 | KX288046 |
| *R. phacae-frigidae* | CBS 234.55 | *Phaca frigida* | Switzerland | KP894671 | KP894450 |
| *R. plurivora* | CBS 118743 | *Human bone marrow* | Netherlands | KJ504651 | KJ504695 |
| *R. plurivora* | CPC 16123 | *Melon in storage* | Netherlands | KJ504653 | KJ504697 |
| *R. pratensis var. pratensis* | CBS 122105 | *Rumex* sp. | Taiwan | KX288647 | KX288048 |
| *R. pratensis var. pratensis* | CPC 16868 | *Verbascum* sp. | Canada | KX288648 | KX288049 |
| *R. pratensis var. pratensis* | CPC 19448 | *Prunus domestica* | - | KX288649 | KX288050 |
| *R. pusilla* | CBS 124973 | *Poa annua* | Germany | KP894687 | KP894466 |
| *R. rhabdospora* | CBS 312.92 | *-* | Germany | KX288651 | KX288051 |
| *R. rhabdospora* | CBS 118415 | *Plantago lanceolata* | New Zealand | KX288652 | KX288052 |
| *R. rubella* | CPC 15748 | *Rumex* sp. | Mexico | KX288653 | KX288053 |
| *R. rubella* | CBS 120161 | *Rumex obtusifolius* | New Zealand | KX288656 | KX288056 |
| *R. rubella* | CBS 114440 | *Rumex longifolius* | Sweden | KX288657 | KX288057 |
| *R. rubella* | CPC 19471 | *Prunus* sp. | Netherlands | KX288658 | KX288058 |
| *R. rufibasis* | CBS 114567 | *Myrica gale* | Sweden | KX288662 | KX288062 |
| *R. rumicicola* | CBS 141118 | *Rumex crispus* | South Korea | KX348081 | KX288063 |
| *R. rumicis* | CBS 114300 | *Rumex aquaticus* | Sweden | KJ504658 | KJ504702 |
| *R. sphaeroidae* | CBS 112891 | *Vicia villosa* subsp. *varia* | USA | KX288673 | KX288074 |
| *R. stellariicola* | CBS 130592 | *Stellaria aquatica* | South Korea | KX288675 | KX288076 |
| *R. stellenboschensis* | CBS 130600 | *Protea* sp*.,*with *Vizella interupta* | South Africa | KX288676 | - |
| *R. tovarae* | CBS 113305 | *Polygonum ﬁliforme* | South Korea | KJ504678 | KJ504722 |
| *R. tricherae* | CBS 108973 | *Knautia arvensis* | Netherlands | KP894688 | KP894467 |
| *R. tricherae* | CBS 108989 | *Knautia dipsacifolia* | Austria | KP894689 | KP894468 |
| *R. tricherae* | CBS 236.73 | *Knautia drymeia* | Former Czechoslovakia | KP894692 | KP894471 |
| *R. trigonotidis* | CPC 14766 | *Trigonotis nakaii* | South Korea | KX288681 | KX288081 |
| *R. trollii* | CBS 109118 | *Trollius europaeus* | Austria | KX288682 | KX288082 |
| *R. unterseheri* | CBS 124846 | *Fagus sylvatica* | Germany | KP894706 | KP894485 |
| *R. unterseheri* | CBS 130721 | Room inside a castle | Germany | KP894710 | KP894489 |
| *R. unterseheri* | CBS 117879 | *Acer pseudoplatanus* | Netherlands | KP894696 | KP894475 |
| *R. unterseheri* | CBS 124884 | *Fagus sylvatica* | Germany | KP894709 | KP894488 |
| *R. uredinicola* | CPC 11852 | *Melampsora sp.* on *Salix babylonica* | Iran | KX288684 | KX288084 |
| *R. uredinicola* | CBS 179.68 | *Melampsora sp.* on *Populus* sp. | Italy | KX288685 | KX288085 |
| *R. uredinicola* | CPC 11482 | *Melampsora* sp*.* on *Salix* sp*.* | South Korea | KX288690 | KX288090 |
| *R. uredinicola* | CBS 131769 | *Melampsora* sp*.* on *Salix gracilistyla* | South Korea | KX288691 | KX288091 |
| *R. uredinicola* | CBS 131770 | *Melampsora* sp*.* on *Populus alba × glandulosa* | South Korea | KX288692 | KX288092 |
| *R. uredinicola* | CBS 131771 | *Melampsora* sp*.* on *Salix koreensis* | South Korea | KX288693 | KX288093 |
| *R. uredinicola* | CBS 131772 | *Melampsora* sp*. on Salix matsudana* for*. tortuosa* | South Korea | - | KX288094 |
| *R. urticae* | CBS 105.26 | *-* | - | KP894715 | KP894494 |
| *R. urticae* | CBS 113974 | *Urtica dioica* | Sweden | KP894714 | KP894493 |
| *R. urticae* | CBS 162.91 | *Urtica dioica* | Germany | KP894716 | KP894495 |
| *R. urticae* | CPC 14807 | *Aconitum pseudo-laeve var. erectum* | South Korea | KX288694 | KX288095 |
| *R. valerianae var. valerianae* | CBS 109122 | *Valeriana* sp*.* | Austria | KX288696 | KX288096 |
| *R. villisumbrosae* | CBS 271.38 | *Narcissus* cv*. Victoria* | UK | KX288698 | KX288098 |
| *R. villisumbrosae* | CBS 272.38 | *Narcissus* cv*.* Golden Spur | UK | KX288699 | KX288099 |
| *R. variabilis* | CPC 16865 | *Verbascum* sp*.* | Canada | KP894717 | KP894496 |
| *R. veronicicola* | CBS 113981 | *Veronica spicata* | Sweden | KX288700 | KX288100 |
| *R. visellae* | CBS 130601 | *Protea* sp*.,*in association with *Vizella interupta* | South Africa | KJ504679 | KJ504723 |
| *R. visellae* | CBS 117798 | *Carpinus betulus* | Netherlands | KP894728 | KP894507 |
| *R. weberiana* | CBS 136.23 | *-* | - | KJ504677 | KJ504721 |
| *R. weigelae* | CBS 113309 | *Weigela subsessilis* | South Korea | KX288701 | KX288101 |
| *Xenoramularia arxii* | CBS 342.49 | *Acorus calamus* | Netherlands | KX288720 | KX288116 |
| *X. polygonicola* | CPC 10854 | *Polygonum* sp*.* | South Korea | KX288723 | KX288120 |
| *X. Neerlandica* | CBS 113615 | *Sparganium ramosum* | Netherlands | KX288721 | KX288117 |
| *X. Neerlandica* | CPC 18378 | *Iris pseudacorus* | Netherlands | KX348108 | KX288119 |
