## Supplementary material for "Detection of *Ramularia collo-cygni* from barley (*Hordeum vulgare*) in Australia using triplex quantitative and digital PCR": Supp Table S2

**Supplementary Table S2.** GenBank accession numbers for DNA sequences of translation elongation factor 1-alpha (*tef1-α*) of *Hordeum vulgare* used for alignments and specific primer and probe design.

| Species | Cultivar | GenBank Accession |
| --- | --- | --- |
| *Hordeum vulgare* | Igri | Z23130 |
| *Hordeum vulgare* | Golden Promise | KP293845 |
| *Hordeum vulgare* | Golden Promise | KP293846 |
| *Hordeum vulgare* | Haruna Nijo | AK354224 |
