## Supplementary material for "Detection of *Ramularia collo-cygni* from barley (*Hordeum vulgare*) in Australia using triplex quantitative and digital PCR": Supp Table S3

**Supplementary Table S3.** Quantification cycles and copy numbers for a ten-fold dilution series of *R. collo-cygni* (*Rcc*) DNA using triplex quantitative PCR (qPCR) and droplet digital PCR (ddPCR), respectively. Values are reported for the *R. collo-cygni*-specific *Rcc*_139_*rpb2* and *Rcc*_88_*tef1-α* assays.

|  | qPCR | |  | ddPCR | |
| --- | --- | --- | --- | --- | --- |
| *Rcc* DNA (ng) | *Rcc*_139_*rpb2* (Cq)^a^ | *Rcc*_88_*tef1-α* (Cq)^a^ |  | *Rcc*_139_*rpb2* (copies µL^-1^)^b^ | *Rcc*_88_*tef1-α* (copies µL^-1^)^b^ |
| 10 | 20.8 ± 0.02 | 21.1 ± 0.02 |  | 8515.4 ± 286.68 | 11202.1 ± 318.05 |
| 1 | 24.3 ± 0.04 | 24.4 ± 0.02 |  | 1053.2 ± 27.13 | 1052.8 ± 32.21 |
| 0.1 | 27.6 ± 0.08 | 27.8 ± 0.06 |  | 107 ± 1.56 | 109.2 ± 0.56 |
| 0.01 | 31 ± 0.05 | 31.1 ± 0.05 |  | 11.2 ± 0.26 | 11.3 ± 0.89 |
| 0.001 | 34.2 ± 0.16 | 34.7 ± 0.29 |  | 1.1 ± 0.16 | 1.4 ± 0.16 |
| 0.0001 | 38.7 ± 0.01^c^ | 37.9 ± 0.58 |  | 0.1 ± 0.05^c^ | 0.2 ± 0.05^c^ |
| 0.00001 | —^d^ | — |  | 0.06^e^ | — |

^a^ Mean ± standard error of the quantification cycle (Cq) (*n* = 3).

^b^ Mean ± standard error of the number of copies per microliter (*n* = 3).

^c^ Detection failed for one of the replicates (*n* = 2).

^d^ A dash (—) indicates that detection failed for all of the replicates (*n* = 0).

^e^ Detection failed for two of the replicates (*n* = 1).
