## Supplementary material for "Detection of *Ramularia collo-cygni* from barley (*Hordeum vulgare*) in Australia using triplex quantitative and digital PCR": Supp Tables S4

**Supplementary Table S4.** Mean quantification cycle (Cq) and DNA (pg) values with standard errors (*n* = 2) for *Hordeum vulgare* and *Ramularia collo-cygni* DNA in leaf samples from New South Wales (NSW), South Australia (SA), Tasmania (Tas), Victoria (Vic) and Western Australia (WA) using *R. collo-cygni*-specific triplex and Ram6 quantitative PCR assays.

|  |  |  |  |  | *Hv*_116_*tef1-α* |  | *Rcc*_139_*rpb2* | | *Rcc*_88_*tef1-α* | |  | Ram6 (ITS) | |
| --- | --- | --- | --- | --- | --- | --- | --- | --- | --- | --- | --- | --- | --- |
| Sample | Dilution^a^ | Region | State | Year | Cq |  | Cq | DNA (pg) | Cq | DNA (pg) |  | Cq | DNA (pg) |
| 16MUL_001 | — | Mulwala | NSW | 2016 | 32.3 ± 0.09 |  | 0 ± 0 | 0 ± 0 | 0 ± 0 | 0 ± 0 |  | 0 ± 0 | 0 ± 0 |
| 16MUL_002 | — | Mulwala | NSW | 2016 | 35 ± 0.62 |  | 0 ± 0 | 0 ± 0 | 0 ± 0 | 0 ± 0 |  | 0 ± 0 | 0 ± 0 |
| 16MUL_004 | — | Mulwala | NSW | 2016 | 24 ± 0.16 |  | 0 ± 0 | 0 ± 0 | 0 ± 0 | 0 ± 0 |  | 0 ± 0 | 0 ± 0 |
| 16MUL_005 | — | Mulwala | NSW | 2016 | 21.2 ± 0.04 |  | 30.8 ± 0.21 | 8 ± 0 | 31.6 ± 0.19 | 4 ± 0 |  | 27.9 ± 0.49 | 3 ± 0 |
| 16MUL_007 | — | Mulwala | NSW | 2016 | 20.5 ± 0.02 |  | 0 ± 0 | 0 ± 0 | 0 ± 0 | 0 ± 0 |  | 33.9 ± 0.5 | 0 ± 0 |
| 16MUL_008 | — | Mulwala | NSW | 2016 | 27.4 ± 0.09 |  | 0 ± 0 | 0 ± 0 | 0 ± 0 | 0 ± 0 |  | 33.2 ± 0.02 | 0 ± 0 |
| 16MUL_001 | 10 | Mulwala | NSW | 2016 | 35 ± 2.12 |  | 0 ± 0 | 0 ± 0 | 0 ± 0 | 0 ± 0 |  | 0 ± 0 | 0 ± 0 |
| 16MUL_002 | 10 | Mulwala | NSW | 2016 | 0 ± 0 |  | 0 ± 0 | 0 ± 0 | 0 ± 0 | 0 ± 0 |  | 0 ± 0 | 0 ± 0 |
| 16MUL_004 | 10 | Mulwala | NSW | 2016 | 26.3 ± 0.14 |  | 0 ± 0 | 0 ± 0 | 0 ± 0 | 0 ± 0 |  | 0 ± 0 | 0 ± 0 |
| 16MUL_005 | 10 | Mulwala | NSW | 2016 | 23.9 ± 0.12 |  | 0 ± 0 | 0 ± 0 | 0 ± 0 | 0 ± 0 |  | 33.2 ± 0.02 | 0 ± 0 |
| 16MUL_007 | 10 | Mulwala | NSW | 2016 | 23 ± 0.01 |  | 0 ± 0 | 0 ± 0 | 0 ± 0 | 0 ± 0 |  | 0 ± 0 | 0 ± 0 |
| 16MUL_008 | 10 | Mulwala | NSW | 2016 | 29.9 ± 0.35 |  | 0 ± 0 | 0 ± 0 | 0 ± 0 | 0 ± 0 |  | 38.6 ± 1.05 | 0 ± 0 |
| 16DAI_001 | — | Dairy Plains | Tas | 2016 | 20.8 ± 0.11 |  | 23 ± 0.14 | 1010 ± 0.09 | 23.5 ± 0.08 | 1070 ± 0.06 |  | 19.2 ± 0.04 | 631 ± 0.02 |
| 16DAI_002 | — | Dairy Plains | Tas | 2016 | 24 ± 0.08 |  | 26.5 ± 0.08 | 114 ± 0.01 | 26.6 ± 0.05 | 135 ± 0 |  | 22.3 ± 0.25 | 91 ± 0.01 |
| 16DAI_003 | — | Dairy Plains | Tas | 2016 | 30 ± 0.04 |  | 32.2 ± 0.31 | 3 ± 0 | 32.5 ± 0.02 | 3 ± 0 |  | 28.2 ± 0.06 | 2 ± 0 |
| 16DAI_004 | — | Dairy Plains | Tas | 2016 | 21.2 ± 0.11 |  | 29 ± 0.11 | 24 ± 0 | 30.1 ± 0.01 | 13 ± 0 |  | 26.4 ± 0.05 | 7 ± 0 |
| 16DAI_005 | — | Dairy Plains | Tas | 2016 | 24 ± 0.05 |  | 25.8 ± 0.01 | 176 ± 0 | 26 ± 0.03 | 202 ± 0 |  | 21.7 ± 0.12 | 134 ± 0.01 |
| 16DAI_006 | — | Dairy Plains | Tas | 2016 | 21.4 ± 0.1 |  | 23.9 ± 0.2 | 580 ± 0.07 | 24.3 ± 0.06 | 637 ± 0.03 |  | 20.2 ± 0.08 | 337 ± 0.02 |
| 16DAI_007 | — | Dairy Plains | Tas | 2016 | 23.5 ± 0.03 |  | 30.7 ± 0.26 | 8 ± 0 | 31.3 ± 0 | 5 ± 0 |  | 27.2 ± 0.14 | 4 ± 0 |
| 16DAI_008 | — | Dairy Plains | Tas | 2016 | 26.4 ± 0.02 |  | 27.1 ± 0.03 | 79 ± 0 | 27.1 ± 0 | 99 ± 0 |  | 22.8 ± 0.02 | 66 ± 0 |
| 16DAI_009 | — | Dairy Plains | Tas | 2016 | 25.3 ± 0.09 |  | 25.6 ± 0.25 | 202 ± 0.03 | 25.6 ± 0.17 | 272 ± 0.03 |  | 21.4 ± 0.06 | 158 ± 0.01 |
| 16DAI_001 | 10 | Dairy Plains | Tas | 2016 | 23.1 ± 0.12 |  | 26.5 ± 0.07 | 210 ± 0.01 | 27 ± 0.02 | 190 ± 0 |  | 23.5 ± 0.01 | 34 ± 0 |
| 16DAI_002 | 10 | Dairy Plains | Tas | 2016 | 26.2 ± 0.03 |  | 30.6 ± 0.7 | 23 ± 0.01 | 30 ± 0.11 | 33 ± 0 |  | 26.2 ± 0.07 | 7 ± 0 |
| 16DAI_003 | 10 | Dairy Plains | Tas | 2016 | 0 ± 0 |  | 0 ± 0 | 0 ± 0 | 0 ± 0 | 0 ± 0 |  | 32.4 ± 0.03 | 0 ± 0 |
| 16DAI_004 | 10 | Dairy Plains | Tas | 2016 | 23.4 ± 0 |  | 32.4 ± 0.31 | 8 ± 0 | 34.9 ± 0.22 | 2 ± 0 |  | 30.2 ± 0.11 | 1 ± 0 |
| 16DAI_005 | 10 | Dairy Plains | Tas | 2016 | 26.1 ± 0.33 |  | 29.5 ± 0.12 | 40 ± 0 | 29.6 ± 0.14 | 43 ± 0 |  | 25.7 ± 0.05 | 9 ± 0 |
| 16DAI_006 | 10 | Dairy Plains | Tas | 2016 | 23.7 ± 0.13 |  | 27.5 ± 0.07 | 117 ± 0 | 27.8 ± 0.06 | 117 ± 0 |  | 24.2 ± 0.17 | 22 ± 0 |
| 16DAI_007 | 10 | Dairy Plains | Tas | 2016 | 25.7 ± 0.13 |  | 35.1 ± 0.04 | 2 ± 0 | 34.8 ± 0.07 | 2 ± 0 |  | 31.3 ± 0.02 | 0 ± 0 |
| 16DAI_008 | 10 | Dairy Plains | Tas | 2016 | 28.9 ± 0.16 |  | 30.5 ± 0.07 | 23 ± 0 | 30.6 ± 0.05 | 23 ± 0 |  | 27.2 ± 0.13 | 4 ± 0 |
| 16DAI_009 | 10 | Dairy Plains | Tas | 2016 | 27.5 ± 0.11 |  | 29.1 ± 0.07 | 50 ± 0 | 29.1 ± 0 | 55 ± 0 |  | 25.6 ± 0.07 | 9 ± 0 |
| 16HAG_005 | — | Hagley | Tas | 2016 | 25.1 ± 0.05 |  | 0 ± 0 | 0 ± 0 | 0 ± 0 | 0 ± 0 |  | 32.4 ± 0.13 | 0 ± 0 |
| 16HAG_007 | — | Hagley | Tas | 2016 | 19.4 ± 0.22 |  | 24.3 ± 0.25 | 435 ± 0.07 | 25.1 ± 0.13 | 382 ± 0.03 |  | 19.6 ± 0.97 | 602 ± 0.33 |
| 16HAG_008 | — | Hagley | Tas | 2016 | 20.6 ± 0.07 |  | 25.6 ± 0.12 | 202 ± 0.02 | 26.1 ± 0.08 | 187 ± 0.01 |  | 22.1 ± 0.14 | 104 ± 0.01 |
| 16HAG_009 | — | Hagley | Tas | 2016 | 18.4 ± 0.15 |  | 31.8 ± 0.19 | 4 ± 0 | 32.8 ± 0.13 | 2 ± 0 |  | 28.6 ± 0.15 | 2 ± 0 |
| 16HAG_013 | — | Hagley | Tas | 2016 | 23.4 ± 0.02 |  | 33.3 ± 0.37 | 2 ± 0 | 32.7 ± 0.02 | 2 ± 0 |  | 28.8 ± 0.22 | 2 ± 0 |
| 16HAG_014 | — | Hagley | Tas | 2016 | 22 ± 0.05 |  | 25.1 ± 0 | 261 ± 0 | 25.6 ± 0 | 259 ± 0 |  | 21.9 ± 0.05 | 116 ± 0 |
| 16HAG_015 | — | Hagley | Tas | 2016 | 22.5 ± 0.08 |  | 28.9 ± 0.05 | 25 ± 0 | 30 ± 0.02 | 13 ± 0 |  | 26 ± 0.04 | 9 ± 0 |
| 16HAG_019 | — | Hagley | Tas | 2016 | 18.8 ± 0.06 |  | 0 ± 0 | 0 ± 0 | 0 ± 0 | 0 ± 0 |  | 24.3 ± 0.1 | 26 ± 0 |
| 16HAG_020 | — | Hagley | Tas | 2016 | 29.2 ± 0.13 |  | 0 ± 0 | 0 ± 0 | 0 ± 0 | 0 ± 0 |  | 0 ± 0 | 0 ± 0 |
| 16HAG_026 | — | Hagley | Tas | 2016 | 26.1 ± 0.09 |  | 31.8 ± 0.3 | 4 ± 0 | 31.9 ± 0.21 | 4 ± 0 |  | 27.5 ± 0.31 | 4 ± 0 |
| 16HAG_005 | 10 | Hagley | Tas | 2016 | 27.3 ± 0.06 |  | 0 ± 0 | 0 ± 0 | 0 ± 0 | 0 ± 0 |  | 0 ± 0 | 0 ± 0 |
| 16HAG_007 | 10 | Hagley | Tas | 2016 | 21.8 ± 0.07 |  | 27.9 ± 0.01 | 98 ± 0 | 28.4 ± 0.02 | 86 ± 0 |  | 24.3 ± 0.05 | 21 ± 0 |
| 16HAG_008 | 10 | Hagley | Tas | 2016 | 22.9 ± 0.18 |  | 0 ± 0 | 0 ± 0 | 30 ± 0.49 | 34 ± 0.01 |  | 26 ± 0.13 | 8 ± 0 |
| 16HAG_009 | 10 | Hagley | Tas | 2016 | 20.9 ± 0.06 |  | 0 ± 0 | 0 ± 0 | 0 ± 0 | 0 ± 0 |  | 33 ± 0.09 | 0 ± 0 |
| 16HAG_013 | 10 | Hagley | Tas | 2016 | 25.8 ± 0.01 |  | 0 ± 0 | 0 ± 0 | 0 ± 0 | 0 ± 0 |  | 33.3 ± 0.12 | 0 ± 0 |
| 16HAG_014 | 10 | Hagley | Tas | 2016 | 24.4 ± 0.12 |  | 28.9 ± 0.1 | 55 ± 0 | 29.2 ± 0.01 | 52 ± 0 |  | 25.9 ± 0.11 | 8 ± 0 |
| 16HAG_015 | 10 | Hagley | Tas | 2016 | 24.9 ± 0.05 |  | 33.1 ± 0.48 | 6 ± 0 | 33.5 ± 0.1 | 4 ± 0 |  | 30.6 ± 0.06 | 0 ± 0 |
| 16HAG_019 | 10 | Hagley | Tas | 2016 | 20.9 ± 0.08 |  | 30.6 ± 0.33 | 23 ± 0 | 32.1 ± 0.31 | 10 ± 0 |  | 28.3 ± 0.13 | 2 ± 0 |
| 16HAG_020 | 10 | Hagley | Tas | 2016 | 34.6 ± 2.57 |  | 0 ± 0 | 0 ± 0 | 0 ± 0 | 0 ± 0 |  | 0 ± 0 | 0 ± 0 |
| 16HAG_026 | 10 | Hagley | Tas | 2016 | 28.2 ± 0.26 |  | 0 ± 0 | 0 ± 0 | 0 ± 0 | 0 ± 0 |  | 31.6 ± 0 | 0 ± 0 |
| RLS_001 | — | South Stirling | WA | 2018 | 0 ± 0 |  | 0 ± 0 | 0 ± 0 | 0 ± 0 | 0 ± 0 |  | 0 ± 0 | 0 ± 0 |
| RLS_002 | — | South Stirling | WA | 2018 | 0 ± 0 |  | 0 ± 0 | 0 ± 0 | 0 ± 0 | 0 ± 0 |  | 0 ± 0 | 0 ± 0 |
| RLS_003 | — | South Stirling | WA | 2018 | 26.8 ± 0.01 |  | 0 ± 0 | 0 ± 0 | 0 ± 0 | 0 ± 0 |  | 0 ± 0 | 0 ± 0 |
| RLS_004 | — | South Stirling | WA | 2018 | 0 ± 0 |  | 0 ± 0 | 0 ± 0 | 0 ± 0 | 0 ± 0 |  | 0 ± 0 | 0 ± 0 |
| RLS_005 | — | South Stirling | WA | 2018 | 0 ± 0 |  | 0 ± 0 | 0 ± 0 | 0 ± 0 | 0 ± 0 |  | 0 ± 0 | 0 ± 0 |
| RLS_006 | — | South Stirling | WA | 2018 | 19.5 ± 0.04 |  | 0 ± 0 | 0 ± 0 | 0 ± 0 | 0 ± 0 |  | 0 ± 0 | 0 ± 0 |
| RLS_007 | — | South Stirling | WA | 2018 | 24 ± 0.05 |  | 0 ± 0 | 0 ± 0 | 0 ± 0 | 0 ± 0 |  | 0 ± 0 | 0 ± 0 |
| RLS_008 | — | South Stirling | WA | 2018 | 23.5 ± 0.04 |  | 0 ± 0 | 0 ± 0 | 0 ± 0 | 0 ± 0 |  | 0 ± 0 | 0 ± 0 |
| RLS_009 | — | South Stirling | WA | 2018 | 18.6 ± 0.08 |  | 0 ± 0 | 0 ± 0 | 0 ± 0 | 0 ± 0 |  | 0 ± 0 | 0 ± 0 |
| RLS_010 | — | South Stirling | WA | 2018 | 0 ± 0 |  | 0 ± 0 | 0 ± 0 | 0 ± 0 | 0 ± 0 |  | 0 ± 0 | 0 ± 0 |
| RLS_011 | — | South Stirling | WA | 2018 | 17.9 ± 0.03 |  | 0 ± 0 | 0 ± 0 | 0 ± 0 | 0 ± 0 |  | 0 ± 0 | 0 ± 0 |
| RLS_012 | — | South Stirling | WA | 2018 | 21.6 ± 0 |  | 0 ± 0 | 0 ± 0 | 0 ± 0 | 0 ± 0 |  | 0 ± 0 | 0 ± 0 |
| RLS_013 | — | South Stirling | WA | 2018 | 18.4 ± 0.11 |  | 0 ± 0 | 0 ± 0 | 0 ± 0 | 0 ± 0 |  | 0 ± 0 | 0 ± 0 |
| RLS_014 | — | South Stirling | WA | 2018 | 0 ± 0 |  | 0 ± 0 | 0 ± 0 | 0 ± 0 | 0 ± 0 |  | 0 ± 0 | 0 ± 0 |
| RLS_015 | — | South Stirling | WA | 2018 | 21.9 ± 0.01 |  | 0 ± 0 | 0 ± 0 | 0 ± 0 | 0 ± 0 |  | 0 ± 0 | 0 ± 0 |
| RLS_016 | — | South Stirling | WA | 2018 | 21.4 ± 0 |  | 0 ± 0 | 0 ± 0 | 0 ± 0 | 0 ± 0 |  | 0 ± 0 | 0 ± 0 |
| RLS_017 | — | South Stirling | WA | 2018 | 19.5 ± 0.05 |  | 0 ± 0 | 0 ± 0 | 0 ± 0 | 0 ± 0 |  | 0 ± 0 | 0 ± 0 |
| RLS_018 | — | South Stirling | WA | 2018 | 22.1 ± 0.03 |  | 0 ± 0 | 0 ± 0 | 0 ± 0 | 0 ± 0 |  | 0 ± 0 | 0 ± 0 |
| RLS_019 | — | South Stirling | WA | 2018 | 18.6 ± 0.06 |  | 0 ± 0 | 0 ± 0 | 0 ± 0 | 0 ± 0 |  | 0 ± 0 | 0 ± 0 |
| RLS_020 | — | South Stirling | WA | 2018 | 22.6 ± 0 |  | 0 ± 0 | 0 ± 0 | 0 ± 0 | 0 ± 0 |  | 0 ± 0 | 0 ± 0 |
| RLS_021 | — | South Stirling | WA | 2018 | 21 ± 0.06 |  | 0 ± 0 | 0 ± 0 | 0 ± 0 | 0 ± 0 |  | 0 ± 0 | 0 ± 0 |
| RLS_022 | — | South Stirling | WA | 2018 | 20.8 ± 0 |  | 0 ± 0 | 0 ± 0 | 0 ± 0 | 0 ± 0 |  | 0 ± 0 | 0 ± 0 |
| RLS_023 | — | South Stirling | WA | 2018 | 24 ± 0.03 |  | 0 ± 0 | 0 ± 0 | 0 ± 0 | 0 ± 0 |  | 0 ± 0 | 0 ± 0 |
| RLS_024 | — | South Stirling | WA | 2018 | 20.2 ± 0.07 |  | 0 ± 0 | 0 ± 0 | 0 ± 0 | 0 ± 0 |  | 0 ± 0 | 0 ± 0 |
| RLS_025 | — | South Stirling | WA | 2018 | 25.8 ± 0.06 |  | 0 ± 0 | 0 ± 0 | 0 ± 0 | 0 ± 0 |  | 0 ± 0 | 0 ± 0 |
| RLS_026 | — | South Stirling | WA | 2018 | 0 ± 0 |  | 0 ± 0 | 0 ± 0 | 0 ± 0 | 0 ± 0 |  | 0 ± 0 | 0 ± 0 |
| RLS_027 | — | South Stirling | WA | 2018 | 18.7 ± 0.04 |  | 0 ± 0 | 0 ± 0 | 0 ± 0 | 0 ± 0 |  | 0 ± 0 | 0 ± 0 |
| RLS_028 | — | South Stirling | WA | 2018 | 0 ± 0 |  | 0 ± 0 | 0 ± 0 | 0 ± 0 | 0 ± 0 |  | 0 ± 0 | 0 ± 0 |
| RLS_029 | — | South Stirling | WA | 2018 | 23.3 ± 0.06 |  | 0 ± 0 | 0 ± 0 | 0 ± 0 | 0 ± 0 |  | 34.3 ± 0.3 | 1 ± 0 |
| RLS_030 | — | South Stirling | WA | 2018 | 25.5 ± 0.12 |  | 0 ± 0 | 0 ± 0 | 0 ± 0 | 0 ± 0 |  | 29.4 ± 0.55 | 6 ± 0 |
| RLS_141 | — | South Stirling | WA | 2018 | 20 ± 0.01 |  | 0 ± 0 | 0 ± 0 | 0 ± 0 | 0 ± 0 |  | 0 ± 0 | 0 ± 0 |
| RLS_142 | — | South Stirling | WA | 2018 | 19.5 ± 0.11 |  | 0 ± 0 | 0 ± 0 | 0 ± 0 | 0 ± 0 |  | 0 ± 0 | 0 ± 0 |
| RLS_143 | — | South Stirling | WA | 2018 | 21.1 ± 0.04 |  | 0 ± 0 | 0 ± 0 | 0 ± 0 | 0 ± 0 |  | 0 ± 0 | 0 ± 0 |
| RLS_144 | — | South Stirling | WA | 2018 | 19.5 ± 0.03 |  | 0 ± 0 | 0 ± 0 | 0 ± 0 | 0 ± 0 |  | 0 ± 0 | 0 ± 0 |
| RLS_145 | — | South Stirling | WA | 2018 | 20.2 ± 0 |  | 0 ± 0 | 0 ± 0 | 0 ± 0 | 0 ± 0 |  | 0 ± 0 | 0 ± 0 |
| RLS_146 | — | South Stirling | WA | 2018 | 21.1 ± 0.03 |  | 0 ± 0 | 0 ± 0 | 0 ± 0 | 0 ± 0 |  | 0 ± 0 | 0 ± 0 |
| RLS_147 | — | South Stirling | WA | 2018 | 19.2 ± 0.09 |  | 0 ± 0 | 0 ± 0 | 0 ± 0 | 0 ± 0 |  | 0 ± 0 | 0 ± 0 |
| RLS_148 | — | South Stirling | WA | 2018 | 19.5 ± 0.01 |  | 0 ± 0 | 0 ± 0 | 0 ± 0 | 0 ± 0 |  | 0 ± 0 | 0 ± 0 |
| RLS_149 | — | South Stirling | WA | 2018 | 19.4 ± 0.07 |  | 0 ± 0 | 0 ± 0 | 0 ± 0 | 0 ± 0 |  | 0 ± 0 | 0 ± 0 |
| RLS_150 | — | South Stirling | WA | 2018 | 19.8 ± 0.04 |  | 0 ± 0 | 0 ± 0 | 0 ± 0 | 0 ± 0 |  | 0 ± 0 | 0 ± 0 |
| RLS_151 | — | South Stirling | WA | 2018 | 21.1 ± 0.01 |  | 0 ± 0 | 0 ± 0 | 0 ± 0 | 0 ± 0 |  | 0 ± 0 | 0 ± 0 |
| RLS_152 | — | South Stirling | WA | 2018 | 20.4 ± 0.01 |  | 0 ± 0 | 0 ± 0 | 0 ± 0 | 0 ± 0 |  | 0 ± 0 | 0 ± 0 |
| RLS_153 | — | South Stirling | WA | 2018 | 21.1 ± 0.01 |  | 0 ± 0 | 0 ± 0 | 0 ± 0 | 0 ± 0 |  | 0 ± 0 | 0 ± 0 |
| RLS_154 | — | South Stirling | WA | 2018 | 21.5 ± 0.1 |  | 0 ± 0 | 0 ± 0 | 0 ± 0 | 0 ± 0 |  | 0 ± 0 | 0 ± 0 |
| RLS_155 | — | South Stirling | WA | 2018 | 20.7 ± 0.22 |  | 0 ± 0 | 0 ± 0 | 0 ± 0 | 0 ± 0 |  | 0 ± 0 | 0 ± 0 |
| RLS_156 | — | South Stirling | WA | 2018 | 20.7 ± 0.05 |  | 0 ± 0 | 0 ± 0 | 0 ± 0 | 0 ± 0 |  | 0 ± 0 | 0 ± 0 |
| RLS_157 | — | South Stirling | WA | 2018 | 20.2 ± 0 |  | 0 ± 0 | 0 ± 0 | 0 ± 0 | 0 ± 0 |  | 0 ± 0 | 0 ± 0 |
| RLS_158 | — | South Stirling | WA | 2018 | 20.9 ± 0.04 |  | 0 ± 0 | 0 ± 0 | 0 ± 0 | 0 ± 0 |  | 0 ± 0 | 0 ± 0 |
| RLS_159 | — | South Stirling | WA | 2018 | 21.1 ± 0.02 |  | 0 ± 0 | 0 ± 0 | 0 ± 0 | 0 ± 0 |  | 0 ± 0 | 0 ± 0 |
| RLS_160 | — | South Stirling | WA | 2018 | 19.9 ± 0.02 |  | 0 ± 0 | 0 ± 0 | 0 ± 0 | 0 ± 0 |  | 0 ± 0 | 0 ± 0 |
| RLS_161 | — | South Stirling | WA | 2018 | 21.3 ± 0.06 |  | 0 ± 0 | 0 ± 0 | 0 ± 0 | 0 ± 0 |  | 0 ± 0 | 0 ± 0 |
| RLS_162 | — | South Stirling | WA | 2018 | 22 ± 0.03 |  | 0 ± 0 | 0 ± 0 | 0 ± 0 | 0 ± 0 |  | 0 ± 0 | 0 ± 0 |
| RLS_163 | — | South Stirling | WA | 2018 | 21.1 ± 0.02 |  | 0 ± 0 | 0 ± 0 | 0 ± 0 | 0 ± 0 |  | 0 ± 0 | 0 ± 0 |
| RLS_164 | — | South Stirling | WA | 2018 | 20.7 ± 0.02 |  | 0 ± 0 | 0 ± 0 | 0 ± 0 | 0 ± 0 |  | 0 ± 0 | 0 ± 0 |
| RLS_165 | — | South Stirling | WA | 2018 | 20.4 ± 0.07 |  | 0 ± 0 | 0 ± 0 | 0 ± 0 | 0 ± 0 |  | 0 ± 0 | 0 ± 0 |
| RLS_166 | — | South Stirling | WA | 2018 | 21 ± 0.12 |  | 0 ± 0 | 0 ± 0 | 0 ± 0 | 0 ± 0 |  | 0 ± 0 | 0 ± 0 |
| RLS_167 | — | South Stirling | WA | 2018 | 19.9 ± 0.02 |  | 0 ± 0 | 0 ± 0 | 0 ± 0 | 0 ± 0 |  | 0 ± 0 | 0 ± 0 |
| RLS_168 | — | South Stirling | WA | 2018 | 21 ± 0.04 |  | 0 ± 0 | 0 ± 0 | 0 ± 0 | 0 ± 0 |  | 0 ± 0 | 0 ± 0 |
| RLS_169 | — | South Stirling | WA | 2018 | 20.1 ± 0 |  | 0 ± 0 | 0 ± 0 | 0 ± 0 | 0 ± 0 |  | 0 ± 0 | 0 ± 0 |
| RLS_170 | — | South Stirling | WA | 2018 | 20.4 ± 0.07 |  | 0 ± 0 | 0 ± 0 | 0 ± 0 | 0 ± 0 |  | 0 ± 0 | 0 ± 0 |
| RLS_001 | 10 | South Stirling | WA | 2018 | 28.1 ± 0.02 |  | 0 ± 0 | 0 ± 0 | 0 ± 0 | 0 ± 0 |  | 0 ± 0 | 0 ± 0 |
| RLS_002 | 10 | South Stirling | WA | 2018 | 27.3 ± 0.02 |  | 0 ± 0 | 0 ± 0 | 0 ± 0 | 0 ± 0 |  | 0 ± 0 | 0 ± 0 |
| RLS_003 | 10 | South Stirling | WA | 2018 | 31.5 ± 0.03 |  | 0 ± 0 | 0 ± 0 | 0 ± 0 | 0 ± 0 |  | 0 ± 0 | 0 ± 0 |
| RLS_004 | 10 | South Stirling | WA | 2018 | 30.8 ± 0.12 |  | 0 ± 0 | 0 ± 0 | 0 ± 0 | 0 ± 0 |  | 0 ± 0 | 0 ± 0 |
| RLS_005 | 10 | South Stirling | WA | 2018 | 29.3 ± 0.03 |  | 0 ± 0 | 0 ± 0 | 0 ± 0 | 0 ± 0 |  | 0 ± 0 | 0 ± 0 |
| RLS_006 | 10 | South Stirling | WA | 2018 | 23.3 ± 0.01 |  | 0 ± 0 | 0 ± 0 | 0 ± 0 | 0 ± 0 |  | 0 ± 0 | 0 ± 0 |
| RLS_007 | 10 | South Stirling | WA | 2018 | 27.9 ± 0.05 |  | 0 ± 0 | 0 ± 0 | 0 ± 0 | 0 ± 0 |  | 0 ± 0 | 0 ± 0 |
| RLS_008 | 10 | South Stirling | WA | 2018 | 26.7 ± 0.01 |  | 0 ± 0 | 0 ± 0 | 0 ± 0 | 0 ± 0 |  | 0 ± 0 | 0 ± 0 |
| RLS_009 | 10 | South Stirling | WA | 2018 | 23.4 ± 0.09 |  | 0 ± 0 | 0 ± 0 | 0 ± 0 | 0 ± 0 |  | 0 ± 0 | 0 ± 0 |
| RLS_010 | 10 | South Stirling | WA | 2018 | 26.2 ± 0.07 |  | 0 ± 0 | 0 ± 0 | 0 ± 0 | 0 ± 0 |  | 0 ± 0 | 0 ± 0 |
| RLS_011 | 10 | South Stirling | WA | 2018 | 22 ± 0.06 |  | 0 ± 0 | 0 ± 0 | 0 ± 0 | 0 ± 0 |  | 0 ± 0 | 0 ± 0 |
| RLS_012 | 10 | South Stirling | WA | 2018 | 26.2 ± 0.02 |  | 0 ± 0 | 0 ± 0 | 0 ± 0 | 0 ± 0 |  | 0 ± 0 | 0 ± 0 |
| RLS_013 | 10 | South Stirling | WA | 2018 | 23 ± 0 |  | 0 ± 0 | 0 ± 0 | 0 ± 0 | 0 ± 0 |  | 0 ± 0 | 0 ± 0 |
| RLS_014 | 10 | South Stirling | WA | 2018 | 30.1 ± 0 |  | 0 ± 0 | 0 ± 0 | 0 ± 0 | 0 ± 0 |  | 0 ± 0 | 0 ± 0 |
| RLS_015 | 10 | South Stirling | WA | 2018 | 26.8 ± 0.02 |  | 0 ± 0 | 0 ± 0 | 0 ± 0 | 0 ± 0 |  | 0 ± 0 | 0 ± 0 |
| RLS_016 | 10 | South Stirling | WA | 2018 | 26.2 ± 0.06 |  | 0 ± 0 | 0 ± 0 | 0 ± 0 | 0 ± 0 |  | 0 ± 0 | 0 ± 0 |
| RLS_017 | 10 | South Stirling | WA | 2018 | 23.9 ± 0.04 |  | 0 ± 0 | 0 ± 0 | 0 ± 0 | 0 ± 0 |  | 0 ± 0 | 0 ± 0 |
| RLS_018 | 10 | South Stirling | WA | 2018 | 26.5 ± 0.02 |  | 0 ± 0 | 0 ± 0 | 0 ± 0 | 0 ± 0 |  | 0 ± 0 | 0 ± 0 |
| RLS_019 | 10 | South Stirling | WA | 2018 | 23.2 ± 0.05 |  | 0 ± 0 | 0 ± 0 | 0 ± 0 | 0 ± 0 |  | 0 ± 0 | 0 ± 0 |
| RLS_020 | 10 | South Stirling | WA | 2018 | 26.5 ± 0.02 |  | 0 ± 0 | 0 ± 0 | 0 ± 0 | 0 ± 0 |  | 0 ± 0 | 0 ± 0 |
| RLS_021 | 10 | South Stirling | WA | 2018 | 25.7 ± 0.14 |  | 0 ± 0 | 0 ± 0 | 0 ± 0 | 0 ± 0 |  | 0 ± 0 | 0 ± 0 |
| RLS_022 | 10 | South Stirling | WA | 2018 | 24.7 ± 0.01 |  | 0 ± 0 | 0 ± 0 | 0 ± 0 | 0 ± 0 |  | 0 ± 0 | 0 ± 0 |
| RLS_023 | 10 | South Stirling | WA | 2018 | 28.5 ± 0.03 |  | 0 ± 0 | 0 ± 0 | 0 ± 0 | 0 ± 0 |  | 0 ± 0 | 0 ± 0 |
| RLS_024 | 10 | South Stirling | WA | 2018 | 24.2 ± 0.04 |  | 0 ± 0 | 0 ± 0 | 0 ± 0 | 0 ± 0 |  | 0 ± 0 | 0 ± 0 |
| RLS_025 | 10 | South Stirling | WA | 2018 | 30.1 ± 0.02 |  | 0 ± 0 | 0 ± 0 | 0 ± 0 | 0 ± 0 |  | 0 ± 0 | 0 ± 0 |
| RLS_026 | 10 | South Stirling | WA | 2018 | 34.4 ± 0.36 |  | 0 ± 0 | 0 ± 0 | 0 ± 0 | 0 ± 0 |  | 0 ± 0 | 0 ± 0 |
| RLS_027 | 10 | South Stirling | WA | 2018 | 23.3 ± 0.02 |  | 0 ± 0 | 0 ± 0 | 0 ± 0 | 0 ± 0 |  | 0 ± 0 | 0 ± 0 |
| RLS_028 | 10 | South Stirling | WA | 2018 | 28.4 ± 0.04 |  | 0 ± 0 | 0 ± 0 | 0 ± 0 | 0 ± 0 |  | 0 ± 0 | 0 ± 0 |
| RLS_029 | 10 | South Stirling | WA | 2018 | 27.6 ± 0.09 |  | 0 ± 0 | 0 ± 0 | 0 ± 0 | 0 ± 0 |  | 0 ± 0 | 0 ± 0 |
| RLS_030 | 10 | South Stirling | WA | 2018 | 30.1 ± 0.01 |  | 0 ± 0 | 0 ± 0 | 0 ± 0 | 0 ± 0 |  | 0 ± 0 | 0 ± 0 |
| RLS_141 | 10 | South Stirling | WA | 2018 | 23.5 ± 0.2 |  | 0 ± 0 | 0 ± 0 | 0 ± 0 | 0 ± 0 |  | 0 ± 0 | 0 ± 0 |
| RLS_142 | 10 | South Stirling | WA | 2018 | 22.7 ± 0.07 |  | 0 ± 0 | 0 ± 0 | 0 ± 0 | 0 ± 0 |  | 0 ± 0 | 0 ± 0 |
| RLS_143 | 10 | South Stirling | WA | 2018 | 24.2 ± 0.04 |  | 0 ± 0 | 0 ± 0 | 0 ± 0 | 0 ± 0 |  | 0 ± 0 | 0 ± 0 |
| RLS_144 | 10 | South Stirling | WA | 2018 | 22.9 ± 0.09 |  | 0 ± 0 | 0 ± 0 | 0 ± 0 | 0 ± 0 |  | 0 ± 0 | 0 ± 0 |
| RLS_145 | 10 | South Stirling | WA | 2018 | 23.5 ± 0.1 |  | 0 ± 0 | 0 ± 0 | 0 ± 0 | 0 ± 0 |  | 0 ± 0 | 0 ± 0 |
| RLS_146 | 10 | South Stirling | WA | 2018 | 24.3 ± 0.03 |  | 0 ± 0 | 0 ± 0 | 0 ± 0 | 0 ± 0 |  | 0 ± 0 | 0 ± 0 |
| RLS_147 | 10 | South Stirling | WA | 2018 | 22.3 ± 0.06 |  | 0 ± 0 | 0 ± 0 | 0 ± 0 | 0 ± 0 |  | 0 ± 0 | 0 ± 0 |
| RLS_148 | 10 | South Stirling | WA | 2018 | 22.9 ± 0.02 |  | 0 ± 0 | 0 ± 0 | 0 ± 0 | 0 ± 0 |  | 0 ± 0 | 0 ± 0 |
| RLS_149 | 10 | South Stirling | WA | 2018 | 22.8 ± 0.05 |  | 0 ± 0 | 0 ± 0 | 0 ± 0 | 0 ± 0 |  | 0 ± 0 | 0 ± 0 |
| RLS_150 | 10 | South Stirling | WA | 2018 | 22.9 ± 0.11 |  | 0 ± 0 | 0 ± 0 | 0 ± 0 | 0 ± 0 |  | 0 ± 0 | 0 ± 0 |
| RLS_151 | 10 | South Stirling | WA | 2018 | 24.4 ± 0.07 |  | 0 ± 0 | 0 ± 0 | 0 ± 0 | 0 ± 0 |  | 0 ± 0 | 0 ± 0 |
| RLS_152 | 10 | South Stirling | WA | 2018 | 23.9 ± 0.11 |  | 0 ± 0 | 0 ± 0 | 0 ± 0 | 0 ± 0 |  | 0 ± 0 | 0 ± 0 |
| RLS_153 | 10 | South Stirling | WA | 2018 | 24.6 ± 0.06 |  | 0 ± 0 | 0 ± 0 | 0 ± 0 | 0 ± 0 |  | 0 ± 0 | 0 ± 0 |
| RLS_154 | 10 | South Stirling | WA | 2018 | 24.7 ± 0.12 |  | 0 ± 0 | 0 ± 0 | 0 ± 0 | 0 ± 0 |  | 0 ± 0 | 0 ± 0 |
| RLS_155 | 10 | South Stirling | WA | 2018 | 23.9 ± 0.05 |  | 0 ± 0 | 0 ± 0 | 0 ± 0 | 0 ± 0 |  | 0 ± 0 | 0 ± 0 |
| RLS_156 | 10 | South Stirling | WA | 2018 | 24.2 ± 0.09 |  | 0 ± 0 | 0 ± 0 | 0 ± 0 | 0 ± 0 |  | 0 ± 0 | 0 ± 0 |
| RLS_157 | 10 | South Stirling | WA | 2018 | 23.5 ± 0.15 |  | 0 ± 0 | 0 ± 0 | 0 ± 0 | 0 ± 0 |  | 0 ± 0 | 0 ± 0 |
| RLS_158 | 10 | South Stirling | WA | 2018 | 24 ± 0.18 |  | 0 ± 0 | 0 ± 0 | 0 ± 0 | 0 ± 0 |  | 0 ± 0 | 0 ± 0 |
| RLS_159 | 10 | South Stirling | WA | 2018 | 24.3 ± 0.16 |  | 0 ± 0 | 0 ± 0 | 0 ± 0 | 0 ± 0 |  | 0 ± 0 | 0 ± 0 |
| RLS_160 | 10 | South Stirling | WA | 2018 | 23.2 ± 0.07 |  | 0 ± 0 | 0 ± 0 | 0 ± 0 | 0 ± 0 |  | 0 ± 0 | 0 ± 0 |
| RLS_161 | 10 | South Stirling | WA | 2018 | 24.7 ± 0.24 |  | 0 ± 0 | 0 ± 0 | 0 ± 0 | 0 ± 0 |  | 0 ± 0 | 0 ± 0 |
| RLS_162 | 10 | South Stirling | WA | 2018 | 25.2 ± 0.06 |  | 0 ± 0 | 0 ± 0 | 0 ± 0 | 0 ± 0 |  | 0 ± 0 | 0 ± 0 |
| RLS_163 | 10 | South Stirling | WA | 2018 | 24 ± 0.08 |  | 0 ± 0 | 0 ± 0 | 0 ± 0 | 0 ± 0 |  | 0 ± 0 | 0 ± 0 |
| RLS_164 | 10 | South Stirling | WA | 2018 | 24 ± 0 |  | 0 ± 0 | 0 ± 0 | 0 ± 0 | 0 ± 0 |  | 0 ± 0 | 0 ± 0 |
| RLS_165 | 10 | South Stirling | WA | 2018 | 23.7 ± 0.05 |  | 0 ± 0 | 0 ± 0 | 0 ± 0 | 0 ± 0 |  | 0 ± 0 | 0 ± 0 |
| RLS_166 | 10 | South Stirling | WA | 2018 | 23.9 ± 0.11 |  | 0 ± 0 | 0 ± 0 | 0 ± 0 | 0 ± 0 |  | 0 ± 0 | 0 ± 0 |
| RLS_167 | 10 | South Stirling | WA | 2018 | 23.1 ± 0.08 |  | 0 ± 0 | 0 ± 0 | 0 ± 0 | 0 ± 0 |  | 0 ± 0 | 0 ± 0 |
| RLS_168 | 10 | South Stirling | WA | 2018 | 24.3 ± 0.1 |  | 0 ± 0 | 0 ± 0 | 0 ± 0 | 0 ± 0 |  | 0 ± 0 | 0 ± 0 |
| RLS_169 | 10 | South Stirling | WA | 2018 | 23.4 ± 0.21 |  | 0 ± 0 | 0 ± 0 | 0 ± 0 | 0 ± 0 |  | 0 ± 0 | 0 ± 0 |
| RLS_170 | 10 | South Stirling | WA | 2018 | 23.6 ± 0.08 |  | 0 ± 0 | 0 ± 0 | 0 ± 0 | 0 ± 0 |  | 0 ± 0 | 0 ± 0 |
| RLS_111 | — | Conmurra | SA | 2020 | 19.1 ± 0.03 |  | 25.5 ± 0.06 | 211 ± 0.01 | 26.8 ± 0.03 | 119 ± 0 |  | 22.9 ± 0.04 | 62 ± 0 |
| RLS_112 | — | Conmurra | SA | 2020 | 18.6 ± 0 |  | 30 ± 0.02 | 12 ± 0 | 30.9 ± 0.13 | 8 ± 0 |  | 26.6 ± 0.04 | 6 ± 0 |
| RLS_113 | — | Conmurra | SA | 2020 | 18.8 ± 0.07 |  | 27.4 ± 0.45 | 66 ± 0.02 | 28.7 ± 0.02 | 33 ± 0 |  | 24.1 ± 0.01 | 30 ± 0 |
| RLS_114 | — | Conmurra | SA | 2020 | 18.5 ± 0.07 |  | 23.9 ± 0.06 | 569 ± 0.02 | 25.1 ± 0.03 | 367 ± 0.01 |  | 20.3 ± 0.08 | 328 ± 0.02 |
| RLS_115 | — | Conmurra | SA | 2020 | 18.2 ± 0 |  | 27.5 ± 0.09 | 60 ± 0 | 29.6 ± 0.06 | 18 ± 0 |  | 25.2 ± 0.02 | 15 ± 0 |
| RLS_116 | — | Conmurra | SA | 2020 | 18.3 ± 0.01 |  | 25.5 ± 0.12 | 215 ± 0.02 | 27.1 ± 0.07 | 96 ± 0 |  | 23 ± 0.13 | 61 ± 0 |
| RLS_117 | — | Conmurra | SA | 2020 | 18.3 ± 0.01 |  | 30.2 ± 0.21 | 11 ± 0 | 30.8 ± 0.04 | 8 ± 0 |  | 26.6 ± 0.13 | 6 ± 0 |
| RLS_118 | — | Conmurra | SA | 2020 | 18.3 ± 0.05 |  | 25.4 ± 0.05 | 219 ± 0.01 | 26.8 ± 0.05 | 119 ± 0 |  | 22.6 ± 0.06 | 76 ± 0 |
| RLS_119 | — | Conmurra | SA | 2020 | 18.2 ± 0 |  | 26.1 ± 0.09 | 144 ± 0.01 | 28.1 ± 0.11 | 48 ± 0 |  | 23.6 ± 0.02 | 40 ± 0 |
| RLS_120 | — | Conmurra | SA | 2020 | 18.4 ± 0.04 |  | 24.7 ± 0.03 | 342 ± 0.01 | 26.1 ± 0.03 | 190 ± 0 |  | 21.8 ± 0.04 | 129 ± 0 |
| RLS_111 | 10 | Conmurra | SA | 2020 | 22.4 ± 0.01 |  | 29.1 ± 0.08 | 22 ± 0 | 30.3 ± 0.08 | 11 ± 0 |  | 26.3 ± 0 | 7 ± 0 |
| RLS_112 | 10 | Conmurra | SA | 2020 | 22 ± 0.07 |  | 33.3 ± 0.77 | 2 ± 0 | 34.8 ± 0.01 | 1 ± 0 |  | 30.3 ± 0.28 | 1 ± 0 |
| RLS_113 | 10 | Conmurra | SA | 2020 | 22.1 ± 0.02 |  | 30.3 ± 0.36 | 11 ± 0 | 31.9 ± 0.14 | 4 ± 0 |  | 27.8 ± 0.15 | 3 ± 0 |
| RLS_114 | 10 | Conmurra | SA | 2020 | 21.7 ± 0.14 |  | 27.1 ± 0.02 | 77 ± 0 | 28.3 ± 0.07 | 44 ± 0 |  | 23.9 ± 0.04 | 34 ± 0 |
| RLS_115 | 10 | Conmurra | SA | 2020 | 21.7 ± 0 |  | 32.3 ± 0.17 | 3 ± 0 | 33.3 ± 0.14 | 1 ± 0 |  | 29.1 ± 0 | 1 ± 0 |
| RLS_116 | 10 | Conmurra | SA | 2020 | 21.7 ± 0.01 |  | 29.3 ± 0.06 | 20 ± 0 | 30.7 ± 0.11 | 9 ± 0 |  | 26.9 ± 0 | 5 ± 0 |
| RLS_117 | 10 | Conmurra | SA | 2020 | 21.9 ± 0.12 |  | 34.9 ± 0.76 | 1 ± 0 | 35.2 ± 0.5 | 0 ± 0 |  | 30.8 ± 0.05 | 0 ± 0 |
| RLS_118 | 10 | Conmurra | SA | 2020 | 21.6 ± 0.03 |  | 28.7 ± 0.18 | 29 ± 0 | 30.1 ± 0.16 | 13 ± 0 |  | 26.3 ± 0.04 | 7 ± 0 |
| RLS_119 | 10 | Conmurra | SA | 2020 | 21.6 ± 0.02 |  | 30 ± 0.14 | 13 ± 0 | 31.1 ± 0.25 | 6 ± 0 |  | 27.3 ± 0.13 | 4 ± 0 |
| RLS_120 | 10 | Conmurra | SA | 2020 | 21.9 ± 0.06 |  | 28.3 ± 0.1 | 37 ± 0 | 29.6 ± 0.02 | 18 ± 0 |  | 25.6 ± 0.03 | 12 ± 0 |
| RLS_131 | — | Hagley | Tas | 2020 | 18.4 ± 0.08 |  | 26.2 ± 0.35 | 142 ± 0.03 | 28 ± 0.09 | 54 ± 0 |  | 23.2 ± 0.05 | 53 ± 0 |
| RLS_132 | — | Hagley | Tas | 2020 | 18.6 ± 0.05 |  | 22.3 ± 0.03 | 1520 ± 0 | 23.1 ± 0.04 | 1430 ± 0.04 |  | 18.4 ± 0.12 | 1050 ± 0.08 |
| RLS_133 | — | Hagley | Tas | 2020 | 18.9 ± 0.03 |  | 24.1 ± 0 | 505 ± 0 | 25.3 ± 0.01 | 334 ± 0 |  | 20.7 ± 0.04 | 255 ± 0.01 |
| RLS_134 | — | Hagley | Tas | 2020 | 18.9 ± 0.06 |  | 25.7 ± 0.17 | 188 ± 0.02 | 26.2 ± 0.05 | 178 ± 0.01 |  | 21.8 ± 0.03 | 129 ± 0 |
| RLS_135 | — | Hagley | Tas | 2020 | 18.6 ± 0.07 |  | 25 ± 0 | 276 ± 0 | 26.4 ± 0.01 | 159 ± 0 |  | 22 ± 0.07 | 115 ± 0 |
| RLS_136 | — | Hagley | Tas | 2020 | 18.7 ± 0.02 |  | 22.8 ± 0.03 | 1080 ± 0.02 | 23.7 ± 0 | 924 ± 0 |  | 19.1 ± 0.12 | 674 ± 0.05 |
| RLS_137 | — | Hagley | Tas | 2020 | 18.7 ± 0.01 |  | 23.6 ± 0.09 | 692 ± 0.04 | 24.6 ± 0.03 | 524 ± 0.01 |  | 20 ± 0.03 | 394 ± 0.01 |
| RLS_138 | — | Hagley | Tas | 2020 | 19 ± 0.09 |  | 21.4 ± 0.05 | 2650 ± 0.08 | 22.3 ± 0.05 | 2450 ± 0.09 |  | 16.7 ± 0.87 | 3560 ± 1.77 |
| RLS_139 | — | Hagley | Tas | 2020 | 20.3 ± 0.03 |  | 19.9 ± 0.02 | 6800 ± 0.07 | 20.5 ± 0.01 | 8250 ± 0.08 |  | 15.6 ± 0 | 5990 ± 0.01 |
| RLS_140 | — | Hagley | Tas | 2020 | 18.7 ± 0 |  | 0 ± 0 | 0 ± 0 | 25.5 ± 0.3 | 291 ± 0.06 |  | 19.8 ± 0.16 | 448 ± 0.04 |
| RLS_131 | 10 | Hagley | Tas | 2020 | 22.2 ± 0.12 |  | 30.1 ± 0.45 | 12 ± 0 | 31.4 ± 0.19 | 5 ± 0 |  | 27.3 ± 0.17 | 4 ± 0 |
| RLS_132 | 10 | Hagley | Tas | 2020 | 22.1 ± 0.01 |  | 25.6 ± 0.02 | 191 ± 0 | 26.5 ± 0.01 | 142 ± 0 |  | 22.2 ± 0.04 | 99 ± 0 |
| RLS_133 | 10 | Hagley | Tas | 2020 | 22 ± 0.03 |  | 27.3 ± 0.14 | 68 ± 0.01 | 28.5 ± 0.01 | 37 ± 0 |  | 24 ± 0.03 | 31 ± 0 |
| RLS_134 | 10 | Hagley | Tas | 2020 | 22.4 ± 0.05 |  | 28.4 ± 0.04 | 35 ± 0 | 29.7 ± 0.02 | 17 ± 0 |  | 25.6 ± 0.03 | 12 ± 0 |
| RLS_135 | 10 | Hagley | Tas | 2020 | 21.8 ± 0.04 |  | 28.6 ± 0.28 | 32 ± 0.01 | 29.7 ± 0.01 | 17 ± 0 |  | 24.7 ± 1.05 | 25 ± 0.01 |
| RLS_136 | 10 | Hagley | Tas | 2020 | 21.9 ± 0.05 |  | 26.2 ± 0.06 | 136 ± 0 | 27 ± 0.01 | 100 ± 0 |  | 22.8 ± 0.06 | 70 ± 0 |
| RLS_137 | 10 | Hagley | Tas | 2020 | 22.2 ± 0.1 |  | 27 ± 0.07 | 80 ± 0 | 28.1 ± 0.08 | 49 ± 0 |  | 20 ± 0.03 | 394 ± 0.01 |
| RLS_138 | 10 | Hagley | Tas | 2020 | 22.3 ± 0.05 |  | 24.9 ± 0.05 | 297 ± 0.01 | 25.8 ± 0.08 | 236 ± 0.01 |  | 21.2 ± 0.1 | 185 ± 0.01 |
| RLS_139 | 10 | Hagley | Tas | 2020 | 23.8 ± 0.02 |  | 23.3 ± 0 | 810 ± 0 | 24.1 ± 0.01 | 735 ± 0 |  | 19.3 ± 0.12 | 592 ± 0.04 |
| RLS_140 | 10 | Hagley | Tas | 2020 | 21.9 ± 0.11 |  | 26.8 ± 0.03 | 91 ± 0 | 27.8 ± 0.08 | 61 ± 0 |  | 23.4 ± 0.17 | 47 ± 0.01 |
| RLS_121 | — | Gnarwarre | Vic | 2020 | 0 ± 0 |  | 0 ± 0 | 0 ± 0 | 0 ± 0 | 0 ± 0 |  | 0 ± 0 | 0 ± 0 |
| RLS_122 | — | Gnarwarre | Vic | 2020 | 0 ± 0 |  | 0 ± 0 | 0 ± 0 | 0 ± 0 | 0 ± 0 |  | 0 ± 0 | 0 ± 0 |
| RLS_123 | — | Gnarwarre | Vic | 2020 | 0 ± 0 |  | 0 ± 0 | 0 ± 0 | 0 ± 0 | 0 ± 0 |  | 0 ± 0 | 0 ± 0 |
| RLS_124 | — | Gnarwarre | Vic | 2020 | 0 ± 0 |  | 0 ± 0 | 0 ± 0 | 0 ± 0 | 0 ± 0 |  | 0 ± 0 | 0 ± 0 |
| RLS_125 | — | Gnarwarre | Vic | 2020 | 0 ± 0 |  | 0 ± 0 | 0 ± 0 | 0 ± 0 | 0 ± 0 |  | 0 ± 0 | 0 ± 0 |
| RLS_126 | — | Gnarwarre | Vic | 2020 | 0 ± 0 |  | 0 ± 0 | 0 ± 0 | 0 ± 0 | 0 ± 0 |  | 0 ± 0 | 0 ± 0 |
| RLS_127 | — | Gnarwarre | Vic | 2020 | 0 ± 0 |  | 25.5 ± 0.36 | 515 ± 0.11 | 22.4 ± 0.09 | 4120 ± 0.25 |  | 19.8 ± 0.59 | 460 ± 0.16 |
| RLS_128 | — | Gnarwarre | Vic | 2020 | 0 ± 0 |  | 0 ± 0 | 0 ± 0 | 0 ± 0 | 0 ± 0 |  | 0 ± 0 | 0 ± 0 |
| RLS_129 | — | Gnarwarre | Vic | 2020 | 0 ± 0 |  | 0 ± 0 | 0 ± 0 | 0 ± 0 | 0 ± 0 |  | 0 ± 0 | 0 ± 0 |
| RLS_130 | — | Gnarwarre | Vic | 2020 | 0 ± 0 |  | 0 ± 0 | 0 ± 0 | 0 ± 0 | 0 ± 0 |  | 0 ± 0 | 0 ± 0 |
| RLS_121 | 10 | Gnarwarre | Vic | 2020 | 30.4 ± 0.03 |  | 23.6 ± 0.07 | 849 ± 0.03 | 24.2 ± 0.01 | 797 ± 0 |  | 19.4 ± 0.04 | 724 ± 0.01 |
| RLS_122 | 10 | Gnarwarre | Vic | 2020 | 31.3 ± 0.23 |  | 23.7 ± 0.1 | 825 ± 0.04 | 24 ± 0 | 919 ± 0 |  | 19.1 ± 0.05 | 849 ± 0.02 |
| RLS_123 | 10 | Gnarwarre | Vic | 2020 | 29.2 ± 0.11 |  | 22.5 ± 0.03 | 1540 ± 0.03 | 23.1 ± 0 | 1550 ± 0 |  | 18.1 ± 0.01 | 1480 ± 0 |
| RLS_124 | 10 | Gnarwarre | Vic | 2020 | 29.5 ± 0.05 |  | 22.4 ± 0.02 | 1660 ± 0.02 | 22.6 ± 0.01 | 2160 ± 0.01 |  | 17.9 ± 0.13 | 1640 ± 0.11 |
| RLS_125 | 10 | Gnarwarre | Vic | 2020 | 28.8 ± 0.05 |  | 23.3 ± 0 | 1020 ± 0 | 23.9 ± 0.01 | 960 ± 0 |  | 19.2 ± 0.01 | 845 ± 0 |
| RLS_126 | 10 | Gnarwarre | Vic | 2020 | 30.2 ± 0.12 |  | 23.1 ± 0.03 | 1100 ± 0.02 | 23.7 ± 0.08 | 1130 ± 0.05 |  | 18.8 ± 0.12 | 1040 ± 0.06 |
| RLS_127 | 10 | Gnarwarre | Vic | 2020 | 29.6 ± 0.07 |  | 23.1 ± 0.04 | 1100 ± 0.02 | 23.8 ± 0.06 | 1060 ± 0.04 |  | 18.8 ± 0.03 | 1010 ± 0.01 |
| RLS_128 | 10 | Gnarwarre | Vic | 2020 | 29.1 ± 0.1 |  | 23.1 ± 0.1 | 1110 ± 0.06 | 23.6 ± 0.02 | 1150 ± 0.01 |  | 18.7 ± 0.07 | 1090 ± 0.04 |
| RLS_129 | 10 | Gnarwarre | Vic | 2020 | 28.8 ± 0.08 |  | 22.1 ± 0.17 | 1900 ± 0.16 | 22.7 ± 0.08 | 2020 ± 0.1 |  | 17.9 ± 0.12 | 1630 ± 0.1 |
| RLS_130 | 10 | Gnarwarre | Vic | 2020 | 29 ± 0.06 |  | 22.5 ± 0.01 | 1560 ± 0 | 23.1 ± 0.03 | 1620 ± 0.03 |  | 18 ± 0.01 | 1540 ± 0.01 |
| RLS_031 | — | South Stirling | WA | 2020 | 0 ± 0 |  | 0 ± 0 | 0 ± 0 | 0 ± 0 | 0 ± 0 |  | 0 ± 0 | 0 ± 0 |
| RLS_032 | — | South Stirling | WA | 2020 | 0 ± 0 |  | 0 ± 0 | 0 ± 0 | 0 ± 0 | 0 ± 0 |  | 0 ± 0 | 0 ± 0 |
| RLS_033 | — | South Stirling | WA | 2020 | 0 ± 0 |  | 0 ± 0 | 0 ± 0 | 0 ± 0 | 0 ± 0 |  | 0 ± 0 | 0 ± 0 |
| RLS_034 | — | South Stirling | WA | 2020 | 0 ± 0 |  | 0 ± 0 | 0 ± 0 | 0 ± 0 | 0 ± 0 |  | 0 ± 0 | 0 ± 0 |
| RLS_035 | — | South Stirling | WA | 2020 | 0 ± 0 |  | 0 ± 0 | 0 ± 0 | 0 ± 0 | 0 ± 0 |  | 0 ± 0 | 0 ± 0 |
| RLS_036 | — | South Stirling | WA | 2020 | 0 ± 0 |  | 0 ± 0 | 0 ± 0 | 0 ± 0 | 0 ± 0 |  | 0 ± 0 | 0 ± 0 |
| RLS_037 | — | South Stirling | WA | 2020 | 0 ± 0 |  | 0 ± 0 | 0 ± 0 | 0 ± 0 | 0 ± 0 |  | 0 ± 0 | 0 ± 0 |
| RLS_038 | — | South Stirling | WA | 2020 | 0 ± 0 |  | 0 ± 0 | 0 ± 0 | 0 ± 0 | 0 ± 0 |  | 0 ± 0 | 0 ± 0 |
| RLS_039 | — | South Stirling | WA | 2020 | 0 ± 0 |  | 0 ± 0 | 0 ± 0 | 0 ± 0 | 0 ± 0 |  | 0 ± 0 | 0 ± 0 |
| RLS_040 | — | South Stirling | WA | 2020 | 0 ± 0 |  | 0 ± 0 | 0 ± 0 | 0 ± 0 | 0 ± 0 |  | 0 ± 0 | 0 ± 0 |
| RLS_041 | — | South Stirling | WA | 2020 | 0 ± 0 |  | 0 ± 0 | 0 ± 0 | 0 ± 0 | 0 ± 0 |  | 0 ± 0 | 0 ± 0 |
| RLS_042 | — | South Stirling | WA | 2020 | 0 ± 0 |  | 0 ± 0 | 0 ± 0 | 0 ± 0 | 0 ± 0 |  | 0 ± 0 | 0 ± 0 |
| RLS_043 | — | South Stirling | WA | 2020 | 0 ± 0 |  | 0 ± 0 | 0 ± 0 | 0 ± 0 | 0 ± 0 |  | 0 ± 0 | 0 ± 0 |
| RLS_044 | — | South Stirling | WA | 2020 | 0 ± 0 |  | 0 ± 0 | 0 ± 0 | 0 ± 0 | 0 ± 0 |  | 0 ± 0 | 0 ± 0 |
| RLS_045 | — | South Stirling | WA | 2020 | 0 ± 0 |  | 0 ± 0 | 0 ± 0 | 0 ± 0 | 0 ± 0 |  | 0 ± 0 | 0 ± 0 |
| RLS_046 | — | South Stirling | WA | 2020 | 0 ± 0 |  | 0 ± 0 | 0 ± 0 | 0 ± 0 | 0 ± 0 |  | 0 ± 0 | 0 ± 0 |
| RLS_047 | — | South Stirling | WA | 2020 | 0 ± 0 |  | 0 ± 0 | 0 ± 0 | 0 ± 0 | 0 ± 0 |  | 0 ± 0 | 0 ± 0 |
| RLS_048 | — | South Stirling | WA | 2020 | 0 ± 0 |  | 0 ± 0 | 0 ± 0 | 0 ± 0 | 0 ± 0 |  | 0 ± 0 | 0 ± 0 |
| RLS_049 | — | South Stirling | WA | 2020 | 0 ± 0 |  | 0 ± 0 | 0 ± 0 | 0 ± 0 | 0 ± 0 |  | 0 ± 0 | 0 ± 0 |
| RLS_050 | — | South Stirling | WA | 2020 | 29.2 ± 0.21 |  | 0 ± 0 | 0 ± 0 | 0 ± 0 | 0 ± 0 |  | 0 ± 0 | 0 ± 0 |
| RLS_051 | — | South Stirling | WA | 2020 | 0 ± 0 |  | 0 ± 0 | 0 ± 0 | 0 ± 0 | 0 ± 0 |  | 0 ± 0 | 0 ± 0 |
| RLS_052 | — | South Stirling | WA | 2020 | 0 ± 0 |  | 0 ± 0 | 0 ± 0 | 0 ± 0 | 0 ± 0 |  | 0 ± 0 | 0 ± 0 |
| RLS_053 | — | South Stirling | WA | 2020 | 29.2 ± 0.85 |  | 0 ± 0 | 0 ± 0 | 0 ± 0 | 0 ± 0 |  | 0 ± 0 | 0 ± 0 |
| RLS_054 | — | South Stirling | WA | 2020 | 0 ± 0 |  | 0 ± 0 | 0 ± 0 | 0 ± 0 | 0 ± 0 |  | 0 ± 0 | 0 ± 0 |
| RLS_055 | — | South Stirling | WA | 2020 | 0 ± 0 |  | 0 ± 0 | 0 ± 0 | 0 ± 0 | 0 ± 0 |  | 0 ± 0 | 0 ± 0 |
| RLS_056 | — | South Stirling | WA | 2020 | 26.3 ± 0.03 |  | 0 ± 0 | 0 ± 0 | 0 ± 0 | 0 ± 0 |  | 0 ± 0 | 0 ± 0 |
| RLS_057 | — | South Stirling | WA | 2020 | 0 ± 0 |  | 0 ± 0 | 0 ± 0 | 0 ± 0 | 0 ± 0 |  | 0 ± 0 | 0 ± 0 |
| RLS_058 | — | South Stirling | WA | 2020 | 0 ± 0 |  | 0 ± 0 | 0 ± 0 | 0 ± 0 | 0 ± 0 |  | 0 ± 0 | 0 ± 0 |
| RLS_059 | — | South Stirling | WA | 2020 | 0 ± 0 |  | 0 ± 0 | 0 ± 0 | 0 ± 0 | 0 ± 0 |  | 0 ± 0 | 0 ± 0 |
| RLS_060 | — | South Stirling | WA | 2020 | 0 ± 0 |  | 0 ± 0 | 0 ± 0 | 0 ± 0 | 0 ± 0 |  | 0 ± 0 | 0 ± 0 |
| RLS_061 | — | South Stirling | WA | 2020 | 0 ± 0 |  | 0 ± 0 | 0 ± 0 | 0 ± 0 | 0 ± 0 |  | 0 ± 0 | 0 ± 0 |
| RLS_062 | — | South Stirling | WA | 2020 | 0 ± 0 |  | 0 ± 0 | 0 ± 0 | 0 ± 0 | 0 ± 0 |  | 0 ± 0 | 0 ± 0 |
| RLS_063 | — | South Stirling | WA | 2020 | 0 ± 0 |  | 0 ± 0 | 0 ± 0 | 0 ± 0 | 0 ± 0 |  | 0 ± 0 | 0 ± 0 |
| RLS_064 | — | South Stirling | WA | 2020 | 0 ± 0 |  | 0 ± 0 | 0 ± 0 | 0 ± 0 | 0 ± 0 |  | 0 ± 0 | 0 ± 0 |
| RLS_065 | — | South Stirling | WA | 2020 | 27.8 ± 0.04 |  | 0 ± 0 | 0 ± 0 | 0 ± 0 | 0 ± 0 |  | 0 ± 0 | 0 ± 0 |
| RLS_066 | — | South Stirling | WA | 2020 | 0 ± 0 |  | 0 ± 0 | 0 ± 0 | 0 ± 0 | 0 ± 0 |  | 0 ± 0 | 0 ± 0 |
| RLS_067 | — | South Stirling | WA | 2020 | 0 ± 0 |  | 0 ± 0 | 0 ± 0 | 0 ± 0 | 0 ± 0 |  | 0 ± 0 | 0 ± 0 |
| RLS_068 | — | South Stirling | WA | 2020 | 0 ± 0 |  | 0 ± 0 | 0 ± 0 | 0 ± 0 | 0 ± 0 |  | 0 ± 0 | 0 ± 0 |
| RLS_069 | — | South Stirling | WA | 2020 | 0 ± 0 |  | 0 ± 0 | 0 ± 0 | 0 ± 0 | 0 ± 0 |  | 0 ± 0 | 0 ± 0 |
| RLS_070 | — | South Stirling | WA | 2020 | 0 ± 0 |  | 0 ± 0 | 0 ± 0 | 0 ± 0 | 0 ± 0 |  | 0 ± 0 | 0 ± 0 |
| RLS_071 | — | South Stirling | WA | 2020 | 0 ± 0 |  | 0 ± 0 | 0 ± 0 | 0 ± 0 | 0 ± 0 |  | 0 ± 0 | 0 ± 0 |
| RLS_072 | — | South Stirling | WA | 2020 | 30.5 ± 0.15 |  | 0 ± 0 | 0 ± 0 | 0 ± 0 | 0 ± 0 |  | 0 ± 0 | 0 ± 0 |
| RLS_073 | — | South Stirling | WA | 2020 | 0 ± 0 |  | 0 ± 0 | 0 ± 0 | 0 ± 0 | 0 ± 0 |  | 0 ± 0 | 0 ± 0 |
| RLS_074 | — | South Stirling | WA | 2020 | 0 ± 0 |  | 0 ± 0 | 0 ± 0 | 0 ± 0 | 0 ± 0 |  | 0 ± 0 | 0 ± 0 |
| RLS_075 | — | South Stirling | WA | 2020 | 28.6 ± 0.02 |  | 0 ± 0 | 0 ± 0 | 0 ± 0 | 0 ± 0 |  | 30.6 ± 0.09 | 1 ± 0 |
| RLS_076 | — | South Stirling | WA | 2020 | 0 ± 0 |  | 0 ± 0 | 0 ± 0 | 0 ± 0 | 0 ± 0 |  | 0 ± 0 | 0 ± 0 |
| RLS_077 | — | South Stirling | WA | 2020 | 0 ± 0 |  | 0 ± 0 | 0 ± 0 | 0 ± 0 | 0 ± 0 |  | 0 ± 0 | 0 ± 0 |
| RLS_078 | — | South Stirling | WA | 2020 | 0 ± 0 |  | 0 ± 0 | 0 ± 0 | 0 ± 0 | 0 ± 0 |  | 0 ± 0 | 0 ± 0 |
| RLS_079 | — | South Stirling | WA | 2020 | 0 ± 0 |  | 0 ± 0 | 0 ± 0 | 0 ± 0 | 0 ± 0 |  | 0 ± 0 | 0 ± 0 |
| RLS_080 | — | South Stirling | WA | 2020 | 0 ± 0 |  | 0 ± 0 | 0 ± 0 | 0 ± 0 | 0 ± 0 |  | 0 ± 0 | 0 ± 0 |
| RLS_081 | — | South Stirling | WA | 2020 | 0 ± 0 |  | 0 ± 0 | 0 ± 0 | 0 ± 0 | 0 ± 0 |  | 0 ± 0 | 0 ± 0 |
| RLS_082 | — | South Stirling | WA | 2020 | 0 ± 0 |  | 0 ± 0 | 0 ± 0 | 0 ± 0 | 0 ± 0 |  | 0 ± 0 | 0 ± 0 |
| RLS_083 | — | South Stirling | WA | 2020 | 0 ± 0 |  | 0 ± 0 | 0 ± 0 | 0 ± 0 | 0 ± 0 |  | 0 ± 0 | 0 ± 0 |
| RLS_084 | — | South Stirling | WA | 2020 | 0 ± 0 |  | 0 ± 0 | 0 ± 0 | 0 ± 0 | 0 ± 0 |  | 0 ± 0 | 0 ± 0 |
| RLS_085 | — | South Stirling | WA | 2020 | 0 ± 0 |  | 0 ± 0 | 0 ± 0 | 0 ± 0 | 0 ± 0 |  | 0 ± 0 | 0 ± 0 |
| RLS_086 | — | South Stirling | WA | 2020 | 0 ± 0 |  | 0 ± 0 | 0 ± 0 | 0 ± 0 | 0 ± 0 |  | 0 ± 0 | 0 ± 0 |
| RLS_087 | — | South Stirling | WA | 2020 | 0 ± 0 |  | 0 ± 0 | 0 ± 0 | 0 ± 0 | 0 ± 0 |  | 0 ± 0 | 0 ± 0 |
| RLS_088 | — | South Stirling | WA | 2020 | 30.5 ± 0.18 |  | 0 ± 0 | 0 ± 0 | 0 ± 0 | 0 ± 0 |  | 0 ± 0 | 0 ± 0 |
| RLS_089 | — | South Stirling | WA | 2020 | 28.5 ± 0.14 |  | 0 ± 0 | 0 ± 0 | 0 ± 0 | 0 ± 0 |  | 0 ± 0 | 0 ± 0 |
| RLS_090 | — | South Stirling | WA | 2020 | 0 ± 0 |  | 0 ± 0 | 0 ± 0 | 0 ± 0 | 0 ± 0 |  | 0 ± 0 | 0 ± 0 |
| RLS_091 | — | South Stirling | WA | 2020 | 29.6 ± 0.12 |  | 0 ± 0 | 0 ± 0 | 0 ± 0 | 0 ± 0 |  | 0 ± 0 | 0 ± 0 |
| RLS_092 | — | South Stirling | WA | 2020 | 0 ± 0 |  | 0 ± 0 | 0 ± 0 | 0 ± 0 | 0 ± 0 |  | 0 ± 0 | 0 ± 0 |
| RLS_093 | — | South Stirling | WA | 2020 | 0 ± 0 |  | 0 ± 0 | 0 ± 0 | 0 ± 0 | 0 ± 0 |  | 0 ± 0 | 0 ± 0 |
| RLS_094 | — | South Stirling | WA | 2020 | 29.4 ± 0.25 |  | 0 ± 0 | 0 ± 0 | 0 ± 0 | 0 ± 0 |  | 27.7 ± 0.17 | 3 ± 0 |
| RLS_095 | — | South Stirling | WA | 2020 | 0 ± 0 |  | 0 ± 0 | 0 ± 0 | 0 ± 0 | 0 ± 0 |  | 0 ± 0 | 0 ± 0 |
| RLS_096 | — | South Stirling | WA | 2020 | 0 ± 0 |  | 0 ± 0 | 0 ± 0 | 0 ± 0 | 0 ± 0 |  | 0 ± 0 | 0 ± 0 |
| RLS_097 | — | South Stirling | WA | 2020 | 0 ± 0 |  | 0 ± 0 | 0 ± 0 | 0 ± 0 | 0 ± 0 |  | 0 ± 0 | 0 ± 0 |
| RLS_098 | — | South Stirling | WA | 2020 | 0 ± 0 |  | 0 ± 0 | 0 ± 0 | 0 ± 0 | 0 ± 0 |  | 0 ± 0 | 0 ± 0 |
| RLS_099 | — | South Stirling | WA | 2020 | 0 ± 0 |  | 0 ± 0 | 0 ± 0 | 0 ± 0 | 0 ± 0 |  | 0 ± 0 | 0 ± 0 |
| RLS_100 | — | South Stirling | WA | 2020 | 29.5 ± 0.25 |  | 0 ± 0 | 0 ± 0 | 0 ± 0 | 0 ± 0 |  | 0 ± 0 | 0 ± 0 |
| RLS_101 | — | South Stirling | WA | 2020 | 0 ± 0 |  | 0 ± 0 | 0 ± 0 | 0 ± 0 | 0 ± 0 |  | 0 ± 0 | 0 ± 0 |
| RLS_102 | — | South Stirling | WA | 2020 | 28.1 ± 0.08 |  | 0 ± 0 | 0 ± 0 | 0 ± 0 | 0 ± 0 |  | 0 ± 0 | 0 ± 0 |
| RLS_103 | — | South Stirling | WA | 2020 | 0 ± 0 |  | 0 ± 0 | 0 ± 0 | 0 ± 0 | 0 ± 0 |  | 0 ± 0 | 0 ± 0 |
| RLS_104 | — | South Stirling | WA | 2020 | 0 ± 0 |  | 0 ± 0 | 0 ± 0 | 0 ± 0 | 0 ± 0 |  | 0 ± 0 | 0 ± 0 |
| RLS_105 | — | South Stirling | WA | 2020 | 0 ± 0 |  | 0 ± 0 | 0 ± 0 | 0 ± 0 | 0 ± 0 |  | 0 ± 0 | 0 ± 0 |
| RLS_106 | — | South Stirling | WA | 2020 | 0 ± 0 |  | 0 ± 0 | 0 ± 0 | 0 ± 0 | 0 ± 0 |  | 0 ± 0 | 0 ± 0 |
| RLS_107 | — | South Stirling | WA | 2020 | 0 ± 0 |  | 0 ± 0 | 0 ± 0 | 0 ± 0 | 0 ± 0 |  | 0 ± 0 | 0 ± 0 |
| RLS_108 | — | South Stirling | WA | 2020 | 28.8 ± 0.05 |  | 0 ± 0 | 0 ± 0 | 0 ± 0 | 0 ± 0 |  | 0 ± 0 | 0 ± 0 |
| RLS_109 | — | South Stirling | WA | 2020 | 29.2 ± 0.19 |  | 0 ± 0 | 0 ± 0 | 0 ± 0 | 0 ± 0 |  | 0 ± 0 | 0 ± 0 |
| RLS_110 | — | South Stirling | WA | 2020 | 0 ± 0 |  | 0 ± 0 | 0 ± 0 | 0 ± 0 | 0 ± 0 |  | 0 ± 0 | 0 ± 0 |
| RLS_031 | 10 | South Stirling | WA | 2020 | 29.5 ± 0.11 |  | 0 ± 0 | 0 ± 0 | 0 ± 0 | 0 ± 0 |  | 0 ± 0 | 0 ± 0 |
| RLS_032 | 10 | South Stirling | WA | 2020 | 31.3 ± 0.01 |  | 0 ± 0 | 0 ± 0 | 0 ± 0 | 0 ± 0 |  | 0 ± 0 | 0 ± 0 |
| RLS_033 | 10 | South Stirling | WA | 2020 | 30.3 ± 0.08 |  | 0 ± 0 | 0 ± 0 | 0 ± 0 | 0 ± 0 |  | 0 ± 0 | 0 ± 0 |
| RLS_034 | 10 | South Stirling | WA | 2020 | 0 ± 0 |  | 0 ± 0 | 0 ± 0 | 0 ± 0 | 0 ± 0 |  | 0 ± 0 | 0 ± 0 |
| RLS_035 | 10 | South Stirling | WA | 2020 | 30 ± 0.05 |  | 0 ± 0 | 0 ± 0 | 0 ± 0 | 0 ± 0 |  | 0 ± 0 | 0 ± 0 |
| RLS_036 | 10 | South Stirling | WA | 2020 | 31.3 ± 0.01 |  | 0 ± 0 | 0 ± 0 | 0 ± 0 | 0 ± 0 |  | 0 ± 0 | 0 ± 0 |
| RLS_037 | 10 | South Stirling | WA | 2020 | 30.8 ± 0.05 |  | 0 ± 0 | 0 ± 0 | 0 ± 0 | 0 ± 0 |  | 0 ± 0 | 0 ± 0 |
| RLS_038 | 10 | South Stirling | WA | 2020 | 31.2 ± 0.08 |  | 0 ± 0 | 0 ± 0 | 0 ± 0 | 0 ± 0 |  | 0 ± 0 | 0 ± 0 |
| RLS_039 | 10 | South Stirling | WA | 2020 | 0 ± 0 |  | 0 ± 0 | 0 ± 0 | 0 ± 0 | 0 ± 0 |  | 33.8 ± 0.12 | 0 ± 0 |
| RLS_040 | 10 | South Stirling | WA | 2020 | 31.1 ± 0.09 |  | 0 ± 0 | 0 ± 0 | 0 ± 0 | 0 ± 0 |  | 0 ± 0 | 0 ± 0 |
| RLS_041 | 10 | South Stirling | WA | 2020 | 31.2 ± 0.03 |  | 0 ± 0 | 0 ± 0 | 0 ± 0 | 0 ± 0 |  | 0 ± 0 | 0 ± 0 |
| RLS_042 | 10 | South Stirling | WA | 2020 | 30.7 ± 0.03 |  | 0 ± 0 | 0 ± 0 | 0 ± 0 | 0 ± 0 |  | 0 ± 0 | 0 ± 0 |
| RLS_043 | 10 | South Stirling | WA | 2020 | 0 ± 0 |  | 0 ± 0 | 0 ± 0 | 0 ± 0 | 0 ± 0 |  | 0 ± 0 | 0 ± 0 |
| RLS_044 | 10 | South Stirling | WA | 2020 | 31.2 ± 0.13 |  | 0 ± 0 | 0 ± 0 | 0 ± 0 | 0 ± 0 |  | 0 ± 0 | 0 ± 0 |
| RLS_045 | 10 | South Stirling | WA | 2020 | 30.6 ± 0.13 |  | 0 ± 0 | 0 ± 0 | 0 ± 0 | 0 ± 0 |  | 0 ± 0 | 0 ± 0 |
| RLS_046 | 10 | South Stirling | WA | 2020 | 30.8 ± 0.09 |  | 0 ± 0 | 0 ± 0 | 0 ± 0 | 0 ± 0 |  | 0 ± 0 | 0 ± 0 |
| RLS_047 | 10 | South Stirling | WA | 2020 | 30.4 ± 0.22 |  | 0 ± 0 | 0 ± 0 | 0 ± 0 | 0 ± 0 |  | 0 ± 0 | 0 ± 0 |
| RLS_048 | 10 | South Stirling | WA | 2020 | 31.6 ± 0.03 |  | 0 ± 0 | 0 ± 0 | 0 ± 0 | 0 ± 0 |  | 0 ± 0 | 0 ± 0 |
| RLS_049 | 10 | South Stirling | WA | 2020 | 0 ± 0 |  | 0 ± 0 | 0 ± 0 | 0 ± 0 | 0 ± 0 |  | 0 ± 0 | 0 ± 0 |
| RLS_050 | 10 | South Stirling | WA | 2020 | 30.5 ± 0.12 |  | 0 ± 0 | 0 ± 0 | 0 ± 0 | 0 ± 0 |  | 0 ± 0 | 0 ± 0 |
| RLS_051 | 10 | South Stirling | WA | 2020 | 30.9 ± 0.15 |  | 33.6 ± 0.34 | 5 ± 0 | 34.4 ± 0.34 | 2 ± 0 |  | 29.6 ± 0.09 | 3 ± 0 |
| RLS_052 | 10 | South Stirling | WA | 2020 | 31.2 ± 0.04 |  | 0 ± 0 | 0 ± 0 | 0 ± 0 | 0 ± 0 |  | 0 ± 0 | 0 ± 0 |
| RLS_053 | 10 | South Stirling | WA | 2020 | 30.2 ± 0.05 |  | 0 ± 0 | 0 ± 0 | 0 ± 0 | 0 ± 0 |  | 0 ± 0 | 0 ± 0 |
| RLS_054 | 10 | South Stirling | WA | 2020 | 31.1 ± 0.06 |  | 0 ± 0 | 0 ± 0 | 0 ± 0 | 0 ± 0 |  | 0 ± 0 | 0 ± 0 |
| RLS_055 | 10 | South Stirling | WA | 2020 | 30.6 ± 0.19 |  | 0 ± 0 | 0 ± 0 | 0 ± 0 | 0 ± 0 |  | 0 ± 0 | 0 ± 0 |
| RLS_056 | 10 | South Stirling | WA | 2020 | 28.7 ± 0.07 |  | 0 ± 0 | 0 ± 0 | 0 ± 0 | 0 ± 0 |  | 0 ± 0 | 0 ± 0 |
| RLS_057 | 10 | South Stirling | WA | 2020 | 30.4 ± 0.1 |  | 0 ± 0 | 0 ± 0 | 0 ± 0 | 0 ± 0 |  | 0 ± 0 | 0 ± 0 |
| RLS_058 | 10 | South Stirling | WA | 2020 | 0 ± 0 |  | 0 ± 0 | 0 ± 0 | 0 ± 0 | 0 ± 0 |  | 34.3 ± 0.24 | 0 ± 0 |
| RLS_059 | 10 | South Stirling | WA | 2020 | 0 ± 0 |  | 0 ± 0 | 0 ± 0 | 0 ± 0 | 0 ± 0 |  | 0 ± 0 | 0 ± 0 |
| RLS_060 | 10 | South Stirling | WA | 2020 | 31.2 ± 0.24 |  | 0 ± 0 | 0 ± 0 | 0 ± 0 | 0 ± 0 |  | 0 ± 0 | 0 ± 0 |
| RLS_061 | 10 | South Stirling | WA | 2020 | 31.2 ± 0.15 |  | 0 ± 0 | 0 ± 0 | 0 ± 0 | 0 ± 0 |  | 0 ± 0 | 0 ± 0 |
| RLS_062 | 10 | South Stirling | WA | 2020 | 0 ± 0 |  | 0 ± 0 | 0 ± 0 | 0 ± 0 | 0 ± 0 |  | 0 ± 0 | 0 ± 0 |
| RLS_063 | 10 | South Stirling | WA | 2020 | 32.6 ± 0.08 |  | 0 ± 0 | 0 ± 0 | 0 ± 0 | 0 ± 0 |  | 0 ± 0 | 0 ± 0 |
| RLS_064 | 10 | South Stirling | WA | 2020 | 29.6 ± 0.05 |  | 0 ± 0 | 0 ± 0 | 0 ± 0 | 0 ± 0 |  | 0 ± 0 | 0 ± 0 |
| RLS_065 | 10 | South Stirling | WA | 2020 | 30.1 ± 0.02 |  | 0 ± 0 | 0 ± 0 | 0 ± 0 | 0 ± 0 |  | 34.7 ± 0.11 | 0 ± 0 |
| RLS_066 | 10 | South Stirling | WA | 2020 | 0 ± 0 |  | 0 ± 0 | 0 ± 0 | 0 ± 0 | 0 ± 0 |  | 0 ± 0 | 0 ± 0 |
| RLS_067 | 10 | South Stirling | WA | 2020 | 29.8 ± 0.01 |  | 0 ± 0 | 0 ± 0 | 0 ± 0 | 0 ± 0 |  | 0 ± 0 | 0 ± 0 |
| RLS_068 | 10 | South Stirling | WA | 2020 | 30.8 ± 0.05 |  | 0 ± 0 | 0 ± 0 | 0 ± 0 | 0 ± 0 |  | 0 ± 0 | 0 ± 0 |
| RLS_069 | 10 | South Stirling | WA | 2020 | 0 ± 0 |  | 0 ± 0 | 0 ± 0 | 0 ± 0 | 0 ± 0 |  | 0 ± 0 | 0 ± 0 |
| RLS_070 | 10 | South Stirling | WA | 2020 | 30.4 ± 0.16 |  | 0 ± 0 | 0 ± 0 | 0 ± 0 | 0 ± 0 |  | 0 ± 0 | 0 ± 0 |
| RLS_071 | 10 | South Stirling | WA | 2020 | 31.3 ± 0.28 |  | 0 ± 0 | 0 ± 0 | 0 ± 0 | 0 ± 0 |  | 0 ± 0 | 0 ± 0 |
| RLS_072 | 10 | South Stirling | WA | 2020 | 0 ± 0 |  | 0 ± 0 | 0 ± 0 | 0 ± 0 | 0 ± 0 |  | 0 ± 0 | 0 ± 0 |
| RLS_073 | 10 | South Stirling | WA | 2020 | 29.9 ± 0.15 |  | 0 ± 0 | 0 ± 0 | 0 ± 0 | 0 ± 0 |  | 0 ± 0 | 0 ± 0 |
| RLS_074 | 10 | South Stirling | WA | 2020 | 30.2 ± 0.01 |  | 0 ± 0 | 0 ± 0 | 0 ± 0 | 0 ± 0 |  | 0 ± 0 | 0 ± 0 |
| RLS_075 | 10 | South Stirling | WA | 2020 | 0 ± 0 |  | 0 ± 0 | 0 ± 0 | 0 ± 0 | 0 ± 0 |  | 0 ± 0 | 0 ± 0 |
| RLS_076 | 10 | South Stirling | WA | 2020 | 32 ± 0.05 |  | 0 ± 0 | 0 ± 0 | 0 ± 0 | 0 ± 0 |  | 0 ± 0 | 0 ± 0 |
| RLS_077 | 10 | South Stirling | WA | 2020 | 0 ± 0 |  | 0 ± 0 | 0 ± 0 | 0 ± 0 | 0 ± 0 |  | 0 ± 0 | 0 ± 0 |
| RLS_078 | 10 | South Stirling | WA | 2020 | 31.4 ± 0.21 |  | 0 ± 0 | 0 ± 0 | 0 ± 0 | 0 ± 0 |  | 0 ± 0 | 0 ± 0 |
| RLS_079 | 10 | South Stirling | WA | 2020 | 30 ± 0.14 |  | 0 ± 0 | 0 ± 0 | 0 ± 0 | 0 ± 0 |  | 0 ± 0 | 0 ± 0 |
| RLS_080 | 10 | South Stirling | WA | 2020 | 30.5 ± 0.08 |  | 0 ± 0 | 0 ± 0 | 0 ± 0 | 0 ± 0 |  | 0 ± 0 | 0 ± 0 |
| RLS_081 | 10 | South Stirling | WA | 2020 | 31.5 ± 0.11 |  | 0 ± 0 | 0 ± 0 | 0 ± 0 | 0 ± 0 |  | 0 ± 0 | 0 ± 0 |
| RLS_082 | 10 | South Stirling | WA | 2020 | 27.1 ± 0.04 |  | 0 ± 0 | 0 ± 0 | 0 ± 0 | 0 ± 0 |  | 0 ± 0 | 0 ± 0 |
| RLS_083 | 10 | South Stirling | WA | 2020 | 0 ± 0 |  | 0 ± 0 | 0 ± 0 | 0 ± 0 | 0 ± 0 |  | 0 ± 0 | 0 ± 0 |
| RLS_084 | 10 | South Stirling | WA | 2020 | 30.8 ± 0.01 |  | 0 ± 0 | 0 ± 0 | 0 ± 0 | 0 ± 0 |  | 34.1 ± 0.24 | 0 ± 0 |
| RLS_085 | 10 | South Stirling | WA | 2020 | 30.6 ± 0.15 |  | 0 ± 0 | 0 ± 0 | 0 ± 0 | 0 ± 0 |  | 0 ± 0 | 0 ± 0 |
| RLS_086 | 10 | South Stirling | WA | 2020 | 30.8 ± 0.02 |  | 0 ± 0 | 0 ± 0 | 0 ± 0 | 0 ± 0 |  | 0 ± 0 | 0 ± 0 |
| RLS_087 | 10 | South Stirling | WA | 2020 | 0 ± 0 |  | 0 ± 0 | 0 ± 0 | 0 ± 0 | 0 ± 0 |  | 0 ± 0 | 0 ± 0 |
| RLS_088 | 10 | South Stirling | WA | 2020 | 29.3 ± 0.09 |  | 0 ± 0 | 0 ± 0 | 0 ± 0 | 0 ± 0 |  | 0 ± 0 | 0 ± 0 |
| RLS_089 | 10 | South Stirling | WA | 2020 | 31.7 ± 0.06 |  | 0 ± 0 | 0 ± 0 | 0 ± 0 | 0 ± 0 |  | 34.6 ± 0.13 | 0 ± 0 |
| RLS_090 | 10 | South Stirling | WA | 2020 | 31.3 ± 0.08 |  | 0 ± 0 | 0 ± 0 | 0 ± 0 | 0 ± 0 |  | 0 ± 0 | 0 ± 0 |
| RLS_091 | 10 | South Stirling | WA | 2020 | 32.3 ± 0.17 |  | 0 ± 0 | 0 ± 0 | 0 ± 0 | 0 ± 0 |  | 0 ± 0 | 0 ± 0 |
| RLS_092 | 10 | South Stirling | WA | 2020 | 30.6 ± 0.07 |  | 0 ± 0 | 0 ± 0 | 0 ± 0 | 0 ± 0 |  | 0 ± 0 | 0 ± 0 |
| RLS_093 | 10 | South Stirling | WA | 2020 | 30.7 ± 0.01 |  | 0 ± 0 | 0 ± 0 | 0 ± 0 | 0 ± 0 |  | 0 ± 0 | 0 ± 0 |
| RLS_094 | 10 | South Stirling | WA | 2020 | 0 ± 0 |  | 0 ± 0 | 0 ± 0 | 0 ± 0 | 0 ± 0 |  | 31.7 ± 0.31 | 1 ± 0 |
| RLS_095 | 10 | South Stirling | WA | 2020 | 31.2 ± 0.27 |  | 0 ± 0 | 0 ± 0 | 0 ± 0 | 0 ± 0 |  | 0 ± 0 | 0 ± 0 |
| RLS_096 | 10 | South Stirling | WA | 2020 | 31.1 ± 0.03 |  | 0 ± 0 | 0 ± 0 | 0 ± 0 | 0 ± 0 |  | 0 ± 0 | 0 ± 0 |
| RLS_097 | 10 | South Stirling | WA | 2020 | 31.4 ± 0.06 |  | 0 ± 0 | 0 ± 0 | 0 ± 0 | 0 ± 0 |  | 0 ± 0 | 0 ± 0 |
| RLS_098 | 10 | South Stirling | WA | 2020 | 31.3 ± 0.11 |  | 0 ± 0 | 0 ± 0 | 0 ± 0 | 0 ± 0 |  | 0 ± 0 | 0 ± 0 |
| RLS_099 | 10 | South Stirling | WA | 2020 | 29.8 ± 0.16 |  | 0 ± 0 | 0 ± 0 | 0 ± 0 | 0 ± 0 |  | 0 ± 0 | 0 ± 0 |
| RLS_100 | 10 | South Stirling | WA | 2020 | 31.5 ± 0.19 |  | 0 ± 0 | 0 ± 0 | 0 ± 0 | 0 ± 0 |  | 0 ± 0 | 0 ± 0 |
| RLS_101 | 10 | South Stirling | WA | 2020 | 30.7 ± 0.01 |  | 0 ± 0 | 0 ± 0 | 0 ± 0 | 0 ± 0 |  | 0 ± 0 | 0 ± 0 |
| RLS_102 | 10 | South Stirling | WA | 2020 | 30.9 ± 0.06 |  | 0 ± 0 | 0 ± 0 | 0 ± 0 | 0 ± 0 |  | 0 ± 0 | 0 ± 0 |
| RLS_103 | 10 | South Stirling | WA | 2020 | 30.8 ± 0.13 |  | 0 ± 0 | 0 ± 0 | 0 ± 0 | 0 ± 0 |  | 0 ± 0 | 0 ± 0 |
| RLS_104 | 10 | South Stirling | WA | 2020 | 0 ± 0 |  | 0 ± 0 | 0 ± 0 | 0 ± 0 | 0 ± 0 |  | 0 ± 0 | 0 ± 0 |
| RLS_105 | 10 | South Stirling | WA | 2020 | 30.1 ± 0.07 |  | 0 ± 0 | 0 ± 0 | 0 ± 0 | 0 ± 0 |  | 33.1 ± 0.1 | 1 ± 0 |
| RLS_106 | 10 | South Stirling | WA | 2020 | 30.1 ± 0.03 |  | 0 ± 0 | 0 ± 0 | 0 ± 0 | 0 ± 0 |  | 0 ± 0 | 0 ± 0 |
| RLS_107 | 10 | South Stirling | WA | 2020 | 30.5 ± 0.14 |  | 0 ± 0 | 0 ± 0 | 0 ± 0 | 0 ± 0 |  | 0 ± 0 | 0 ± 0 |
| RLS_108 | 10 | South Stirling | WA | 2020 | 31.1 ± 0 |  | 0 ± 0 | 0 ± 0 | 0 ± 0 | 0 ± 0 |  | 33.3 ± 0.13 | 0 ± 0 |
| RLS_109 | 10 | South Stirling | WA | 2020 | 30.7 ± 0.18 |  | 0 ± 0 | 0 ± 0 | 0 ± 0 | 0 ± 0 |  | 31.1 ± 0.04 | 2 ± 0 |
| RLS_110 | 10 | South Stirling | WA | 2020 | 31.2 ± 0.1 |  | 0 ± 0 | 0 ± 0 | 0 ± 0 | 0 ± 0 |  | 30.1 ± 0.09 | 3 ± 0 |

^a^ A dash (—) indicates no dilution was performed. A ‘10’ indicates a ten-fold dilution was performed.
