## Supplementary material for "Detection of *Ramularia collo-cygni* from barley (*Hordeum vulgare*) in Australia using triplex quantitative and digital PCR": S5

**Supplementary Table S5.** Mean copy number values and standard errors (*n* = 2) for *Hordeum vulgare* and *Ramularia collo-cygni* DNA in leaf samples from New South Wales (NSW), South Australia (SA), Tasmania (Tas), Victoria (Vic) and Western Australia (WA) using *R. collo-cygni*-specific triplex droplet digital PCR.

| Sample | Region | State | Year | Dilution^a^ | *Hv*_116_*tef1-α* copies µL^-1^ | *Rcc*_139_*rpb2* copies µL^-1^ | *Rcc*_88_*tef1-α* copies µL^-1^ |
| --- | --- | --- | --- | --- | --- | --- | --- |
| 16MUL_001 | Mulwala | NSW | 2016 | — | 0.93 ± 0.055 | 0 ± 0 | 0 ± 0 |
| 16MUL_002 | Mulwala | NSW | 2016 | — | 0.28 ± 0.278 | 0.05 ± 0.048 | 0 ± 0 |
| 16MUL_004 | Mulwala | NSW | 2016 | — | 337.62 ± 29.6 | 0 ± 0 | 0 ± 0 |
| 16MUL_005 | Mulwala | NSW | 2016 | — | 1881.48 ± 91 | 1.15 ± 0.048 | 1.15 ± 0.048 |
| 16MUL_007 | Mulwala | NSW | 2016 | — | 2909.5 ± 181.381 | 0 ± 0 | 0 ± 0 |
| 16MUL_008 | Mulwala | NSW | 2016 | — | 32.32 ± 2.586 | 0 ± 0 | 0.07 ± 0.068 |
| 16DAI_001 | Dairy Plains | TAS | 2016 | — | 2659.59 ± 6.578 | 460.96 ± 4.07 | 512.79 ± 2.035 |
| 16DAI_002 | Dairy Plains | TAS | 2016 | — | 416.26 ± 5.036 | 67.02 ± 3.009 | 93.67 ± 1.059 |
| 16DAI_003 | Dairy Plains | TAS | 2016 | — | 6.05 ± 0.611 | 1.09 ± 0.023 | 1.78 ± 0.1 |
| 16DAI_004 | Dairy Plains | TAS | 2016 | — | 2328.89 ± 32.329 | 4.08 ± 0.575 | 4.17 ± 0.18 |
| 16DAI_005 | Dairy Plains | TAS | 2016 | — | 374.25 ± 14.735 | 80.37 ± 2.711 | 116.58 ± 1.689 |
| 16DAI_006 | Dairy Plains | TAS | 2016 | — | 2034.69 ± 15.739 | 273.76 ± 4.056 | 361.31 ± 3.576 |
| 16DAI_007 | Dairy Plains | TAS | 2016 | — | 518.51 ± 12.257 | 2.06 ± 0.019 | 2.98 ± 0.067 |
| 16DAI_008 | Dairy Plains | TAS | 2016 | — | 60.94 ± 0.288 | 37.24 ± 3.122 | 59.69 ± 2.756 |
| 16DAI_009 | Dairy Plains | TAS | 2016 | — | 116.86 ± 31.493 | 108.33 ± 9.861 | 130.07 ± 49.25 |
| 16DAI_001 | Dairy Plains | TAS | 2016 | 10 | 265.64 ± 0.663 | 43.96 ± 0.426 | 47.51 ± 0.724 |
| 16DAI_002 | Dairy Plains | TAS | 2016 | 10 | 41.55 ± 0.265 | 6.99 ± 0.543 | 9.64 ± 0.655 |
| 16DAI_003 | Dairy Plains | TAS | 2016 | 10 | 0.49 ± 0.175 | 0.07 ± 0.067 | 0.16 ± 0.037 |
| 16DAI_004 | Dairy Plains | TAS | 2016 | 10 | 272.62 ± 0.216 | 0.34 ± 0.141 | 0.45 ± 0.036 |
| 16DAI_005 | Dairy Plains | TAS | 2016 | 10 | 39.54 ± 0.322 | 8.95 ± 0.541 | 12.67 ± 0.796 |
| 16DAI_006 | Dairy Plains | TAS | 2016 | 10 | 233.47 ± 1.573 | 28.98 ± 1.023 | 35.26 ± 1.667 |
| 16DAI_007 | Dairy Plains | TAS | 2016 | 10 | 52.99 ± 0.553 | 0.14 ± 0.072 | 0.23 ± 0.042 |
| 16DAI_008 | Dairy Plains | TAS | 2016 | 10 | 7.48 ± 0.755 | 3.28 ± 0.009 | 6.14 ± 0.777 |
| 16DAI_009 | Dairy Plains | TAS | 2016 | 10 | 15.47 ± 0.57 | 12.35 ± 0.227 | 17.46 ± 1.07 |
| 16HAG_005 | Hagley | TAS | 2016 | — | 180.91 ± 2.716 | 0 ± 0 | 0.07 ± 0.071 |
| 16HAG_007 | Hagley | TAS | 2016 | — | 6716.2 ± 39.927 | 95.44 ± 3.509 | 190.13 ± 1.838 |
| 16HAG_008 | Hagley | TAS | 2016 | — | 3228.57 ± 16.504 | 57.47 ± 0.009 | 92.93 ± 0.376 |
| 16HAG_009 | Hagley | TAS | 2016 | — | 12370 ± 145.414 | 0.41 ± 0.002 | 0.88 ± 0.073 |
| 16HAG_013 | Hagley | TAS | 2016 | — | 527.29 ± 0.49 | 0.35 ± 0.089 | 1.26 ± 0.379 |
| 16HAG_014 | Hagley | TAS | 2016 | — | 1307.43 ± 3.143 | 101.52 ± 0.818 | 145.02 ± 1.764 |
| 16HAG_015 | Hagley | TAS | 2016 | — | 862.09 ± 16.348 | 4.06 ± 0.402 | 6.76 ± 0.121 |
| 16HAG_019 | Hagley | TAS | 2016 | — | 2652.35 ± 399.774 | 0.07 ± 0.002 | 14.37 ± 0.952 |
| 16HAG_020 | Hagley | TAS | 2016 | — | 7.82 ± 0.407 | 0 ± 0 | 0 ± 0 |
| 16HAG_026 | Hagley | TAS | 2016 | — | 99.01 ± 0.627 | 1.26 ± 0.296 | 2.06 ± 0.129 |
| 16HAG_005 | Hagley | TAS | 2016 | 10 | 22.07 ± 0.338 | 0 ± 0 | 0 ± 0 |
| 16HAG_007 | Hagley | TAS | 2016 | 10 | 819.34 ± 9.755 | 15.54 ± 0.732 | 19.13 ± 0.805 |
| 16HAG_008 | Hagley | TAS | 2016 | 10 | 335.65 ± 14.27 | 5.85 ± 0.171 | 8.58 ± 1.165 |
| 16HAG_009 | Hagley | TAS | 2016 | 10 | 1237.96 ± 14.541 | 0.12 ± 0.121 | 0.09 ± 0.091 |
| 16HAG_013 | Hagley | TAS | 2016 | 10 | 55.41 ± 0.862 | 0.03 ± 0.03 | 0.09 ± 0.088 |
| 16HAG_014 | Hagley | TAS | 2016 | 10 | 145.15 ± 2.062 | 11.56 ± 0.937 | 14.02 ± 1.195 |
| 16HAG_015 | Hagley | TAS | 2016 | 10 | 97.85 ± 1.875 | 0.43 ± 0.121 | 0.36 ± 0.118 |
| 16HAG_019 | Hagley | TAS | 2016 | 10 | 1289.51 ± 23.519 | 1.66 ± 0.58 | 1.66 ± 0.262 |
| 16HAG_020 | Hagley | TAS | 2016 | 10 | 0.92 ± 0.012 | 0 ± 0 | 0 ± 0 |
| 16HAG_026 | Hagley | TAS | 2016 | 10 | 9.82 ± 0.649 | 0.18 ± 0.062 | 0.32 ± 0.084 |
| RLS_001 | South Stirling | WA | 2018 | — | 517.14 ± 23.822 | 0 ± 0 | 0 ± 0 |
| RLS_002 | South Stirling | WA | 2018 | — | 853.97 ± 25.762 | 0 ± 0 | 0 ± 0 |
| RLS_003 | South Stirling | WA | 2018 | — | 40.36 ± 3.464 | 0 ± 0 | 0 ± 0 |
| RLS_004 | South Stirling | WA | 2018 | — | 73.44 ± 0.716 | 0 ± 0 | 0 ± 0 |
| RLS_005 | South Stirling | WA | 2018 | — | 100.47 ± 100.472 | 0.03 ± 0.032 | 0 ± 0 |
| RLS_006 | South Stirling | WA | 2018 | — | 7700.3 ± 309.106 | 0 ± 0 | 0 ± 0 |
| RLS_007 | South Stirling | WA | 2018 | — | 367.16 ± 9.169 | 0 ± 0 | 0.03 ± 0.033 |
| RLS_008 | South Stirling | WA | 2018 | — | 362.71 ± 1.847 | 0 ± 0 | 0 ± 0 |
| RLS_009 | South Stirling | WA | 2018 | — | 11730 ± 115.29 | 0 ± 0 | 0 ± 0 |
| RLS_010 | South Stirling | WA | 2018 | — | 1466.64 ± 3.232 | 0 ± 0 | 0 ± 0 |
| RLS_011 | South Stirling | WA | 2018 | — | 7534.37 ± 112.936 | 0 ± 0 | 0 ± 0 |
| RLS_012 | South Stirling | WA | 2018 | — | 1655.35 ± 22.081 | 0 ± 0 | 0 ± 0 |
| RLS_013 | South Stirling | WA | 2018 | — | 10600.52 ± 771.906 | 0 ± 0 | 0 ± 0 |
| RLS_014 | South Stirling | WA | 2018 | — | 130.66 ± 0.439 | 0 ± 0 | 0 ± 0 |
| RLS_015 | South Stirling | WA | 2018 | — | 1277.89 ± 15.063 | 0 ± 0 | 0 ± 0 |
| RLS_016 | South Stirling | WA | 2018 | — | 1649.95 ± 23.414 | 0 ± 0 | 0 ± 0 |
| RLS_017 | South Stirling | WA | 2018 | — | 7180.37 ± 192.807 | 0 ± 0 | 0 ± 0 |
| RLS_018 | South Stirling | WA | 2018 | — | 1283.68 ± 5.154 | 0.04 ± 0.036 | 0 ± 0 |
| RLS_019 | South Stirling | WA | 2018 | — | 9420.62 ± 711.453 | 0.16 ± 0.159 | 0 ± 0 |
| RLS_020 | South Stirling | WA | 2018 | — | 884.32 ± 176.986 | 0 ± 0 | 0 ± 0 |
| RLS_021 | South Stirling | WA | 2018 | — | 2675.82 ± 12.651 | 0 ± 0 | 0 ± 0 |
| RLS_022 | South Stirling | WA | 2018 | — | 2837.21 ± 27.109 | 0 ± 0 | 0 ± 0 |
| RLS_023 | South Stirling | WA | 2018 | — | 365.82 ± 2.402 | 0 ± 0 | 0 ± 0 |
| RLS_024 | South Stirling | WA | 2018 | — | 2446.55 ± 1404.436 | 0.04 ± 0.035 | 0 ± 0 |
| RLS_025 | South Stirling | WA | 2018 | — | 111.79 ± 6.316 | 0 ± 0 | 0 ± 0 |
| RLS_026 | South Stirling | WA | 2018 | — | 6.93 ± 0.4 | 0 ± 0 | 0 ± 0 |
| RLS_027 | South Stirling | WA | 2018 | — | 4929.45 ± 1863.109 | 0 ± 0 | 0 ± 0 |
| RLS_028 | South Stirling | WA | 2018 | — | 322.03 ± 5.559 | 0 ± 0 | 0 ± 0 |
| RLS_029 | South Stirling | WA | 2018 | — | 546.85 ± 100.041 | 0 ± 0 | 0 ± 0 |
| RLS_030 | South Stirling | WA | 2018 | — | 63.27 ± 63.274 | 0.09 ± 0.087 | 0.04 ± 0.044 |
| RLS_141 | South Stirling | WA | 2018 | — | 3786.67 ± 47.479 | 0.05 ± 0.05 | 0 ± 0 |
| RLS_142 | South Stirling | WA | 2018 | — | 3537.45 ± 82.477 | 0 ± 0 | 0 ± 0 |
| RLS_143 | South Stirling | WA | 2018 | — | 1755.84 ± 24.099 | 0 ± 0 | 0.03 ± 0.034 |
| RLS_144 | South Stirling | WA | 2018 | — | 4927.09 ± 6.845 | 0 ± 0 | 0 ± 0 |
| RLS_145 | South Stirling | WA | 2018 | — | 3209.52 ± 4.44 | 0.11 ± 0.035 | 0 ± 0 |
| RLS_146 | South Stirling | WA | 2018 | — | 1674.16 ± 21.898 | 0 ± 0 | 0.08 ± 0.076 |
| RLS_147 | South Stirling | WA | 2018 | — | 6736.02 ± 142.992 | 0 ± 0 | 0.04 ± 0.044 |
| RLS_148 | South Stirling | WA | 2018 | — | 4956.86 ± 200.601 | 0 ± 0 | 0.07 ± 0.069 |
| RLS_149 | South Stirling | WA | 2018 | — | 4468.38 ± 310.648 | 0 ± 0 | 0 ± 0 |
| RLS_150 | South Stirling | WA | 2018 | — | 3581.7 ± 406.236 | 0 ± 0 | 0 ± 0 |
| RLS_151 | South Stirling | WA | 2018 | — | 1691.18 ± 43.617 | 0 ± 0 | 0 ± 0 |
| RLS_152 | South Stirling | WA | 2018 | — | 2746.27 ± 53.944 | 0 ± 0 | 0 ± 0 |
| RLS_153 | South Stirling | WA | 2018 | — | 1725.28 ± 85.098 | 0 ± 0 | 0 ± 0 |
| RLS_154 | South Stirling | WA | 2018 | — | 1468.86 ± 27.404 | 0 ± 0 | 0 ± 0 |
| RLS_155 | South Stirling | WA | 2018 | — | 2589.98 ± 12.844 | 0 ± 0 | 0.03 ± 0.035 |
| RLS_156 | South Stirling | WA | 2018 | — | 2625.51 ± 35.14 | 0 ± 0 | 0 ± 0 |
| RLS_157 | South Stirling | WA | 2018 | — | 4055.1 ± 51.696 | 0.03 ± 0.034 | 0.03 ± 0.034 |
| RLS_158 | South Stirling | WA | 2018 | — | 2689.08 ± 116.801 | 0 ± 0 | 0.04 ± 0.036 |
| RLS_159 | South Stirling | WA | 2018 | — | 1961.34 ± 112.705 | 0 ± 0 | 0 ± 0 |
| RLS_160 | South Stirling | WA | 2018 | — | 5261.69 ± 149.027 | 0 ± 0 | 0.04 ± 0.04 |
| RLS_161 | South Stirling | WA | 2018 | — | 1849.76 ± 92.714 | 0 ± 0 | 0 ± 0 |
| RLS_162 | South Stirling | WA | 2018 | — | 1205.76 ± 38.987 | 0 ± 0 | 0.04 ± 0.037 |
| RLS_163 | South Stirling | WA | 2018 | — | 2606.5 ± 4.387 | 0 ± 0 | 0.04 ± 0.037 |
| RLS_164 | South Stirling | WA | 2018 | — | 3002.25 ± 39.903 | 0 ± 0 | 0 ± 0 |
| RLS_165 | South Stirling | WA | 2018 | — | 3524.76 ± 62.181 | 0 ± 0 | 0.03 ± 0.033 |
| RLS_166 | South Stirling | WA | 2018 | — | 2599.63 ± 16.447 | 0.03 ± 0.033 | 0 ± 0 |
| RLS_167 | South Stirling | WA | 2018 | — | 5428.42 ± 5.738 | 0 ± 0 | 0 ± 0 |
| RLS_168 | South Stirling | WA | 2018 | — | 2432.67 ± 44.726 | 0.03 ± 0.033 | 0 ± 0 |
| RLS_169 | South Stirling | WA | 2018 | — | 4561.01 ± 18.943 | 0 ± 0 | 0.04 ± 0.036 |
| RLS_170 | South Stirling | WA | 2018 | — | 3572.89 ± 38.957 | 0 ± 0 | 0 ± 0 |
| RLS_111 | Conmurra | SA | 2020 | — | 7441.82 ± 79.105 | 35.19 ± 5.357 | 38.61 ± 5.624 |
| RLS_112 | Conmurra | SA | 2020 | — | 11465.52 ± 9.023 | 3 ± 0.15 | 3.28 ± 0.06 |
| RLS_113 | Conmurra | SA | 2020 | — | 8803.54 ± 344.264 | 10.27 ± 0.281 | 13.6 ± 0.611 |
| RLS_114 | Conmurra | SA | 2020 | — | 11170 ± 69.527 | 151.3 ± 2.387 | 173.66 ± 2.706 |
| RLS_115 | Conmurra | SA | 2020 | — | 12610 ± 63.044 | 6.97 ± 0.404 | 8.1 ± 0.441 |
| RLS_116 | Conmurra | SA | 2020 | — | 6030 ± 634.615 | 34.24 ± 3 | 35.32 ± 1.094 |
| RLS_117 | Conmurra | SA | 2020 | — | 10710 ± 71.359 | 0.52 ± 0.005 | 2.37 ± 0.318 |
| RLS_118 | Conmurra | SA | 2020 | — | 9325.8 ± 1402.229 | 24.44 ± 0.248 | 47.02 ± 3.692 |
| RLS_119 | Conmurra | SA | 2020 | — | 7618.22 ± 39.664 | 0.15 ± 0.076 | 21.27 ± 1.83 |
| RLS_120 | Conmurra | SA | 2020 | — | 9568.32 ± 583.552 | 56.4 ± 2.562 | 69.64 ± 1.543 |
| RLS_111 | Conmurra | SA | 2020 | 10 | 625.1 ± 28.63 | 2.76 ± 0.596 | 3.03 ± 0.25 |
| RLS_112 | Conmurra | SA | 2020 | 10 | 1470.29 ± 431.725 | 0.75 ± 0.609 | 1.57 ± 1.223 |
| RLS_113 | Conmurra | SA | 2020 | 10 | 879.98 ± 16.464 | 1.25 ± 0.212 | 1.06 ± 0.039 |
| RLS_114 | Conmurra | SA | 2020 | 10 | 1117.17 ± 6.321 | 13.16 ± 0.011 | 15.76 ± 0.2 |
| RLS_115 | Conmurra | SA | 2020 | 10 | 1261.38 ± 5.731 | 0.53 ± 0.107 | 0.35 ± 0.142 |
| RLS_116 | Conmurra | SA | 2020 | 10 | 603.15 ± 603.147 | 1.67 ± 1.667 | 1.13 ± 1.133 |
| RLS_117 | Conmurra | SA | 2020 | 10 | 1071.85 ± 6.487 | 0.19 ± 0.056 | 0.47 ± 0.154 |
| RLS_118 | Conmurra | SA | 2020 | 10 | 1169.74 ± 6.691 | 2.93 ± 0.12 | 2.93 ± 0.12 |
| RLS_119 | Conmurra | SA | 2020 | 10 | 1351.91 ± 0.334 | 1.4 ± 0.292 | 1.45 ± 0.239 |
| RLS_120 | Conmurra | SA | 2020 | 10 | 1084 ± 24.354 | 4.14 ± 0.087 | 4.4 ± 0.351 |
| RLS_131 | Hagley | TAS | 2020 | — | 9286.45 ± 596.681 | 1.86 ± 0.268 | 19.96 ± 0.432 |
| RLS_132 | Hagley | TAS | 2020 | — | 9295.78 ± 360.95 | 426.8 ± 8.323 | 627.06 ± 22.661 |
| RLS_133 | Hagley | TAS | 2020 | — | 10498.78 ± 219.674 | 141.06 ± 6.868 | 161.75 ± 1.37 |
| RLS_134 | Hagley | TAS | 2020 | — | 9013.36 ± 280.008 | 65.59 ± 0.21 | 74.39 ± 0.388 |
| RLS_135 | Hagley | TAS | 2020 | — | 11860 ± 229.56 | 60.15 ± 0.079 | 60.89 ± 0.287 |
| RLS_136 | Hagley | TAS | 2020 | — | 9968.03 ± 363.393 | 377.51 ± 13.743 | 442.45 ± 1.489 |
| RLS_137 | Hagley | TAS | 2020 | — | 9559.94 ± 497.271 | 200.97 ± 0.032 | 233.11 ± 0.213 |
| RLS_138 | Hagley | TAS | 2020 | — | 6864.25 ± 499.626 | 511.67 ± 36.675 | 1133.47 ± 9.84 |
| RLS_139 | Hagley | TAS | 2020 | — | 2278.73 ± 49.722 | 838.85 ± 573.967 | 3565.62 ± 207.967 |
| RLS_140 | Hagley | TAS | 2020 | — | 8531.32 ± 345.4 | 41.76 ± 6.195 | 243.98 ± 0.451 |
| RLS_131 | Hagley | TAS | 2020 | 10 | 1000.92 ± 16.972 | 1.8 ± 0.002 | 1.78 ± 0.431 |
| RLS_132 | Hagley | TAS | 2020 | 10 | 1071.33 ± 12.313 | 62.08 ± 0.18 | 66.28 ± 1.754 |
| RLS_133 | Hagley | TAS | 2020 | 10 | 1053.55 ± 30.777 | 14.53 ± 0.774 | 14.99 ± 0.034 |
| RLS_134 | Hagley | TAS | 2020 | 10 | 896.64 ± 20.16 | 5.84 ± 0.397 | 6.22 ± 0.705 |
| RLS_135 | Hagley | TAS | 2020 | 10 | 1186.81 ± 21.544 | 7.2 ± 0.146 | 5.95 ± 0.258 |
| RLS_136 | Hagley | TAS | 2020 | 10 | 997.68 ± 20.442 | 32.73 ± 0.672 | 32.73 ± 0.966 |
| RLS_137 | Hagley | TAS | 2020 | 10 | 962.25 ± 3.656 | 22.48 ± 0.502 | 21.77 ± 1.414 |
| RLS_138 | Hagley | TAS | 2020 | 10 | 902.53 ± 6.219 | 104.59 ± 1.995 | 110.46 ± 0.852 |
| RLS_139 | Hagley | TAS | 2020 | 10 | 354.4 ± 3.649 | 371.51 ± 5.543 | 387.92 ± 6.014 |
| RLS_140 | Hagley | TAS | 2020 | 10 | 1150.23 ± 16.913 | 24.91 ± 0.329 | 25.6 ± 0.443 |
| RLS_121 | Gnarwarre | VIC | 2020 | — | 1.81 ± 0.071 | 62.87 ± 4.305 | 3114.72 ± 12.047 |
| RLS_122 | Gnarwarre | VIC | 2020 | — | 0.73 ± 0.004 | 21.66 ± 3.418 | 2657.38 ± 43.4 |
| RLS_123 | Gnarwarre | VIC | 2020 | — | 1.23 ± 0.296 | 4.48 ± 0.336 | 6486.34 ± 162.413 |
| RLS_124 | Gnarwarre | VIC | 2020 | — | 0.11 ± 0.031 | 0.73 ± 0.112 | 7711.02 ± 157.775 |
| RLS_125 | Gnarwarre | VIC | 2020 | — | 2.79 ± 0.252 | 37.45 ± 1.209 | 3770.98 ± 16.952 |
| RLS_126 | Gnarwarre | VIC | 2020 | — | 0.76 ± 0.231 | 19.29 ± 2.532 | 4302.34 ± 51.734 |
| RLS_127 | Gnarwarre | VIC | 2020 | — | 1.93 ± 0.386 | 18.37 ± 0.459 | 4744.25 ± 68.34 |
| RLS_128 | Gnarwarre | VIC | 2020 | — | 1.74 ± 0.742 | 19.53 ± 5.099 | 4454.3 ± 159.617 |
| RLS_129 | Gnarwarre | VIC | 2020 | — | 0.03 ± 0.034 | 0.63 ± 0.123 | 8299.79 ± 88.275 |
| RLS_130 | Gnarwarre | VIC | 2020 | — | 0.11 ± 0.044 | 9.47 ± 6.503 | 5269.54 ± 543.403 |
| RLS_121 | Gnarwarre | VIC | 2020 | 10 | 1.76 ± 0.077 | 229.38 ± 3.826 | 242.8 ± 4.461 |
| RLS_122 | Gnarwarre | VIC | 2020 | 10 | 1.4 ± 0.348 | 259.1 ± 3.894 | 265.1 ± 5.06 |
| RLS_123 | Gnarwarre | VIC | 2020 | 10 | 5.75 ± 0.313 | 599.5 ± 4.116 | 614.98 ± 6.552 |
| RLS_124 | Gnarwarre | VIC | 2020 | 10 | 9.49 ± 0.585 | 717.76 ± 42.676 | 741.74 ± 62.187 |
| RLS_125 | Gnarwarre | VIC | 2020 | 10 | 5.33 ± 0.556 | 349.09 ± 0.638 | 364.84 ± 14.035 |
| RLS_126 | Gnarwarre | VIC | 2020 | 10 | 2.73 ± 0.243 | 390.75 ± 1.824 | 394.72 ± 1.439 |
| RLS_127 | Gnarwarre | VIC | 2020 | 10 | 3.67 ± 0.001 | 405.94 ± 11.859 | 408 ± 15.132 |
| RLS_128 | Gnarwarre | VIC | 2020 | 10 | 6.3 ± 0.466 | 376.14 ± 23.072 | 378.18 ± 19.37 |
| RLS_129 | Gnarwarre | VIC | 2020 | 10 | 7.67 ± 0.057 | 774.31 ± 4.467 | 808.74 ± 9.022 |
| RLS_130 | Gnarwarre | VIC | 2020 | 10 | 5.44 ± 0.314 | 540.14 ± 21.191 | 544.79 ± 17.866 |
| RLS_031 | South Stirling | WA | 2020 | — | 2.28 ± 2.279 | 0 ± 0 | 0 ± 0 |
| RLS_032 | South Stirling | WA | 2020 | — | 1.4 ± 0.299 | 0 ± 0 | 0 ± 0 |
| RLS_033 | South Stirling | WA | 2020 | — | 5.1 ± 1.142 | 0 ± 0 | 0 ± 0 |
| RLS_034 | South Stirling | WA | 2020 | — | 0.48 ± 0.053 | 0.13 ± 0.133 | 0 ± 0 |
| RLS_035 | South Stirling | WA | 2020 | — | 10.63 ± 10.625 | 0 ± 0 | 0 ± 0 |
| RLS_036 | South Stirling | WA | 2020 | — | 4.07 ± 0.017 | 0 ± 0 | 0 ± 0 |
| RLS_037 | South Stirling | WA | 2020 | — | 8.45 ± 0.289 | 0 ± 0 | 0 ± 0 |
| RLS_038 | South Stirling | WA | 2020 | — | 2.23 ± 0.142 | 0 ± 0 | 0 ± 0 |
| RLS_039 | South Stirling | WA | 2020 | — | 11.3 ± 1.181 | 0 ± 0 | 0 ± 0 |
| RLS_040 | South Stirling | WA | 2020 | — | 2.12 ± 0.003 | 0 ± 0 | 0 ± 0 |
| RLS_041 | South Stirling | WA | 2020 | — | 3.09 ± 0.761 | 0 ± 0 | 0 ± 0 |
| RLS_042 | South Stirling | WA | 2020 | — | 8.16 ± 0.7 | 0 ± 0 | 0 ± 0 |
| RLS_043 | South Stirling | WA | 2020 | — | 2.79 ± 0.811 | 0.09 ± 0.086 | 0 ± 0 |
| RLS_044 | South Stirling | WA | 2020 | — | 15.11 ± 1.072 | 0 ± 0 | 0 ± 0 |
| RLS_045 | South Stirling | WA | 2020 | — | 0.76 ± 0.043 | 0 ± 0 | 0 ± 0 |
| RLS_046 | South Stirling | WA | 2020 | — | 1.66 ± 0.413 | 0 ± 0 | 0 ± 0 |
| RLS_047 | South Stirling | WA | 2020 | — | 17.04 ± 0.535 | 0.04 ± 0.037 | 0 ± 0 |
| RLS_048 | South Stirling | WA | 2020 | — | 2.03 ± 0.449 | 0 ± 0 | 0 ± 0 |
| RLS_049 | South Stirling | WA | 2020 | — | 0.44 ± 0.002 | 0.04 ± 0.037 | 0 ± 0 |
| RLS_050 | South Stirling | WA | 2020 | — | 6.81 ± 0.925 | 0 ± 0 | 0 ± 0 |
| RLS_051 | South Stirling | WA | 2020 | — | 6.3 ± 0.224 | 2.29 ± 2.975 | 4.05 ± 0.414 |
| RLS_052 | South Stirling | WA | 2020 | — | 1.68 ± 0.082 | 0 ± 0 | 0 ± 0 |
| RLS_053 | South Stirling | WA | 2020 | — | 14.44 ± 1.071 | 0.04 ± 0.041 | 0 ± 0 |
| RLS_054 | South Stirling | WA | 2020 | — | 3.27 ± 0.551 | 0 ± 0 | 0 ± 0 |
| RLS_055 | South Stirling | WA | 2020 | — | 0.55 ± 0.175 | 0 ± 0 | 0 ± 0 |
| RLS_056 | South Stirling | WA | 2020 | — | 35.67 ± 1.168 | 0 ± 0 | 0 ± 0 |
| RLS_057 | South Stirling | WA | 2020 | — | 9.47 ± 3.262 | 0 ± 0 | 0 ± 0 |
| RLS_058 | South Stirling | WA | 2020 | — | 0.55 ± 0.059 | 0.28 ± 0.064 | 0.17 ± 0.101 |
| RLS_059 | South Stirling | WA | 2020 | — | 5.01 ± 0.716 | 0 ± 0 | 0 ± 0 |
| RLS_060 | South Stirling | WA | 2020 | — | 1.63 ± 0.13 | 0 ± 0 | 0 ± 0 |
| RLS_061 | South Stirling | WA | 2020 | — | 1.14 ± 0.319 | 0 ± 0 | 0 ± 0 |
| RLS_062 | South Stirling | WA | 2020 | — | 1.32 ± 0.352 | 0 ± 0 | 0 ± 0 |
| RLS_063 | South Stirling | WA | 2020 | — | 3.54 ± 0.229 | 0 ± 0 | 0 ± 0 |
| RLS_064 | South Stirling | WA | 2020 | — | 15.28 ± 1.664 | 0 ± 0 | 0 ± 0 |
| RLS_065 | South Stirling | WA | 2020 | — | 15.07 ± 0.577 | 0 ± 0 | 0.28 ± 0.066 |
| RLS_066 | South Stirling | WA | 2020 | — | 6.05 ± 0.331 | 0 ± 0 | 0 ± 0 |
| RLS_067 | South Stirling | WA | 2020 | — | 26.3 ± 1.361 | 0 ± 0 | 0 ± 0 |
| RLS_068 | South Stirling | WA | 2020 | — | 1.3 ± 0.288 | 0 ± 0 | 0 ± 0 |
| RLS_069 | South Stirling | WA | 2020 | — | 0.93 ± 0.053 | 0 ± 0 | 0 ± 0 |
| RLS_070 | South Stirling | WA | 2020 | — | 4.5 ± 0.638 | 0 ± 0 | 0 ± 0 |
| RLS_071 | South Stirling | WA | 2020 | — | 2.04 ± 0.215 | 0 ± 0 | 0 ± 0 |
| RLS_072 | South Stirling | WA | 2020 | — | 1.12 ± 0.172 | 0 ± 0 | 0 ± 0 |
| RLS_073 | South Stirling | WA | 2020 | — | 15.34 ± 0.535 | 0 ± 0 | 0 ± 0 |
| RLS_074 | South Stirling | WA | 2020 | — | 11.95 ± 2.058 | 0.05 ± 0.053 | 0 ± 0 |
| RLS_075 | South Stirling | WA | 2020 | — | 0.58 ± 0.121 | 0 ± 0 | 0 ± 0 |
| RLS_076 | South Stirling | WA | 2020 | — | 4.39 ± 0.603 | 0 ± 0 | 0 ± 0 |
| RLS_077 | South Stirling | WA | 2020 | — | 0.76 ± 0.033 | 0 ± 0 | 0 ± 0 |
| RLS_078 | South Stirling | WA | 2020 | — | 1.3 ± 0.06 | 0.03 ± 0.033 | 0 ± 0 |
| RLS_079 | South Stirling | WA | 2020 | — | 17.94 ± 0.646 | 0 ± 0 | 0 ± 0 |
| RLS_080 | South Stirling | WA | 2020 | — | 7.7 ± 0.658 | 0.04 ± 0.037 | 0 ± 0 |
| RLS_081 | South Stirling | WA | 2020 | — | 1.79 ± 0.09 | 0 ± 0 | 0 ± 0 |
| RLS_082 | South Stirling | WA | 2020 | — | 263.61 ± 98.757 | 0.04 ± 0.035 | 0 ± 0 |
| RLS_083 | South Stirling | WA | 2020 | — | 6.28 ± 0.266 | 0 ± 0 | 0 ± 0 |
| RLS_084 | South Stirling | WA | 2020 | — | 1.13 ± 0.002 | 0.21 ± 0.03 | 0.21 ± 0.089 |
| RLS_085 | South Stirling | WA | 2020 | — | 5.18 ± 0.357 | 0 ± 0 | 0 ± 0 |
| RLS_086 | South Stirling | WA | 2020 | — | 11.67 ± 0.167 | 0 ± 0 | 0 ± 0 |
| RLS_087 | South Stirling | WA | 2020 | — | 24.31 ± 23.319 | 0.32 ± 0.32 | 0 ± 0 |
| RLS_088 | South Stirling | WA | 2020 | — | 54.97 ± 0.969 | 0 ± 0 | 0 ± 0 |
| RLS_089 | South Stirling | WA | 2020 | — | 3.6 ± 0.194 | 0.23 ± 0.159 | 0.23 ± 0.028 |
| RLS_090 | South Stirling | WA | 2020 | — | 5.52 ± 0.051 | 0.04 ± 0.035 | 0 ± 0 |
| RLS_091 | South Stirling | WA | 2020 | — | 3.63 ± 0.338 | 0 ± 0 | 0 ± 0 |
| RLS_092 | South Stirling | WA | 2020 | — | 13.27 ± 0.309 | 0 ± 0 | 0 ± 0 |
| RLS_093 | South Stirling | WA | 2020 | — | 4.12 ± 0.479 | 0 ± 0 | 0 ± 0 |
| RLS_094 | South Stirling | WA | 2020 | — | 3.14 ± 3.136 | 0 ± 0 | 0 ± 0 |
| RLS_095 | South Stirling | WA | 2020 | — | 0.3 ± 0.065 | 0 ± 0 | 0 ± 0 |
| RLS_096 | South Stirling | WA | 2020 | — | 5.26 ± 0.467 | 0 ± 0 | 0 ± 0 |
| RLS_097 | South Stirling | WA | 2020 | — | 1.7 ± 0.558 | 0 ± 0 | 0 ± 0 |
| RLS_098 | South Stirling | WA | 2020 | — | 2.39 ± 0.6 | 0 ± 0 | 0 ± 0 |
| RLS_099 | South Stirling | WA | 2020 | — | 0.37 ± 0.207 | 0 ± 0 | 0 ± 0 |
| RLS_100 | South Stirling | WA | 2020 | — | 2.71 ± 1.009 | 0.31 ± 0.31 | 0 ± 0 |
| RLS_101 | South Stirling | WA | 2020 | — | 0.53 ± 0.529 | 0.11 ± 0.106 | 0 ± 0 |
| RLS_102 | South Stirling | WA | 2020 | — | 4.39 ± 0.435 | 0.03 ± 0.033 | 0 ± 0 |
| RLS_103 | South Stirling | WA | 2020 | — | 8.87 ± 0.369 | 0 ± 0 | 0.03 ± 0.032 |
| RLS_104 | South Stirling | WA | 2020 | — | 1.85 ± 0.156 | 0.21 ± 0.073 | 0 ± 0 |
| RLS_105 | South Stirling | WA | 2020 | — | 6.24 ± 1.034 | 0.03 ± 0.034 | 0 ± 0 |
| RLS_106 | South Stirling | WA | 2020 | — | 18.18 ± 0.515 | 0 ± 0 | 0 ± 0 |
| RLS_107 | South Stirling | WA | 2020 | — | 8.22 ± 1.188 | 0 ± 0 | 0 ± 0 |
| RLS_108 | South Stirling | WA | 2020 | — | 5.95 ± 0.935 | 4.6 ± 4.596 | 0 ± 0 |
| RLS_109 | South Stirling | WA | 2020 | — | 2.59 ± 0.373 | 0 ± 0 | 0 ± 0 |
| RLS_110 | South Stirling | WA | 2020 | — | 9.44 ± 0.486 | 0 ± 0 | 0 ± 0 |

^a^ A dash (—) indicates no dilution was performed. A ‘10’ indicates a ten-fold dilution was performed.
