## Supplementary material for "Detection of *Ramularia collo-cygni* from barley (*Hordeum vulgare*) in Australia using triplex quantitative and digital PCR": S6

**Supplementary Table S6.** Mean quantification cycle (Cq) and DNA (pg) values with standard errors (*n* = 2) for *Hordeum vulgare* and *Ramularia collo-cygni* DNA in seed samples from Western Australia (WA) using *R. collo-cygni*-specific triplex and Ram6 quantitative PCR assays.

|  |  |  | *Hv*_116_*tef1-α* |  | *Rcc*_139_*rpb2* | | *Rcc*_88_*tef1-α* | |  | Ram6 (ITS) | |
| --- | --- | --- | --- | --- | --- | --- | --- | --- | --- | --- | --- |
| Sample | State | Year | Cq |  | Cq | DNA (pg) | Cq | DNA (pg) |  | Cq | DNA (pg) |
| RSS19_22_01 | WA | 2019 | 18.7 ± 0.05 |  | 0 ± 0 | 0 ± 0 | 0 ± 0 | 0 ± 0 |  | 29.8 ± 0.23 | 5 ± 0.51 |
| RSS19_22_02 | WA | 2019 | 18.4 ± 0.47 |  | 0 ± 0 | 0 ± 0 | 0 ± 0 | 0 ± 0 |  | 29.1 ± 0 | 6 ± 0.01 |
| RSS19_22_03 | WA | 2019 | 18.7 ± 0.15 |  | 0 ± 0 | 0 ± 0 | 0 ± 0 | 0 ± 0 |  | 28.9 ± 0.01 | 7 ± 0.05 |
| RSS19_22_04 | WA | 2019 | 20.3 ± 0.11 |  | 0 ± 0 | 0 ± 0 | 0 ± 0 | 0 ± 0 |  | 29 ± 0.08 | 7 ± 0.25 |
| RSS19_22_05 | WA | 2019 | 18.2 ± 0.01 |  | 0 ± 0 | 0 ± 0 | 0 ± 0 | 0 ± 0 |  | 28.1 ± 0.02 | 10 ± 0.1 |
| RSS19_22_06 | WA | 2019 | 20 ± 0.04 |  | 0 ± 0 | 0 ± 0 | 0 ± 0 | 0 ± 0 |  | 0 ± 0 | 0 ± 0 |
| RSS19_22_07 | WA | 2019 | 18.8 ± 0.17 |  | 0 ± 0 | 0 ± 0 | 0 ± 0 | 0 ± 0 |  | 28.6 ± 0.09 | 8 ± 0.33 |
| RSS19_22_08 | WA | 2019 | 21.4 ± 0.02 |  | 0 ± 0 | 0 ± 0 | 0 ± 0 | 0 ± 0 |  | 0 ± 0 | 0 ± 0 |
| RSS19_22_09 | WA | 2019 | 18 ± 0.18 |  | 0 ± 0 | 0 ± 0 | 0 ± 0 | 0 ± 0 |  | 29.9 ± 0.02 | 4 ± 0.05 |
| RSS19_22_10 | WA | 2019 | 19.2 ± 0.04 |  | 0 ± 0 | 0 ± 0 | 0 ± 0 | 0 ± 0 |  | 34.4 ± 0.41 | 1 ± 0.1 |
| RSS19_22_11 | WA | 2019 | 19.4 ± 0.04 |  | 0 ± 0 | 0 ± 0 | 0 ± 0 | 0 ± 0 |  | 34.4 ± 0.16 | 1 ± 0.04 |
| RSS19_22_12 | WA | 2019 | 19.1 ± 0 |  | 0 ± 0 | 0 ± 0 | 0 ± 0 | 0 ± 0 |  | 34.7 ± 0.31 | 0 ± 0.07 |
| RSS19_22_13 | WA | 2019 | 19.8 ± 0.16 |  | 0 ± 0 | 0 ± 0 | 0 ± 0 | 0 ± 0 |  | 33.9 ± 0.1 | 1 ± 0.03 |
| RSS19_22_14 | WA | 2019 | 20.2 ± 0.22 |  | 0 ± 0 | 0 ± 0 | 0 ± 0 | 0 ± 0 |  | 34.7 ± 0.14 | 0 ± 0.03 |
| RSS19_22_15 | WA | 2019 | 20.8 ± 0.39 |  | 0 ± 0 | 0 ± 0 | 0 ± 0 | 0 ± 0 |  | 34.7 ± 0.14 | 0 ± 0.03 |
| RSS19_22_16 | WA | 2019 | 21.6 ± 0.12 |  | 0 ± 0 | 0 ± 0 | 0 ± 0 | 0 ± 0 |  | 34.6 ± 0.07 | 0 ± 0 |
| RSS19_22_17 | WA | 2019 | 19.4 ± 0.08 |  | 0 ± 0 | 0 ± 0 | 0 ± 0 | 0 ± 0 |  | 34.6 ± 0.11 | 0 ± 0.03 |
| RSS19_22_18 | WA | 2019 | 21 ± 0.03 |  | 0 ± 0 | 0 ± 0 | 0 ± 0 | 0 ± 0 |  | 34.8 ± 0.28 | 0 ± 0.06 |
| RSS19_22_19 | WA | 2019 | 20.2 ± 0.04 |  | 0 ± 0 | 0 ± 0 | 0 ± 0 | 0 ± 0 |  | 34.5 ± 0 | 0 ± 0 |
| RSS19_22_20 | WA | 2019 | 19.9 ± 0.1 |  | 0 ± 0 | 0 ± 0 | 0 ± 0 | 0 ± 0 |  | 34 ± 0.08 | 1 ± 0.02 |
| RSS19_22_21 | WA | 2019 | 19.3 ± 0.15 |  | 0 ± 0 | 0 ± 0 | 0 ± 0 | 0 ± 0 |  | 34.5 ± 0.12 | 0 ± 0.03 |
| RSS19_22_22 | WA | 2019 | 19.5 ± 0.06 |  | 0 ± 0 | 0 ± 0 | 0 ± 0 | 0 ± 0 |  | 34.6 ± 0.4 | 0 ± 0.09 |
| RSS20_73_01 | WA | 2020 | 20.9 ± 0.11 |  | 0 ± 0 | 0 ± 0 | 0 ± 0 | 0 ± 0 |  | 0 ± 0 | 0 ± 0 |
| RSS20_73_02 | WA | 2020 | 19.6 ± 0.07 |  | 0 ± 0 | 0 ± 0 | 0 ± 0 | 0 ± 0 |  | 0 ± 0 | 0 ± 0 |
| RSS20_73_03 | WA | 2020 | 21 ± 0.11 |  | 0 ± 0 | 0 ± 0 | 0 ± 0 | 0 ± 0 |  | 0 ± 0 | 0 ± 0 |
| RSS20_73_04 | WA | 2020 | 22.5 ± 0.04 |  | 0 ± 0 | 0 ± 0 | 0 ± 0 | 0 ± 0 |  | 0 ± 0 | 0 ± 0 |
| RSS20_73_05 | WA | 2020 | 21 ± 0.11 |  | 0 ± 0 | 0 ± 0 | 0 ± 0 | 0 ± 0 |  | 0 ± 0 | 0 ± 0 |
| RSS20_73_06 | WA | 2020 | 21.5 ± 0.01 |  | 0 ± 0 | 0 ± 0 | 0 ± 0 | 0 ± 0 |  | 0 ± 0 | 0 ± 0 |
| RSS20_73_07 | WA | 2020 | 25.5 ± 0.46 |  | 0 ± 0 | 0 ± 0 | 0 ± 0 | 0 ± 0 |  | 0 ± 0 | 0 ± 0 |
| RSS20_73_08 | WA | 2020 | 26.4 ± 0.09 |  | 0 ± 0 | 0 ± 0 | 0 ± 0 | 0 ± 0 |  | 0 ± 0 | 0 ± 0 |
| RSS20_73_09 | WA | 2020 | 23.8 ± 0.48 |  | 0 ± 0 | 0 ± 0 | 0 ± 0 | 0 ± 0 |  | 0 ± 0 | 0 ± 0 |
| RSS20_73_10 | WA | 2020 | 25.1 ± 0.09 |  | 0 ± 0 | 0 ± 0 | 0 ± 0 | 0 ± 0 |  | 0 ± 0 | 0 ± 0 |
| RSS20_73_11 | WA | 2020 | 27.4 ± 0.04 |  | 0 ± 0 | 0 ± 0 | 0 ± 0 | 0 ± 0 |  | 0 ± 0 | 0 ± 0 |
| RSS20_73_12 | WA | 2020 | 23.7 ± 0.1 |  | 0 ± 0 | 0 ± 0 | 0 ± 0 | 0 ± 0 |  | 0 ± 0 | 0 ± 0 |
| RSS20_73_13 | WA | 2020 | 23.7 ± 0.42 |  | 0 ± 0 | 0 ± 0 | 0 ± 0 | 0 ± 0 |  | 0 ± 0 | 0 ± 0 |
| RSS20_73_14 | WA | 2020 | 24.4 ± 0.07 |  | 0 ± 0 | 0 ± 0 | 0 ± 0 | 0 ± 0 |  | 0 ± 0 | 0 ± 0 |
| RSS20_73_15 | WA | 2020 | 23.8 ± 0.39 |  | 0 ± 0 | 0 ± 0 | 0 ± 0 | 0 ± 0 |  | 0 ± 0 | 0 ± 0 |
| RSS20_73_16 | WA | 2020 | 22.1 ± 0.07 |  | 0 ± 0 | 0 ± 0 | 0 ± 0 | 0 ± 0 |  | 0 ± 0 | 0 ± 0 |
| RSS20_73_17 | WA | 2020 | 23.7 ± 0.15 |  | 0 ± 0 | 0 ± 0 | 0 ± 0 | 0 ± 0 |  | 0 ± 0 | 0 ± 0 |
| RSS20_73_18 | WA | 2020 | 25.5 ± 0.08 |  | 0 ± 0 | 0 ± 0 | 0 ± 0 | 0 ± 0 |  | 0 ± 0 | 0 ± 0 |
| RSS20_73_19 | WA | 2020 | 24.3 ± 0.07 |  | 0 ± 0 | 0 ± 0 | 0 ± 0 | 0 ± 0 |  | 0 ± 0 | 0 ± 0 |
| RSS20_73_20 | WA | 2020 | 21.2 ± 1.05 |  | 0 ± 0 | 0 ± 0 | 0 ± 0 | 0 ± 0 |  | 0 ± 0 | 0 ± 0 |
| RSS20_73_21 | WA | 2020 | 23 ± 0.26 |  | 0 ± 0 | 0 ± 0 | 0 ± 0 | 0 ± 0 |  | 0 ± 0 | 0 ± 0 |
| RSS20_73_22 | WA | 2020 | 21.8 ± 0.03 |  | 0 ± 0 | 0 ± 0 | 0 ± 0 | 0 ± 0 |  | 0 ± 0 | 0 ± 0 |
| RSS20_73_23 | WA | 2020 | 25.5 ± 1.18 |  | 0 ± 0 | 0 ± 0 | 0 ± 0 | 0 ± 0 |  | 0 ± 0 | 0 ± 0 |
| RSS20_73_24 | WA | 2020 | 23.5 ± 0.72 |  | 0 ± 0 | 0 ± 0 | 0 ± 0 | 0 ± 0 |  | 0 ± 0 | 0 ± 0 |
| RSS20_73_25 | WA | 2020 | 23.7 ± 0.36 |  | 0 ± 0 | 0 ± 0 | 0 ± 0 | 0 ± 0 |  | 0 ± 0 | 0 ± 0 |
| RSS20_73_26 | WA | 2020 | 24.7 ± 0.22 |  | 0 ± 0 | 0 ± 0 | 0 ± 0 | 0 ± 0 |  | 0 ± 0 | 0 ± 0 |
| RSS20_73_27 | WA | 2020 | 24.9 ± 0.01 |  | 0 ± 0 | 0 ± 0 | 0 ± 0 | 0 ± 0 |  | 0 ± 0 | 0 ± 0 |
| RSS20_73_28 | WA | 2020 | 25.1 ± 0.11 |  | 0 ± 0 | 0 ± 0 | 0 ± 0 | 0 ± 0 |  | 0 ± 0 | 0 ± 0 |
| RSS20_73_29 | WA | 2020 | 26.6 ± 0.77 |  | 0 ± 0 | 0 ± 0 | 0 ± 0 | 0 ± 0 |  | 0 ± 0 | 0 ± 0 |
| RSS20_73_30 | WA | 2020 | 24.2 ± 0.21 |  | 0 ± 0 | 0 ± 0 | 0 ± 0 | 0 ± 0 |  | 0 ± 0 | 0 ± 0 |
| RSS20_73_31 | WA | 2020 | 22.3 ± 0.45 |  | 0 ± 0 | 0 ± 0 | 0 ± 0 | 0 ± 0 |  | 0 ± 0 | 0 ± 0 |
| RSS20_73_32 | WA | 2020 | 22.1 ± 0.21 |  | 0 ± 0 | 0 ± 0 | 0 ± 0 | 0 ± 0 |  | 0 ± 0 | 0 ± 0 |
| RSS20_73_33 | WA | 2020 | 20.9 ± 0.14 |  | 0 ± 0 | 0 ± 0 | 0 ± 0 | 0 ± 0 |  | 0 ± 0 | 0 ± 0 |
| RSS20_73_34 | WA | 2020 | 19.6 ± 0.01 |  | 0 ± 0 | 0 ± 0 | 0 ± 0 | 0 ± 0 |  | 0 ± 0 | 0 ± 0 |
| RSS20_73_35 | WA | 2020 | 21.2 ± 0.03 |  | 0 ± 0 | 0 ± 0 | 0 ± 0 | 0 ± 0 |  | 0 ± 0 | 0 ± 0 |
| RSS20_73_36 | WA | 2020 | 20.6 ± 0.07 |  | 0 ± 0 | 0 ± 0 | 0 ± 0 | 0 ± 0 |  | 0 ± 0 | 0 ± 0 |
| RSS20_73_37 | WA | 2020 | 23.1 ± 5.84 |  | 0 ± 0 | 0 ± 0 | 0 ± 0 | 0 ± 0 |  | 0 ± 0 | 0 ± 0 |
| RSS20_73_38 | WA | 2020 | 20 ± 0.1 |  | 0 ± 0 | 0 ± 0 | 0 ± 0 | 0 ± 0 |  | 0 ± 0 | 0 ± 0 |
| RSS20_73_39 | WA | 2020 | 20.1 ± 0.01 |  | 0 ± 0 | 0 ± 0 | 0 ± 0 | 0 ± 0 |  | 0 ± 0 | 0 ± 0 |
| RSS20_73_40 | WA | 2020 | 26.7 ± 0.68 |  | 0 ± 0 | 0 ± 0 | 0 ± 0 | 0 ± 0 |  | 0 ± 0 | 0 ± 0 |
| RSS20_73_41 | WA | 2020 | 21 ± 0.1 |  | 0 ± 0 | 0 ± 0 | 0 ± 0 | 0 ± 0 |  | 0 ± 0 | 0 ± 0 |
| RSS20_73_42 | WA | 2020 | 21 ± 0.02 |  | 0 ± 0 | 0 ± 0 | 0 ± 0 | 0 ± 0 |  | 0 ± 0 | 0 ± 0 |
| RSS20_73_43 | WA | 2020 | 23.1 ± 0.04 |  | 0 ± 0 | 0 ± 0 | 0 ± 0 | 0 ± 0 |  | 0 ± 0 | 0 ± 0 |
| RSS20_73_44 | WA | 2020 | 21.8 ± 0.06 |  | 0 ± 0 | 0 ± 0 | 0 ± 0 | 0 ± 0 |  | 0 ± 0 | 0 ± 0 |
| RSS20_73_45 | WA | 2020 | 22.5 ± 0.04 |  | 0 ± 0 | 0 ± 0 | 0 ± 0 | 0 ± 0 |  | 0 ± 0 | 0 ± 0 |
| RSS20_73_46 | WA | 2020 | 23.2 ± 0.06 |  | 0 ± 0 | 0 ± 0 | 0 ± 0 | 0 ± 0 |  | 0 ± 0 | 0 ± 0 |
| RSS20_73_47 | WA | 2020 | 21.3 ± 0.01 |  | 0 ± 0 | 0 ± 0 | 0 ± 0 | 0 ± 0 |  | 0 ± 0 | 0 ± 0 |
| RSS20_73_48 | WA | 2020 | 22.2 ± 0.31 |  | 0 ± 0 | 0 ± 0 | 0 ± 0 | 0 ± 0 |  | 0 ± 0 | 0 ± 0 |
| RSS20_73_49 | WA | 2020 | 25.3 ± 0.05 |  | 0 ± 0 | 0 ± 0 | 0 ± 0 | 0 ± 0 |  | 0 ± 0 | 0 ± 0 |
| RSS20_73_50 | WA | 2020 | 23.3 ± 0.3 |  | 0 ± 0 | 0 ± 0 | 0 ± 0 | 0 ± 0 |  | 0 ± 0 | 0 ± 0 |
| RSS20_73_51 | WA | 2020 | 22.4 ± 0.1 |  | 0 ± 0 | 0 ± 0 | 0 ± 0 | 0 ± 0 |  | 0 ± 0 | 0 ± 0 |
| RSS20_73_52 | WA | 2020 | 23.7 ± 0.09 |  | 0 ± 0 | 0 ± 0 | 0 ± 0 | 0 ± 0 |  | 0 ± 0 | 0 ± 0 |
| RSS20_73_53 | WA | 2020 | 22.9 ± 0.06 |  | 0 ± 0 | 0 ± 0 | 0 ± 0 | 0 ± 0 |  | 0 ± 0 | 0 ± 0 |
| RSS20_73_54 | WA | 2020 | 23.4 ± 0.2 |  | 0 ± 0 | 0 ± 0 | 0 ± 0 | 0 ± 0 |  | 0 ± 0 | 0 ± 0 |
| RSS20_73_55 | WA | 2020 | 22.4 ± 0.17 |  | 0 ± 0 | 0 ± 0 | 0 ± 0 | 0 ± 0 |  | 0 ± 0 | 0 ± 0 |
| RSS20_73_56 | WA | 2020 | 22.3 ± 0 |  | 0 ± 0 | 0 ± 0 | 0 ± 0 | 0 ± 0 |  | 0 ± 0 | 0 ± 0 |
| RSS20_73_57 | WA | 2020 | 22.9 ± 0.13 |  | 0 ± 0 | 0 ± 0 | 0 ± 0 | 0 ± 0 |  | 0 ± 0 | 0 ± 0 |
| RSS20_73_58 | WA | 2020 | 22.8 ± 0.12 |  | 0 ± 0 | 0 ± 0 | 0 ± 0 | 0 ± 0 |  | 0 ± 0 | 0 ± 0 |
| RSS20_73_59 | WA | 2020 | 23 ± 0.04 |  | 0 ± 0 | 0 ± 0 | 0 ± 0 | 0 ± 0 |  | 0 ± 0 | 0 ± 0 |
| RSS20_73_60 | WA | 2020 | 23.1 ± 0.04 |  | 0 ± 0 | 0 ± 0 | 0 ± 0 | 0 ± 0 |  | 0 ± 0 | 0 ± 0 |
| RSS20_73_61 | WA | 2020 | 23.4 ± 0.06 |  | 0 ± 0 | 0 ± 0 | 0 ± 0 | 0 ± 0 |  | 0 ± 0 | 0 ± 0 |
| RSS20_73_62 | WA | 2020 | 23 ± 0.07 |  | 0 ± 0 | 0 ± 0 | 0 ± 0 | 0 ± 0 |  | 0 ± 0 | 0 ± 0 |
| RSS20_73_63 | WA | 2020 | 22.6 ± 0.17 |  | 0 ± 0 | 0 ± 0 | 0 ± 0 | 0 ± 0 |  | 0 ± 0 | 0 ± 0 |
| RSS20_73_64 | WA | 2020 | 22.3 ± 0.67 |  | 0 ± 0 | 0 ± 0 | 0 ± 0 | 0 ± 0 |  | 0 ± 0 | 0 ± 0 |
| RSS20_73_65 | WA | 2020 | 22.3 ± 0.05 |  | 0 ± 0 | 0 ± 0 | 0 ± 0 | 0 ± 0 |  | 0 ± 0 | 0 ± 0 |
| RSS20_73_66 | WA | 2020 | 23.1 ± 0.04 |  | 0 ± 0 | 0 ± 0 | 0 ± 0 | 0 ± 0 |  | 0 ± 0 | 0 ± 0 |
| RSS20_73_67 | WA | 2020 | 23.6 ± 0.03 |  | 0 ± 0 | 0 ± 0 | 0 ± 0 | 0 ± 0 |  | 0 ± 0 | 0 ± 0 |
| RSS20_73_68 | WA | 2020 | 23.4 ± 0.32 |  | 0 ± 0 | 0 ± 0 | 0 ± 0 | 0 ± 0 |  | 0 ± 0 | 0 ± 0 |
| RSS20_73_69 | WA | 2020 | 22.3 ± 0.04 |  | 0 ± 0 | 0 ± 0 | 0 ± 0 | 0 ± 0 |  | 0 ± 0 | 0 ± 0 |
| RSS20_73_70 | WA | 2020 | 26.2 ± 1.84 |  | 0 ± 0 | 0 ± 0 | 0 ± 0 | 0 ± 0 |  | 0 ± 0 | 0 ± 0 |
| RSS20_73_71 | WA | 2020 | 33.3 ± 1.57 |  | 0 ± 0 | 0 ± 0 | 0 ± 0 | 0 ± 0 |  | 0 ± 0 | 0 ± 0 |
| RSS20_73_72 | WA | 2020 | 26.1 ± 0.55 |  | 0 ± 0 | 0 ± 0 | 0 ± 0 | 0 ± 0 |  | 0 ± 0 | 0 ± 0 |
| RSS20_73_73 | WA | 2020 | 21.9 ± 0.12 |  | 0 ± 0 | 0 ± 0 | 0 ± 0 | 0 ± 0 |  | 0 ± 0 | 0 ± 0 |
