## Supplementary material for "Detection of *Ramularia collo-cygni* from barley (*Hordeum vulgare*) in Australia using triplex quantitative and digital PCR": S7

**Supplementary Table S7.** Mean copy number values and standard errors (*n* = 2) for *Hordeum vulgare* and *Ramularia collo-cygni* DNA in seed samples from Western Australia (WA) using *R. collo-cygni*-specific triplex droplet digital PCR.

| Sample | State | Year | Dilution^a^ | *Hv*_116_*tef1-α* copies µL^-1^ | *Rcc*_139_*rpb2* copies µL^-1^ | *Rcc*_88_*tef1-α* copies µL^-1^ |
| --- | --- | --- | --- | --- | --- | --- |
| 22-RSS 001 | WA | 2019 | — | 1000000 ± 0 | 0 ± 0 | 0 ± 0 |
| 22-RSS 002 | WA | 2019 | — | 9827.54 ± 921.134 | 0 ± 0 | 0.04 ± 0.038 |
| 22-RSS 003 | WA | 2019 | — | 500000 ± 500000 | 0 ± 0 | 0.04 ± 0.041 |
| 22-RSS 004 | WA | 2019 | — | 4941.6 ± 222.38 | 0 ± 0 | 0.25 ± 0.104 |
| 22-RSS 005 | WA | 2019 | — | 505724.31 ± 494275.686 | 0 ± 0 | 0.03 ± 0.035 |
| 22-RSS 006 | WA | 2019 | — | 5123.74 ± 105.734 | 0.15 ± 0.005 | 0.04 ± 0.037 |
| 22-RSS 007 | WA | 2019 | — | 6319.18 ± 392.534 | 0 ± 0 | 0.08 ± 0.082 |
| 22-RSS 008 | WA | 2019 | — | 2318.03 ± 49.492 | 0 ± 0 | 0 ± 0 |
| 22-RSS 009 | WA | 2019 | — | 6582.79 ± 38.196 | 0 ± 0 | 0 ± 0 |
| 22-RSS 010 | WA | 2019 | — | 7318.81 ± 132.187 | 0 ± 0 | 0 ± 0 |
| 22-RSS 011 | WA | 2019 | — | 7773.92 ± 1149.117 | 0 ± 0 | 0 ± 0 |
| 22-RSS 012 | WA | 2019 | — | 229.6 ± 21.326 | 0 ± 0 | 0 ± 0 |
| 22-RSS 013 | WA | 2019 | — | 6118.26 ± 247.161 | 0 ± 0 | 0 ± 0 |
| 22-RSS 014 | WA | 2019 | — | 4521.21 ± 544.783 | 0 ± 0 | 0.04 ± 0.044 |
| 22-RSS 015 | WA | 2019 | — | 5017.93 ± 458.308 | 0 ± 0 | 0 ± 0 |
| 22-RSS 016 | WA | 2019 | — | 1633.32 ± 107.969 | 0 ± 0 | 0 ± 0 |
| 22-RSS 017 | WA | 2019 | — | 6651.51 ± 221.248 | 0.03 ± 0.034 | 0.03 ± 0.034 |
| 22-RSS 018 | WA | 2019 | — | 2249.74 ± 84.115 | 0 ± 0 | 0.04 ± 0.038 |
| 22-RSS 019 | WA | 2019 | — | 4107.22 ± 128.48 | 0 ± 0 | 0 ± 0 |
| 22-RSS 020 | WA | 2019 | — | 6387.71 ± 617.146 | 0 ± 0 | 0 ± 0 |
| 22-RSS 021 | WA | 2019 | — | 6985.46 ± 132.023 | 0 ± 0 | 0 ± 0 |
| 22-RSS 022 | WA | 2019 | — | 5053.44 ± 1818.959 | 0 ± 0 | 0 ± 0 |
| 22-RSS 001 | WA | 2019 | 10 | 1389.51 ± 127.225 | 0 ± 0 | 0 ± 0 |
| 22-RSS 002 | WA | 2019 | 10 | 856.56 ± 15.103 | 0 ± 0 | 0 ± 0 |
| 22-RSS 003 | WA | 2019 | 10 | 1370.25 ± 72.182 | 0 ± 0 | 0 ± 0 |
| 22-RSS 004 | WA | 2019 | 10 | 369.9 ± 25.923 | 0 ± 0 | 0 ± 0 |
| 22-RSS 005 | WA | 2019 | 10 | 1391.36 ± 50.102 | 0 ± 0 | 0 ± 0 |
| 22-RSS 006 | WA | 2019 | 10 | 1020.33 ± 16.807 | 0 ± 0 | 0.03 ± 0.032 |
| 22-RSS 007 | WA | 2019 | 10 | 1234.74 ± 64.136 | 0 ± 0 | 0.03 ± 0.032 |
| 22-RSS 008 | WA | 2019 | 10 | 203.45 ± 17.919 | 0 ± 0 | 0 ± 0 |
| 22-RSS 009 | WA | 2019 | 10 | 536.15 ± 67.659 | 0 ± 0 | 0 ± 0 |
| 22-RSS 010 | WA | 2019 | 10 | 762.35 ± 61.736 | 0 ± 0 | 0 ± 0 |
| 22-RSS 011 | WA | 2019 | 10 | 736.02 ± 2.379 | 0 ± 0 | 0 ± 0 |
| 22-RSS 012 | WA | 2019 | 10 | 34.23 ± 20.916 | 0 ± 0 | 0 ± 0 |
| 22-RSS 013 | WA | 2019 | 10 | 649.76 ± 1.001 | 0 ± 0 | 0 ± 0 |
| 22-RSS 014 | WA | 2019 | 10 | 444.56 ± 10.256 | 0 ± 0 | 0 ± 0 |
| 22-RSS 015 | WA | 2019 | 10 | 364.3 ± 24.384 | 0 ± 0 | 0 ± 0 |
| 22-RSS 016 | WA | 2019 | 10 | 134.54 ± 4.315 | 0 ± 0 | 0 ± 0 |
| 22-RSS 017 | WA | 2019 | 10 | 669.27 ± 10.121 | 0 ± 0 | 0 ± 0 |
| 22-RSS 018 | WA | 2019 | 10 | 183.02 ± 19.134 | 0 ± 0 | 0 ± 0 |
| 22-RSS 019 | WA | 2019 | 10 | 286.9 ± 6.28 | 0 ± 0 | 0.03 ± 0.031 |
| 22-RSS 020 | WA | 2019 | 10 | 337.75 ± 12.828 | 0 ± 0 | 0 ± 0 |
| 22-RSS 021 | WA | 2019 | 10 | 684.43 ± 35.517 | 0 ± 0 | 0.03 ± 0.03 |
| 22-RSS 022 | WA | 2019 | 10 | 661.5 ± 4.472 | 0 ± 0 | 0 ± 0 |
| 73-RSS 001 | WA | 2020 | — | 6627.1 ± 40.988 | 0 ± 0 | 0 ± 0 |
| 73-RSS 002 | WA | 2020 | — | 6981.06 ± 94.616 | 0 ± 0 | 0 ± 0 |
| 73-RSS 003 | WA | 2020 | — | 3819.99 ± 13.848 | 0 ± 0 | 0 ± 0 |
| 73-RSS 004 | WA | 2020 | — | 1457.02 ± 90.906 | 0 ± 0 | 0 ± 0 |
| 73-RSS 005 | WA | 2020 | — | 2420.06 ± 61.431 | 0 ± 0 | 0 ± 0 |
| 73-RSS 006 | WA | 2020 | — | 4034.26 ± 242.163 | 0 ± 0 | 0 ± 0 |
| 73-RSS 007 | WA | 2020 | — | 774.68 ± 2.967 | 0 ± 0 | 0.04 ± 0.039 |
| 73-RSS 008 | WA | 2020 | — | 403.86 ± 20.733 | 0 ± 0 | 0 ± 0 |
| 73-RSS 009 | WA | 2020 | — | 2175.75 ± 23.666 | 0.04 ± 0.035 | 0 ± 0 |
| 73-RSS 010 | WA | 2020 | — | 1808.6 ± 22.747 | 0 ± 0 | 0 ± 0 |
| 73-RSS 011 | WA | 2020 | — | 190.4 ± 9.056 | 0.03 ± 0.032 | 0 ± 0 |
| 73-RSS 012 | WA | 2020 | — | 1206 ± 19.088 | 0 ± 0 | 0 ± 0 |
| 73-RSS 013 | WA | 2020 | — | 591.78 ± 555.475 | 0 ± 0 | 0 ± 0 |
| 73-RSS 014 | WA | 2020 | — | 3.48 ± 2.548 | 0 ± 0 | 0 ± 0 |
| 73-RSS 015 | WA | 2020 | — | 2446.56 ± 90.907 | 0 ± 0 | 0 ± 0 |
| 73-RSS 016 | WA | 2020 | — | 4258.74 ± 106.662 | 0 ± 0 | 0 ± 0 |
| 73-RSS 017 | WA | 2020 | — | 2612.19 ± 235.683 | 0 ± 0 | 0 ± 0 |
| 73-RSS 018 | WA | 2020 | — | 2667.33 ± 20.369 | 0 ± 0 | 0 ± 0 |
| 73-RSS 019 | WA | 2020 | — | 2067.68 ± 192.067 | 0.07 ± 0.066 | 0 ± 0 |
| 73-RSS 020 | WA | 2020 | — | 1636.5 ± 506.228 | 0.08 ± 0.078 | 0 ± 0 |
| 73-RSS 021 | WA | 2020 | — | 3791.24 ± 259.044 | 0.39 ± 0.082 | 0 ± 0 |
| 73-RSS 022 | WA | 2020 | — | 4464.74 ± 140.847 | 0.08 ± 0.075 | 0.04 ± 0.038 |
| 73-RSS 023 | WA | 2020 | — | 1299.87 ± 12.643 | 0 ± 0 | 0 ± 0 |
| 73-RSS 024 | WA | 2020 | — | 2476.38 ± 87.873 | 0 ± 0 | 0 ± 0 |
| 73-RSS 025 | WA | 2020 | — | 4033.23 ± 141.895 | 0 ± 0 | 0 ± 0 |
| 73-RSS 026 | WA | 2020 | — | 1651.08 ± 104.364 | 0 ± 0 | 0 ± 0 |
| 73-RSS 027 | WA | 2020 | — | 3224.1 ± 21.242 | 0.05 ± 0.047 | 0.04 ± 0.042 |
| 73-RSS 028 | WA | 2020 | — | 4333.68 ± 0.62 | 0 ± 0 | 0 ± 0 |
| 73-RSS 029 | WA | 2020 | — | 5608.07 ± 285.331 | 0 ± 0 | 0 ± 0 |
| 73-RSS 030 | WA | 2020 | — | 6072.81 ± 106.502 | 0 ± 0 | 0.08 ± 0.082 |
| 73-RSS 031 | WA | 2020 | — | 5861.41 ± 104.649 | 0 ± 0 | 0 ± 0 |
| 73-RSS 032 | WA | 2020 | — | 4308.44 ± 64.842 | 0 ± 0 | 0.04 ± 0.038 |
| 73-RSS 033 | WA | 2020 | — | 2700.15 ± 15.368 | 0 ± 0 | 0 ± 0 |
| 73-RSS 034 | WA | 2020 | — | 4840.46 ± 39.559 | 0 ± 0 | 0 ± 0 |
| 73-RSS 035 | WA | 2020 | — | 1618.53 ± 36.343 | 0.1 ± 0.034 | 0.03 ± 0.034 |
| 73-RSS 036 | WA | 2020 | — | 1957.56 ± 12.154 | 0 ± 0 | 0 ± 0 |
| 73-RSS 037 | WA | 2020 | — | 3584.45 ± 38.781 | 0 ± 0 | 0 ± 0 |
| 73-RSS 038 | WA | 2020 | — | 2462.87 ± 24.986 | 0 ± 0 | 0 ± 0 |
| 73-RSS 039 | WA | 2020 | — | 2927.97 ± 21.238 | 0 ± 0 | 0 ± 0 |
| 73-RSS 040 | WA | 2020 | — | 5212.95 ± 11.345 | 0.45 ± 0.121 | 0.04 ± 0.035 |
| 73-RSS 041 | WA | 2020 | — | ND^b^ | ND | ND |
| 73-RSS 042 | WA | 2020 | — | ND | ND | ND |
| 73-RSS 043 | WA | 2020 | — | 2994.82 ± 80.162 | 0 ± 0 | 0 ± 0 |
| 73-RSS 044 | WA | 2020 | — | 4138.36 ± 114.731 | 0 ± 0 | 0 ± 0 |
| 73-RSS 045 | WA | 2020 | — | 4368.13 ± 27.68 | 0 ± 0 | 0 ± 0 |
| 73-RSS 046 | WA | 2020 | — | 4724.2 ± 58.265 | 0.04 ± 0.035 | 0 ± 0 |
| 73-RSS 047 | WA | 2020 | — | 6941.96 ± 129.453 | 0 ± 0 | 0 ± 0 |
| 73-RSS 048 | WA | 2020 | — | 6415.81 ± 126.717 | 0 ± 0 | 0 ± 0 |
| 73-RSS 049 | WA | 2020 | — | 3414.14 ± 242.055 | 0.31 ± 0.032 | 0.15 ± 0.016 |
| 73-RSS 050 | WA | 2020 | — | 3906 ± 1045.977 | 0.04 ± 0.04 | 0 ± 0 |
| 73-RSS 051 | WA | 2020 | — | 5464.21 ± 298.565 | 0.14 ± 0.05 | 0 ± 0 |
| 73-RSS 052 | WA | 2020 | — | 5356.18 ± 45.568 | 0 ± 0 | 0 ± 0 |
| 73-RSS 053 | WA | 2020 | — | 3158.41 ± 99.542 | 0 ± 0 | 0 ± 0 |
| 73-RSS 054 | WA | 2020 | — | 3906.26 ± 14.56 | 0.05 ± 0.048 | 0 ± 0 |
| 73-RSS 055 | WA | 2020 | — | 3908.91 ± 32.426 | 0 ± 0 | 0.04 ± 0.044 |
| 73-RSS 056 | WA | 2020 | — | 3563.3 ± 9.822 | 0 ± 0 | 0 ± 0 |
| 73-RSS 057 | WA | 2020 | — | 3132.36 ± 79.391 | 0 ± 0 | 0 ± 0 |
| 73-RSS 058 | WA | 2020 | — | ND | ND | ND |
| 73-RSS 059 | WA | 2020 | — | 2763.53 ± 57.311 | 3.23 ± 1.88 | 0.25 ± 0.106 |
| 73-RSS 060 | WA | 2020 | — | #DIV/0! | #DIV/0! | #DIV/0! |
| 73-RSS 061 | WA | 2020 | — | 3272.08 ± 164.305 | 0.09 ± 0.031 | 0 ± 0 |
| 73-RSS 062 | WA | 2020 | — | 1662.5 ± 569.997 | 1.26 ± 1.117 | 0.15 ± 0.149 |
| 73-RSS 063 | WA | 2020 | — | ND | ND | ND |
| 73-RSS 064 | WA | 2020 | — | 317.02 ± 317.019 | 0.73 ± 0.733 | 0 ± 0 |
| 73-RSS 065 | WA | 2020 | — | 2630.09 ± 249.924 | 0 ± 0 | 0 ± 0 |
| 73-RSS 066 | WA | 2020 | — | 148.7 ± 50.391 | 0.03 ± 0.033 | 0 ± 0 |
| 73-RSS 067 | WA | 2020 | — | 0 ± 0 | 0 ± 0 | 0 ± 0 |
| 73-RSS 068 | WA | 2020 | — | 0.32 ± 0.318 | 0 ± 0 | 0 ± 0 |
| 73-RSS 069 | WA | 2020 | — | 3349.02 ± 93.998 | 0 ± 0 | 0 ± 0 |
| 73-RSS 070 | WA | 2020 | — | 1822.9 ± 3.683 | 0.08 ± 0.08 | 0.08 ± 0.08 |
| 73-RSS 071 | WA | 2020 | — | 2600.39 ± 72.439 | 0 ± 0 | 0.04 ± 0.04 |
| 73-RSS 072 | WA | 2020 | — | 1801.22 ± 1801.217 | 0 ± 0 | 0 ± 0 |
| 73-RSS 073 | WA | 2020 | — | 2540.67 ± 6.199 | 0.07 ± 0 | 0.03 ± 0.034 |
| 73-RSS 001 | WA | 2020 | 10 | 951.58 ± 52.445 | 0 ± 0 | 0 ± 0 |
| 73-RSS 002 | WA | 2020 | 10 | 1459.59 ± 29.369 | 0 ± 0 | 0 ± 0 |
| 73-RSS 003 | WA | 2020 | 10 | 322.12 ± 5.913 | 0 ± 0 | 0 ± 0 |
| 73-RSS 004 | WA | 2020 | 10 | 118.57 ± 1.025 | 0 ± 0 | 0 ± 0 |
| 73-RSS 005 | WA | 2020 | 10 | 507.01 ± 35.017 | 0 ± 0 | 0 ± 0 |
| 73-RSS 006 | WA | 2020 | 10 | 578.47 ± 16.645 | 0.03 ± 0.034 | 0 ± 0 |
| 73-RSS 007 | WA | 2020 | 10 | 71.85 ± 2.329 | 0 ± 0 | 0 ± 0 |
| 73-RSS 008 | WA | 2020 | 10 | 46.59 ± 1.369 | 0.03 ± 0.03 | 0 ± 0 |
| 73-RSS 009 | WA | 2020 | 10 | 92.84 ± 12.278 | 0 ± 0 | 0 ± 0 |
| 73-RSS 010 | WA | 2020 | 10 | 121.97 ± 10.512 | 0 ± 0 | 0 ± 0 |
| 73-RSS 011 | WA | 2020 | 10 | 3.39 ± 0.92 | 0 ± 0 | 0 ± 0 |
| 73-RSS 012 | WA | 2020 | 10 | 186.57 ± 8.872 | 0 ± 0 | 0 ± 0 |
| 73-RSS 013 | WA | 2020 | 10 | 1630.13 ± 10.571 | 0 ± 0 | 0 ± 0 |
| 73-RSS 014 | WA | 2020 | 10 | 1056.77 ± 73.351 | 0 ± 0 | 0 ± 0 |
| 73-RSS 015 | WA | 2020 | 10 | 308.26 ± 9.937 | 0 ± 0 | 0 ± 0 |
| 73-RSS 016 | WA | 2020 | 10 | 491.05 ± 5.532 | 0 ± 0 | 0 ± 0 |
| 73-RSS 017 | WA | 2020 | 10 | 429.64 ± 27.983 | 0 ± 0 | 0 ± 0 |
| 73-RSS 018 | WA | 2020 | 10 | 416.63 ± 13.616 | 0 ± 0 | 0 ± 0 |
| 73-RSS 019 | WA | 2020 | 10 | 630.1 ± 65.647 | 0 ± 0 | 0 ± 0 |
| 73-RSS 020 | WA | 2020 | 10 | 557.33 ± 29.021 | 0.03 ± 0.033 | 0 ± 0 |
| 73-RSS 021 | WA | 2020 | 10 | 474.6 ± 8.906 | 0 ± 0 | 0 ± 0 |
| 73-RSS 022 | WA | 2020 | 10 | 528.18 ± 10.62 | 0 ± 0 | 0 ± 0 |
| 73-RSS 023 | WA | 2020 | 10 | 57.69 ± 5.811 | 0 ± 0 | 0 ± 0 |
| 73-RSS 024 | WA | 2020 | 10 | 187.8 ± 11.815 | 0 ± 0 | 0 ± 0 |
| 73-RSS 025 | WA | 2020 | 10 | 425.66 ± 13.54 | 0 ± 0 | 0 ± 0 |
| 73-RSS 026 | WA | 2020 | 10 | 97.49 ± 3.197 | 0 ± 0 | 0 ± 0 |
| 73-RSS 027 | WA | 2020 | 10 | 304.33 ± 9.974 | 0 ± 0 | 0 ± 0 |
| 73-RSS 028 | WA | 2020 | 10 | 388.35 ± 18.913 | 0 ± 0 | 0.03 ± 0.035 |
| 73-RSS 029 | WA | 2020 | 10 | 598.72 ± 45.946 | 0 ± 0 | 0.07 ± 0.071 |
| 73-RSS 030 | WA | 2020 | 10 | 505.99 ± 14.718 | 0 ± 0 | 0 ± 0 |
| 73-RSS 031 | WA | 2020 | 10 | 468.54 ± 12.95 | 0 ± 0 | 0 ± 0 |
| 73-RSS 032 | WA | 2020 | 10 | 404.76 ± 21.11 | 0 ± 0 | 0 ± 0 |
| 73-RSS 033 | WA | 2020 | 10 | 132.17 ± 7.658 | 0 ± 0 | 0 ± 0 |
| 73-RSS 034 | WA | 2020 | 10 | 351.2 ± 11.969 | 0 ± 0 | 0 ± 0 |
| 73-RSS 035 | WA | 2020 | 10 | 99.93 ± 8.923 | 0 ± 0 | 0 ± 0 |
| 73-RSS 036 | WA | 2020 | 10 | 101.63 ± 101.634 | 0 ± 0 | 0 ± 0 |
| 73-RSS 037 | WA | 2020 | 10 | 350.33 ± 8.23 | 0 ± 0 | 0 ± 0 |
| 73-RSS 038 | WA | 2020 | 10 | 256.81 ± 2.051 | 0 ± 0 | 0 ± 0 |
| 73-RSS 039 | WA | 2020 | 10 | 233.1 ± 17.754 | 0 ± 0 | 0 ± 0 |
| 73-RSS 040 | WA | 2020 | 10 | 479.9 ± 12.096 | 0 ± 0 | 0 ± 0 |
| 73-RSS 041 | WA | 2020 | 10 | 804.19 ± 16.57 | 0.17 ± 0.17 | 0.09 ± 0.085 |
| 73-RSS 042 | WA | 2020 | 10 | 690.73 ± 15.391 | 0 ± 0 | 0 ± 0 |
| 73-RSS 043 | WA | 2020 | 10 | 230.04 ± 1.023 | 0 ± 0 | 0 ± 0 |
| 73-RSS 044 | WA | 2020 | 10 | 357.59 ± 35.603 | 0 ± 0 | 0 ± 0 |
| 73-RSS 045 | WA | 2020 | 10 | 422.3 ± 8.022 | 0 ± 0 | 0 ± 0 |
| 73-RSS 046 | WA | 2020 | 10 | 379.36 ± 6.326 | 0 ± 0 | 0 ± 0 |
| 73-RSS 047 | WA | 2020 | 10 | 831.37 ± 2.345 | 0 ± 0 | 0 ± 0 |
| 73-RSS 048 | WA | 2020 | 10 | 566.47 ± 11.838 | 0 ± 0 | 0 ± 0 |
| 73-RSS 049 | WA | 2020 | 10 | 215.19 ± 24.709 | 0 ± 0 | 0 ± 0 |
| 73-RSS 050 | WA | 2020 | 10 | 473.02 ± 28.188 | 0 ± 0 | 0 ± 0 |
| 73-RSS 051 | WA | 2020 | 10 | 568.08 ± 24.138 | 0 ± 0 | 0 ± 0 |
| 73-RSS 052 | WA | 2020 | 10 | 485.54 ± 3.335 | 0 ± 0 | 0 ± 0 |
| 73-RSS 053 | WA | 2020 | 10 | 227.14 ± 6.359 | 0 ± 0 | 0 ± 0 |
| 73-RSS 054 | WA | 2020 | 10 | 375.43 ± 3.493 | 0 ± 0 | 0 ± 0 |
| 73-RSS 055 | WA | 2020 | 10 | 292.52 ± 11.155 | 0 ± 0 | 0 ± 0 |
| 73-RSS 056 | WA | 2020 | 10 | 260.76 ± 22.637 | 0.03 ± 0.034 | 0 ± 0 |
| 73-RSS 057 | WA | 2020 | 10 | 233.3 ± 3.88 | 0 ± 0 | 0 ± 0 |
| 73-RSS 058 | WA | 2020 | 10 | 313.81 ± 17.841 | 0.06 ± 0.061 | 0 ± 0 |
| 73-RSS 059 | WA | 2020 | 10 | 202.51 ± 21.21 | 0 ± 0 | 0 ± 0 |
| 73-RSS 060 | WA | 2020 | 10 | 226.43 ± 14.051 | 0 ± 0 | 0 ± 0 |
| 73-RSS 061 | WA | 2020 | 10 | 253.41 ± 3.399 | 0 ± 0 | 0 ± 0 |
| 73-RSS 062 | WA | 2020 | 10 | 202.95 ± 0.685 | 0.1 ± 0.097 | 0 ± 0 |
| 73-RSS 063 | WA | 2020 | 10 | 225.65 ± 18.829 | 0 ± 0 | 0 ± 0 |
| 73-RSS 064 | WA | 2020 | 10 | 205.17 ± 15.718 | 0.07 ± 0.067 | 0 ± 0 |
| 73-RSS 065 | WA | 2020 | 10 | 282.42 ± 6.899 | 0.03 ± 0.031 | 0 ± 0 |
| 73-RSS 066 | WA | 2020 | 10 | 174.1 ± 3.628 | 0 ± 0 | 0 ± 0 |
| 73-RSS 067 | WA | 2020 | 10 | 630.61 ± 24.165 | 0 ± 0 | 0 ± 0 |
| 73-RSS 068 | WA | 2020 | 10 | 677.94 ± 55.083 | 0 ± 0 | 0 ± 0 |
| 73-RSS 069 | WA | 2020 | 10 | 342.73 ± 24.569 | 0 ± 0 | 0 ± 0 |
| 73-RSS 070 | WA | 2020 | 10 | 136.68 ± 3.151 | 0 ± 0 | 0 ± 0 |
| 73-RSS 071 | WA | 2020 | 10 | 232.96 ± 15.249 | 0 ± 0 | 0 ± 0 |
| 73-RSS 072 | WA | 2020 | 10 | 335.22 ± 88.93 | 0 ± 0 | 0 ± 0 |
| 73-RSS 073 | WA | 2020 | 10 | 250.16 ± 22.734 | 0 ± 0 | 0 ± 0 |

^a^ A dash (—) indicates no dilution was performed. A ‘10’ indicates a ten-fold dilution was performed.

^b^ ‘ND’ indicates no fluorescence was detected.
